# A versatile method to expand and compare transporter substrate spectra

**DOI:** 10.1101/2025.07.11.664331

**Authors:** Víctor de Prado Parralejo, Christos Theodorou, Christoph Crocoll, Hussam H. Nour-Eldin

## Abstract

Membrane transport proteins (transporters) are critical players in the physiology of complex organisms. The range of metabolites a transporter translocates across a membrane (substrate spectra) determines the membrane selectivity and, consequently, the transporter’s physiologic function. Nevertheless, determining the full substrate spectra of a transporter, and hence its role(s) in the organism, remains a daunting challenge. Many transporters lack an experimentally determined substrate, and many show a puzzling substrate promiscuity. Substrate promiscuity raises questions about which experimentally defined substrates are physiologic, which are not, and which are missing. Targeted approaches to determine transporter substrate spectra present limitations: Compound availability, lack of unexpected cofactors, and the researchers’ biases. To tackle these issues, transporters can be screened with complex native metabolites to generate new hypotheses on their function. Here, we describe an untargeted method to expand and compare the substrate spectra of sets of plant transporters. This method compares the substrate spectra of different transporters when expressed in *Xenopus laevis* oocytes and exposed to plant metabolic extracts. Comparing the transport-mediated accumulation of metabolites enables us to contextualize the relative transporter activity towards the same set of metabolites from undefined mixtures. Additionally, metabolite accumulation patterns across transporter sets allow us to annotate novel compounds. We validate the method with three well-characterized transporters, where we find unreported substrate spectra specificities and detect the transport of tens of metabolites at a time. We screen a set of fourteen transporters and find unreported activities for eight of them, some of which are confirmed from pure compounds. Finally, we propose to reinterpret the role of two plant transporters previously linked to developmental regulation (phytohormone transport) in light of their substrate promiscuity.

## Introduction

In plants, more than 5% of the predicted coding sequences code for protein transporters, which are responsible for the translocation of most solutes across membranes (Kell & Oliver, 2014; Schwacke et al., 2003). Transporters partake in a wide range of biological processes, such as nutrient uptake and distribution, localization of defense compounds, and developmental coordination (L.-Q. Chen et al., 2010; Hunziker et al., 2021; Verna et al., 2019). Accordingly, gaining fundamental knowledge of plant transport proteins is a promising strategy for identifying breeding targets for crop improvement (Schroeder et al., 2013). However, many plant transporters lack an empirically determined substrate (Schwacke et al., 2003).

Targeted approaches have generated the most knowledge on plant transporter substrate spectra. Usually, a molecule’s transport prompts the search for the causative transporter(s) (Larsen et al., 2017; Niño-González et al., 2019; Nogia & Pati, 2021). Forward genetic screens succeed in identifying transporters for molecules, although they are labor-intensive (Tsay et al., 1993; Yuan et al., 2014). Functional screenings excel at identifying transporters for primary metabolites and nutrients, and can be adapted for the discovery of other molecules (Frommer et al., 1994; Kanno et al., 2012). Although highly successful, targeted approaches are hypothesis-driven and usually fail to reflect whether the transporter displays activity to molecules other than the hypothesized one. Additionally, plant metabolic diversity and commercial unavailability of metabolites hinder the systematic targeted characterization of transporter substrate spectra (Weng et al., 2012). The combination of multiple targeted approaches reveals widespread substrate promiscuity, which is likely underreported (Corratgé-Faillie & Lacombe, 2017). Reverse genetic approaches represent the complementary route, from transporter to substrate, but they rely on the observation of severe phenotypes and can be hampered by gene redundancy (Cusack et al., 2021). Thus, there is a need for untargeted strategies to deorphanize transporters and reassess their current substrate spectra.

Two decades ago, an alternative approach emerged within the biomedical field for the deorphanization of transporters (Gründemann et al., 2005). In this first example, Crohn’s disease-associated transporter OCTN1 was expressed on HEK293 cells and exposed to human plasma. The untargeted analysis of the cells showed import of stachydrine from plasma, which led to the discovery of the structurally related ergothioneine as a novel high-affinity substrate of OCTN1. Later, OCTN1’s link to Crohn’s disease prompted another screening of OCTN1-expressing cells with inflamed colon metabolites, which succeeded in suggesting spermine as a new substrate (Masuo et al., 2018). The physiological substrates of OCTN1 are still debated today (Pochini et al., 2022). Nevertheless, since then, untargeted metabolomic strategies and the usage of complex metabolic mixtures have succeeded in suggesting new substrates to explain and expand the role of transporters (Abplanalp et al., 2013; Jansen et al., 2015; Krumpochova et al., 2012; Kuang et al., 2021; Møller-Hansen et al., 2024; Radi et al., 2024; van de Wetering & Sapthu, 2012).

To our knowledge, one study has adopted the usage of complex mixtures for characterizing plant transporters (Demurtas et al., 2019). In searching for crocin transporters in *Crocus sativa* (saffron), two ABC and three MATE transporters were tested due to their expression in the crocin-accumulating stigmas. Microsomes derived from transporter-expressing yeast cells were exposed to saffron stigma extracts, followed by LC-MS analysis to identify CsABCC4a and CsABCC2 as crocin transporters.

This study underlined three major benefits of using complex metabolite extracts as substrate mixtures: 1-The extracts contained physiologic crocins forms, some of which were not commercially available 2-The simultaneous presence of two substrates resulted in substrate cooperativity leading to higher transporter uptake of the individual compounds 3-The import of compounds other than crocins demonstrated the functional expression of the MATEs and thus eliminated ambiguity regarding their inability to transport crocins.

Despite these advances, the development of data-driven untargeted approaches for identifying plant transporter substrates has not continued. Common procedures for transporter screening, such as human cell line-or vesicle-based systems, require specialized laboratory facilities or laborious preparation procedures (Demurtas et al., 2019; Wright Muelas et al., 2020). Yeast has the disadvantage of harboring high endogenous transport capabilities that may obscure the detection of transporter activities. Transporter-defective yeast mutants offer lower transport activity backgrounds (Demurtas et al., 2019; J. Kang et al., 2010; Riesmeier et al., 1992), but these are restricted to a subset of molecules, and thus, they are less suitable for untargeted screens.

*Xenopus laevis* oocytes represent a well-established model system for characterizing transporters and channels (Bhatt et al., 2022). Oocytes offer high and functional expression of heterologous membrane proteins and a low and well-described transport activity background (Sobczak et al., 2010; Terhag et al., 2010). Moreover, owing to their large size (about 1–1.2 mm in diameter), they can be injected with substrates and assayed for export activity (Xu et al., 2016) and they have been used in the targeted screenings of large transporter libraries for activity towards natural compounds (Nour-Eldin et al., 2012). Oocytes have already proven useful in the context of untargeted metabolomic substrate searches. For example, expression of human MCT12 in oocytes led to a decrease in endogenous creatine, which deorphanized MCT12 (Abplanalp et al., 2013). In another example, oocytes were used to compare the substrate specificities of multiple amino acid transporters when exposed to the same defined cell culture medium (Fairweather et al., 2021). Lastly, the expression capabilities and versatility of *Xenopus* oocytes allowed the screening of a library of 30 yeast transporters with six xenobiotic compounds. This targeted approach identified additional xenobiotic transporters whose activity might interfere with synthetic biology approaches in yeast (Møller-Hansen et al. 2024).

In this study, we aim to build on these various advances to demonstrate the potential of an untargeted metabolomics approach using complex metabolite extracts from plants and *Xenopus* oocytes as expression hosts to expand and reassess the substrate spectra of transporters.

We exposed transporters from *Arabidopsis thaliana* (*A. thaliana)* belonging to the nitrate and peptide transporter family (NPF) to two different metabolite extracts from *A. thaliana*. We identified diverse glucosinolates, flavonols, monolignol glucosides, and sinapate esters as substrates, expanding the substrate spectrum of known transporters. Additionally, we found unreported differences in substrate spectra of AtNPF2.10 and AtNPF2.11 towards endogenous glucosinolates. Moreover, we portrayed substrate promiscuity of AtNPF2.13 and AtNPF3.1, two transporters linked to phytohormone transport.

## Results

### Phytochemical permeation characterization

Membrane integrity is a basic premise for the development of transport assays. i.e., compounds may not leak across the membrane without the mediation of the transporter. The early untargeted approaches with complex media used diluted metabolic mixtures (Gründemann et al., 2005; Krumpochova et al., 2012). However, extract dilution also reduces the concentration of substrates, hence the possibility of detecting transporter activity. To evaluate the effect of extract dilution on membrane integrity, we exposed non-transporter-expressing oocytes to various concentrations of metabolite extracts and monitored the content of selected plant metabolites within the oocytes. We chose glucosinolates as representative *A. thaliana* metabolites that are readily detectable with Liquid Chromatography Mass Spectrometry, and that do not diffuse across uncompromised oocyte membranes (Crocoll et al., 2016; Jørgensen et al., 2015).

We generated two different extracts. One from seeds and one from 2-day-old seedlings. These tissues were chosen due to the metabolic changes that germination elicits in the plant (Kliebenstein et al., 2007; Routaboul et al., 2006; Shirley & Chapple, 2003), and the ease of growing seedlings in axenic conditions, which removes metabolites derived from microorganisms. Additionally, transporter expression and repression can correlate with transporter substrate presence, and since germination alters the expression of genes in this work, it could increase the possibility of extracting relevant substrates (Demurtas et al., 2019; Narsai et al., 2011; Payne et al., 2017). For both tissues, we resuspended extraction pellets corresponding to 100 mg of tissue dry weight in 1 ml of oocyte buffer (1:1), which we diluted two-(1:2) or fivefold (1:5). We monitored the presence of six phytochemicals (glucosinolates) as membrane integrity indicators.

All the oocytes assayed with the different dilutions of seed and seedling extracts presented signal for the most readily detectable compound, glucoerucin/ 4-methylthiobutyl glucosinolate (4mtb) (Fig. 1), as well as other selected glucosinolates (Sup. 1). There was a trend of decreasing oocyte glucosinolate levels with extract dilution for both extracts. The oocytes exposed to the undiluted seed extract accumulated significantly more glucosinolates than the oocytes exposed to the undiluted seedling extract. This difference was likely due to the higher glucosinolate content in the seed extract (Sup. 2). Although not statistically significant, 4mtb levels in oocytes assayed with seedling media lowered with a twofold dilution of the media, and the levels did not lower noticeably upon fivefold dilution. On the other hand, there was a statistical difference between oocytes assayed with the undiluted seed extract and the twofold diluted version, and there was a non-significant difference between oocytes assayed with the twofold and fivefold dilutions of seed extracts. Additionally, there was a high variance in glucosinolate levels in oocytes assayed with any seed extract dilution, which was also observable for the highest concentration of seedling extract. This variance suggested that oocyte membrane integrity was affected by both the extracts and the fitness of the individual oocytes. By plotting phytochemical content within control oocytes as a function of extract dilution factor, we inferred that the indicated dilution of extracts depended on the origin of plant material. In our case, we suggested using a seedling extract diluted more than twofold, and a seed extract diluted more than fivefold. We showed that non-transporter-mediated phytochemical accumulation in oocytes was dilution-dependent, and membrane permeation could be alleviated with extract dilution.

**Figure 1.**
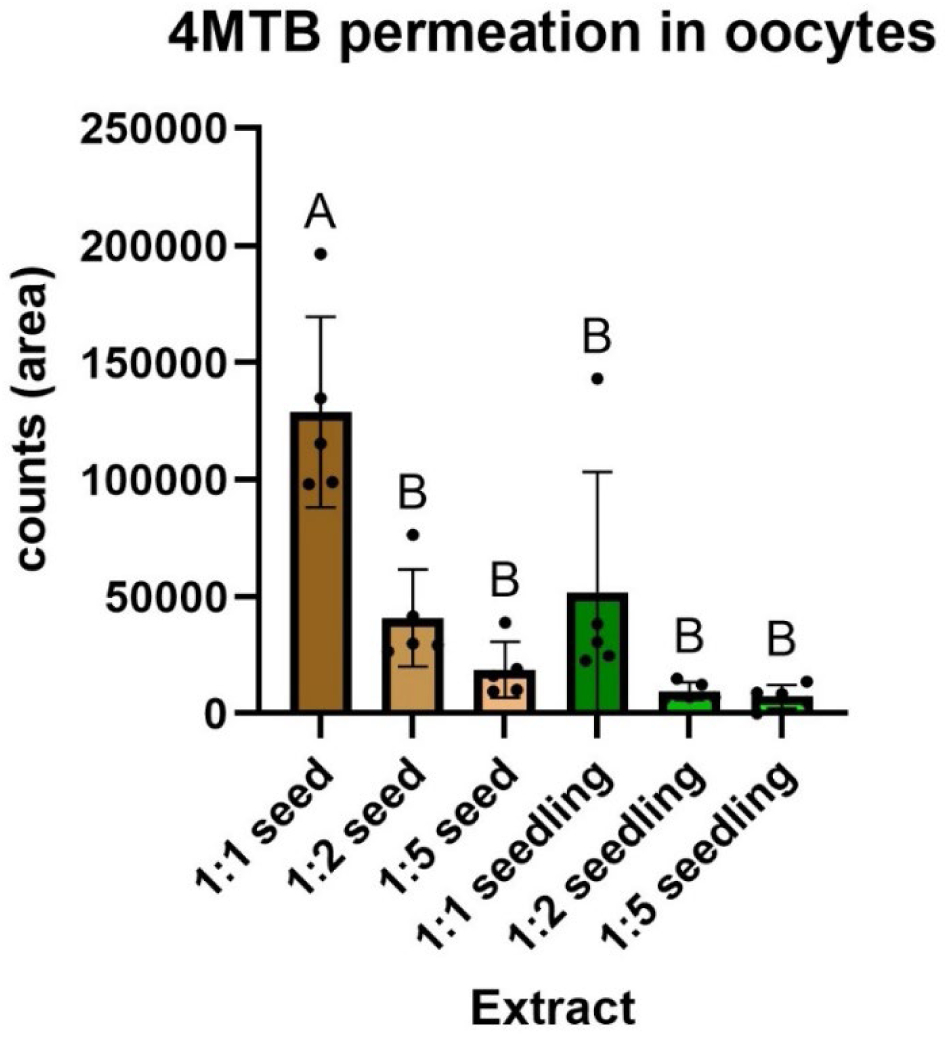
Extract dilution alleviates membrane permeation. Targeted detection of the plant-derived glucoerucin (4-methylsulfinylbutyl glucosinolate, 4mtb) in control oocytes exposed to different dilutions of seed and seedling extracts. Oocytes were assayed at pH 5 for 1 hour; replicates consisted of three oocytes each. Bars show the averages of 5 pools of 3 oocytes (n = 5) measured with multiple reaction monitoring (MRM), and error bars represent the standard deviation. Letters represent the statistically different groups for a Tukey Multiple comparison test preceded by One-way ANOVA, Sig. P-value <0.05.

### Method validation by untargeted screening of the GTRs

As a proof-of-concept, we tested the method by exposing three glucosinolate transporters, NPF2.11, NPF2.10, and NPF2.9 (GTRs1-3, respectively) (Jørgensen et al., 2017; Nour-Eldin et al., 2012) to the metabolite extracts. *A. thaliana* NPF2.9 shows a preference for tryptophan-derived glucosinolates (indole-glucosinolates) (Jørgensen et al., 2017). Recently, NPF2.10 and NPF2.11 showed differences in substrate preference when exposed to artificial fluorophore-tagged glucosinolates, suggesting subtle hitherto unreported differences in substrate preference between NPF2.10 and NPF2.11 (Kanstrup, Jimidar, et al., 2023). Consequently, we asked whether the GTRs would be able to import glucosinolates from the complex metabolite plant extract, whether NPF2.9 would display its preference for indole-derived glucosinolates, and whether we could reveal differences in substrate preference between NPF2.10 and 2.11 towards endogenous substrates. Also, we wanted to test how the substrate profiles of the GTR-expressing oocytes would change according to the metabolic source. For such purpose, we exposed GTR-expressing oocytes to a 1:100 dilution of either the seed or seedling extract to ensure that permeation of phytochemicals in oocytes from the two extracts would be alleviated.

First, we compared glucosinolate accumulation in GTR-expressing oocytes according to extract for the major indole-glucosinolates glucobrassicin (I3M), 4-methoxyglucobrassicin (4MOI3M), and neoglucobrassicin (NMOI3M) and the major aliphatic glucosinolates glucoerucin (4mtb), glucoraphanin (4msb), 8-methylthiooctyl glucosinolate (8mto), and glucohirsutin (8mso). All GTRs exposed to seedling extracts accumulated higher signals of the selected glucosinolates compared to control oocytes (Fig. 2 and Fig. 3), whereas for the seed extract, the GTRs only accumulated noticeable signals for the aliphatic glucosinolates and I3M, reflecting the medium’s low signal for 4MOI3M and NMOI3M (Sup. 3 and Sup. 4), The control oocytes did not accumulate comparable levels to the transporter-expressing oocytes of any of the glucosinolates. All GTRs accumulated the three indole-glucosinolates above media levels from the seedling extract (Fig. 2), and I3M from the seed extract (Sup. 3). NPF2.9 accumulated higher levels of indole-glucosinolates than NPF2.10 or NPF2.11 when exposed to either extract.

**Figure 2.**
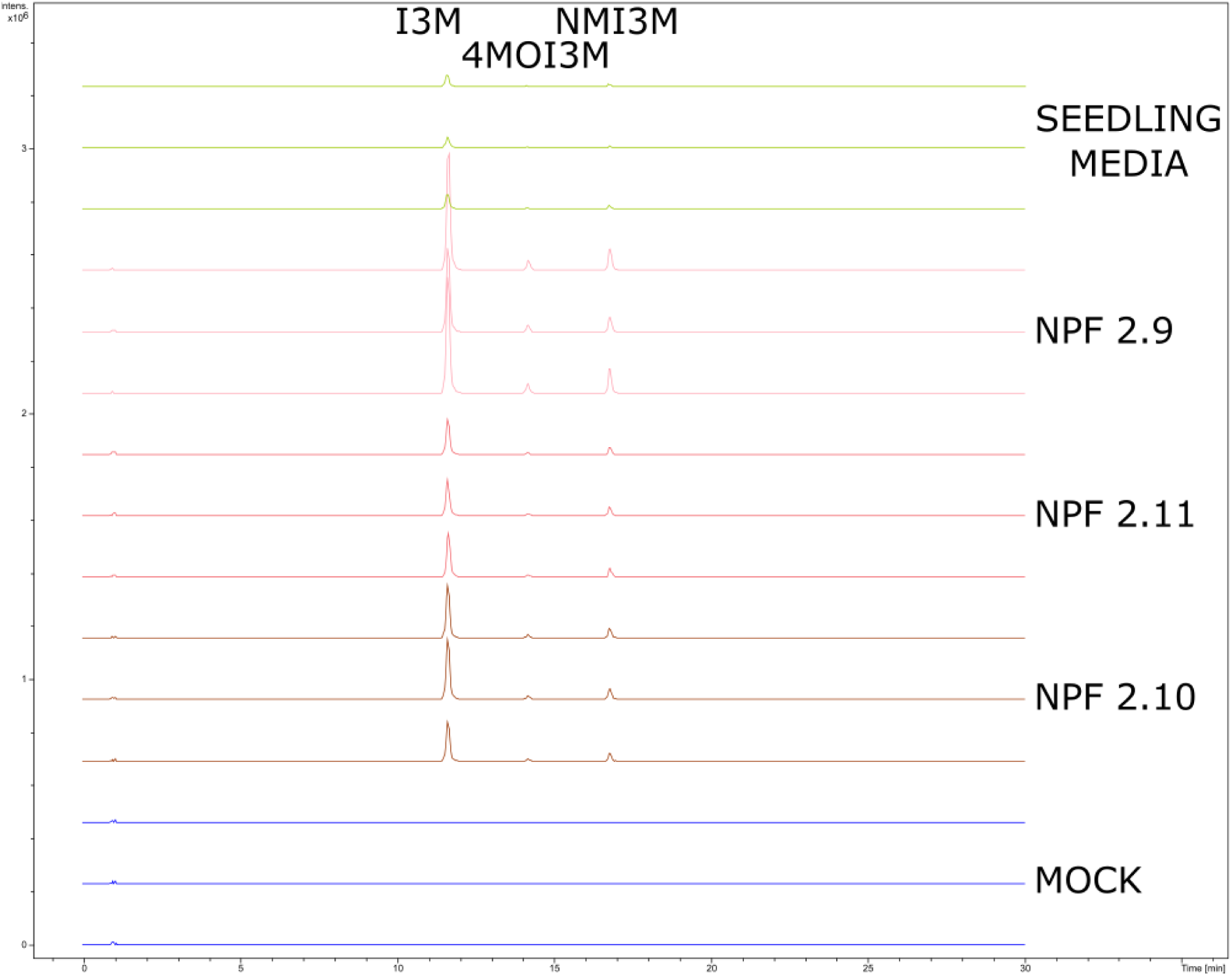
NPF2.9-expressing oocytes accumulate higher levels of indole-glucosinolates than NPF2.10 and NPF2.11-expressing oocytes. Presence of indole-glucosinolates glucobrassicin (I3M), 4-methoxyglucobrassicin (4MOI3M), and neoglucobrassicin (NMOI3M) in oocytes exposed to 1:100 seedling media. NPF2.10 in brown, NPF2.11 in red, and NPF2.9 in pink. Oocytes were assayed at pH 5 for 1 hour; replicates consisted of five oocytes each, and media samples consisted of 5µl each. Control oocytes in blue and seedling media in green. Combined extracted ion chromatogram for I3M (447.0537 +/-0.005 m/z) and 4NMOI3M and NMOI3M (477.0643 +/-0.005 m/z).

**Fig 3.**
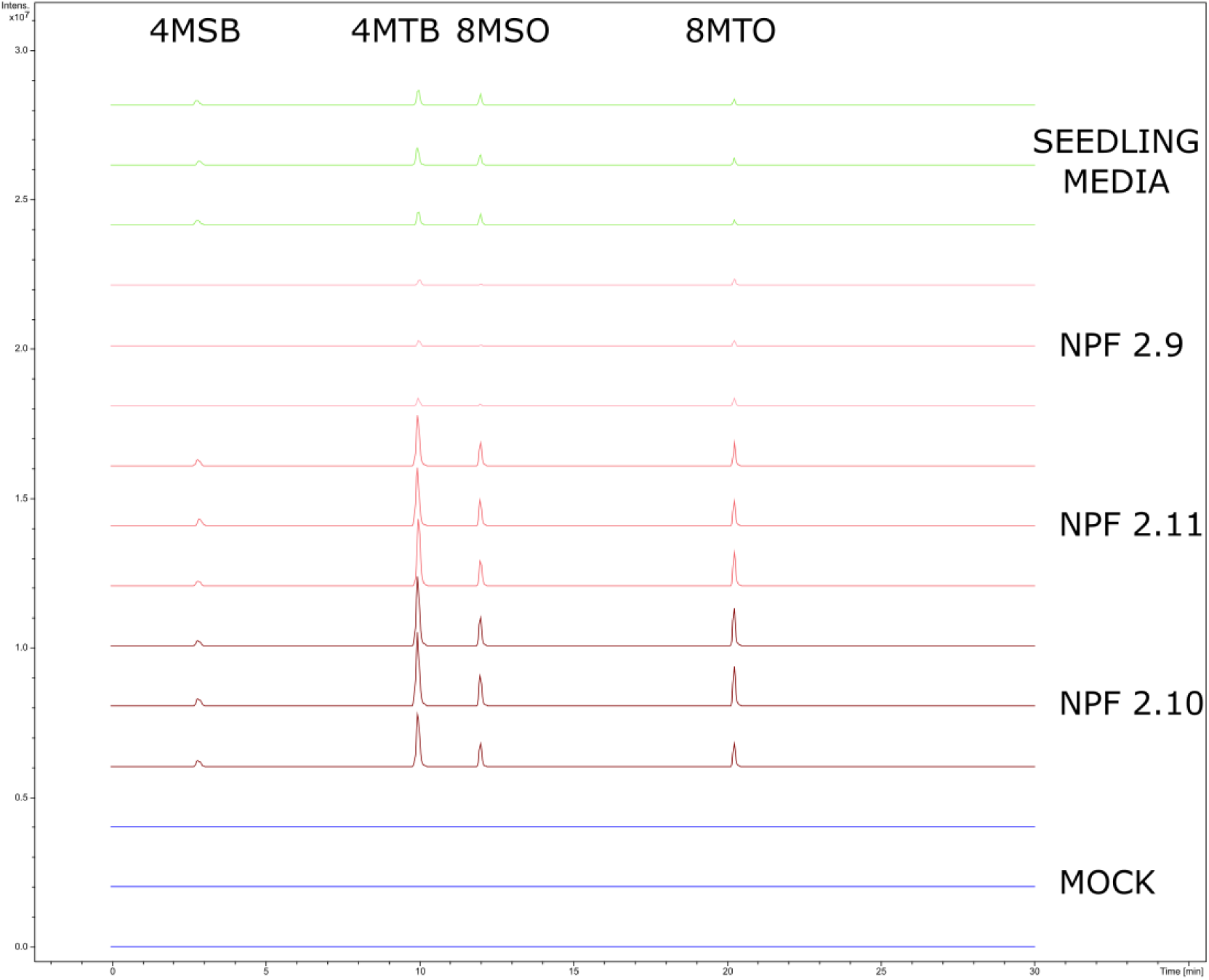
NPF2.9-expressing oocytes cannot accumulate aliphatic glucosinolates to NPF2.10 and NPF2.11 levels. Presence of aliphatic glucosinolates glucoerucin (4mtb), glucoraphanin (4msb), 8-methylthiooctyl glucosinolate (8mto), glucohirsutin (8mso) in oocytes exposed to 1:100 seedling media. NPF2.10 in brown, NPF2.11 in red, and NPF2.9 in pink. Oocytes assayed at pH 5 for 1 hour; replicates consisted of five oocytes each, and media samples consisted of 5µl each. Control oocytes in blue and seedling media in green. Combined extracted ion chromatogram for 4msb (436.0411 +/-0.005 m/z), 4mtb (420.0462 +/-0.005 m/z), 8mso (492.1037 +/-0.005 m/z) and 8mto (476.1088 +/-0.005 m/z).

The higher accumulation of I3M in NPF2.9-expressing oocytes compared to 2.10 is only reported when other non-indole glucosinolates compete with I3M (Jørgensen et al., 2017; Kanstrup, Wulff, et al., 2023), suggesting that in our case, the multiple glucosinolates or other compounds from the media were competing with the import of indole-glucosinolates in NPF2.10 and NPF2.11-expressing oocytes. The competition of substrates when using complex mixtures has been reported before (Krumpochova et al., 2012)

On the other hand, NPF2.10 and NPF2.11 accumulated the major aliphatic glucosinolates over NPF2.9 from both extracts (Fig. 3, Sup. 4). NPF2.9 displayed a higher accumulation of 4mtb and 8mto than 4msb and 8mso relative to media levels, particularly for the seed extract. Oocytes display a total volume between 0.52 and 0.9 µl, but the water-accessible volume is calculated to be as low as 0.36 µl, thus, the media samples likely represented a higher volume than the oocyte samples (Stegen et al., 2000). Thus, it was challenging to assess if levels of 4mtb and 8mto in NPF2.9-expressing oocytes were above media levels without additional measurements, which is out of the scope of this work. Altogether, the data supported the published substrate spectra regarding the indole-glucosinolate preference of NPF2.9.

Next, we performed an untargeted analysis of the data from GTR-expressing oocytes exposed to both media. Principal Component Analysis (PCA) displayed a separation of the control oocyte chemical profiles and the GTR-expressing oocytes (Sup. 5). The first two components of the PCA accounted for more than 60% of the data variability, and thus the analysis was informative. The first component separated the oocyte groups (40% of variability), while the second separated samples within groups (20% of variability), indicating that there was a noticeable level of chemical variability within groups. All control oocytes assayed with non-phytochemical-containing oocyte media (pH 5) clustered together, but Principal Component 2 spread the samples of control oocytes assayed with either extract media. This suggested variable oocyte background levels of transport, adsorption, or permeation of phytochemicals. NPF2.9 oocytes separated distinctly from other oocytes, and also according to the extract used. NPF2.10 and NPF2.11 groups did not cluster separately from each other. Nevertheless, the PCA only represents global chemical profiles. This prompted us to represent the chemical profile of the oocytes with a heatmap as a tool to compare global substrate spectra specificities at feature accumulation level. We represented the 100 statistically most significant features of the untargeted analysis in a heatmap and biclustered features and samples in an unsupervised manner (Fig. 4).

**Fig 4.**
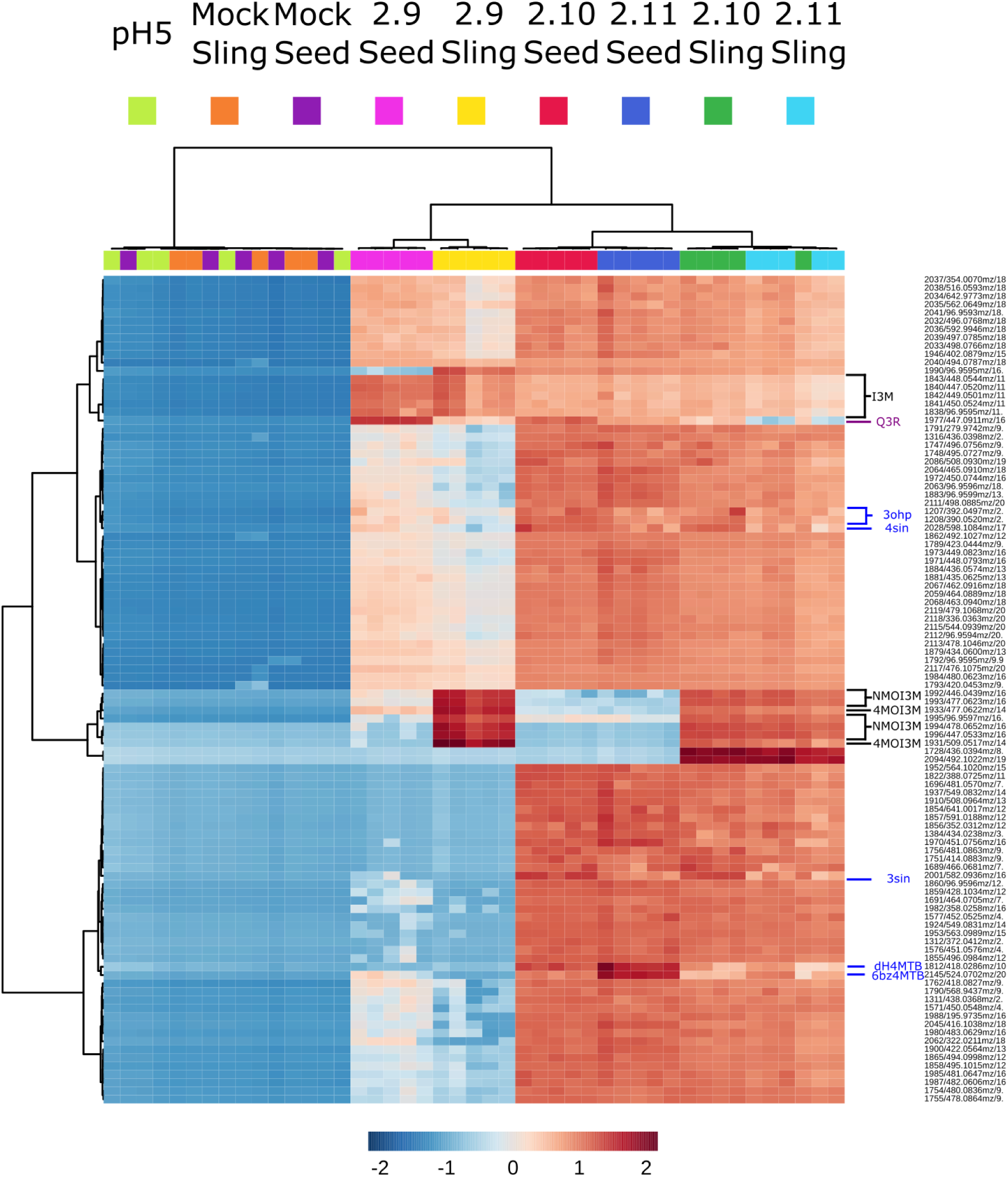
**The metabolic differences of the extracts and transporter activity drive the chemical profile of the oocytes**. Unsupervised bicluster analysis of oocytes assayed with empty buffer (pH5), and control (mock) and GTR-expressing oocyte samples assayed with either seed (Seed) or seedling (Sling) derived media. The relative intensities of metabolic features are represented on a color scale. Columns represent samples, and rows represent metabolic features. Oocytes were assayed at pH 5 for 1 hour with either 1:100 dilutions of seed or seedling extract; replicates consisted of five oocytes each. pH5 oocytes were assayed with an empty pH 5 buffer for 1 hour. Metabolic features identified with chemical standards as indole glucosinolates in black font: glucobrassicin (I3M), neoglucobrassicin (NMOI3M), and 4-methoxyglucobrassicin (4MOI3M). Annotated glucosinolates in blue font: 3-sinapoyloxypropylglucosinolate (3sin), 4-sinapoyloxybutylglucosinolate (4sin), glucoraphasatin (dH4mtb), 6′-O-benzoyloxy-glucoerucin (6’bz4mtb). Annotated flavonoid Quercitrin (Q3R) in purple. Ward’s biclustering of the 100 most significant features (ANOVA). Sig. P-value <0.05. Features displaying a Relative Standard deviation of >30% in the QC were filtered. Features are logarithmically transformed (Log10) and Pareto scaled. Identification of indole glucosinolate-derived features was performed with co-analyzed chemical standards.

The unsupervised biclustering separated the transporter-expressing oocyte samples into clear clusters according to the extract and the transporter. Control samples did not separate according to extract, which clustered together with oocytes assayed with empty media. The successful separation of transporter-expressing oocytes according to extract suggested that extracts conferred the transporter-expressing oocyte groups with unique metabolic profiles. All represented features accumulated in transporter-expressing oocytes relative to control oocytes, which indicated that these features were attributable to import from the media. NPF2.9 samples clustered together regardless of the extract used. On the other hand, NPF2.10 and 2.11 clustered together when assayed with the same extract. This suggested a higher similarity in oocytes expressing NPF 2.10 and 2.11 than in oocytes expressing 2.9. Feature clustering separated metabolites according to the relative activities the transporters display, such as the two clusters with indole-glucosinolate features, which contributed to the separation of NPF2.9 from NPF2.10 and 2.11 and displayed higher intensities for NPF2.9. One cluster predominantly contained I3M-derived features, which present comparable intensities in NPF2.9-expressing transporters exposed to seedling and seed extracts. The other cluster contained features derived from NMOI3M and 4MOI3M, since these metabolites were noticeably lower in the seed extract (Sup. 4) and therefore accumulated to lower intensities in all GTR-expressing oocytes assayed with seed extract. When assayed with the same media, most NPF2.10 samples clustered separately from NPF2.11 samples since specific features accumulated differentially according to the gene.

We annotated two of these features that separated the profiles of NPF2.10 and 2.11 as 3-sinapoyloxypropylglucosinolate (3sin) and 4-sinapoyloxybutylglucosinolate (4sin). Both 3sin and 4sin present a sinapoyl moiety attached to the end of the glucosinolate aliphatic chain. They are the esterification products of sinapic acid and 3-hydroxypropyl or 4-hydroxybutyl glucosinolate, respectively. They are present in *A. thaliana* seeds, and the GTRs also imported them from the seedling media (Fig. 4) (Kliebenstein et al., 2007; Lee et al., 2012). Both 3sin and 4sin were detectable in seed media, although close to detection levels (Fig. 5). NPF2.10 accumulated 3sin and 4sin over 2.11, where NPF2.10 accumulated both over media levels, and NPF2.11 only accumulated 4sin over media levels. NPF2.9 displayed the lowest accumulation for both.

**Fig 5.**
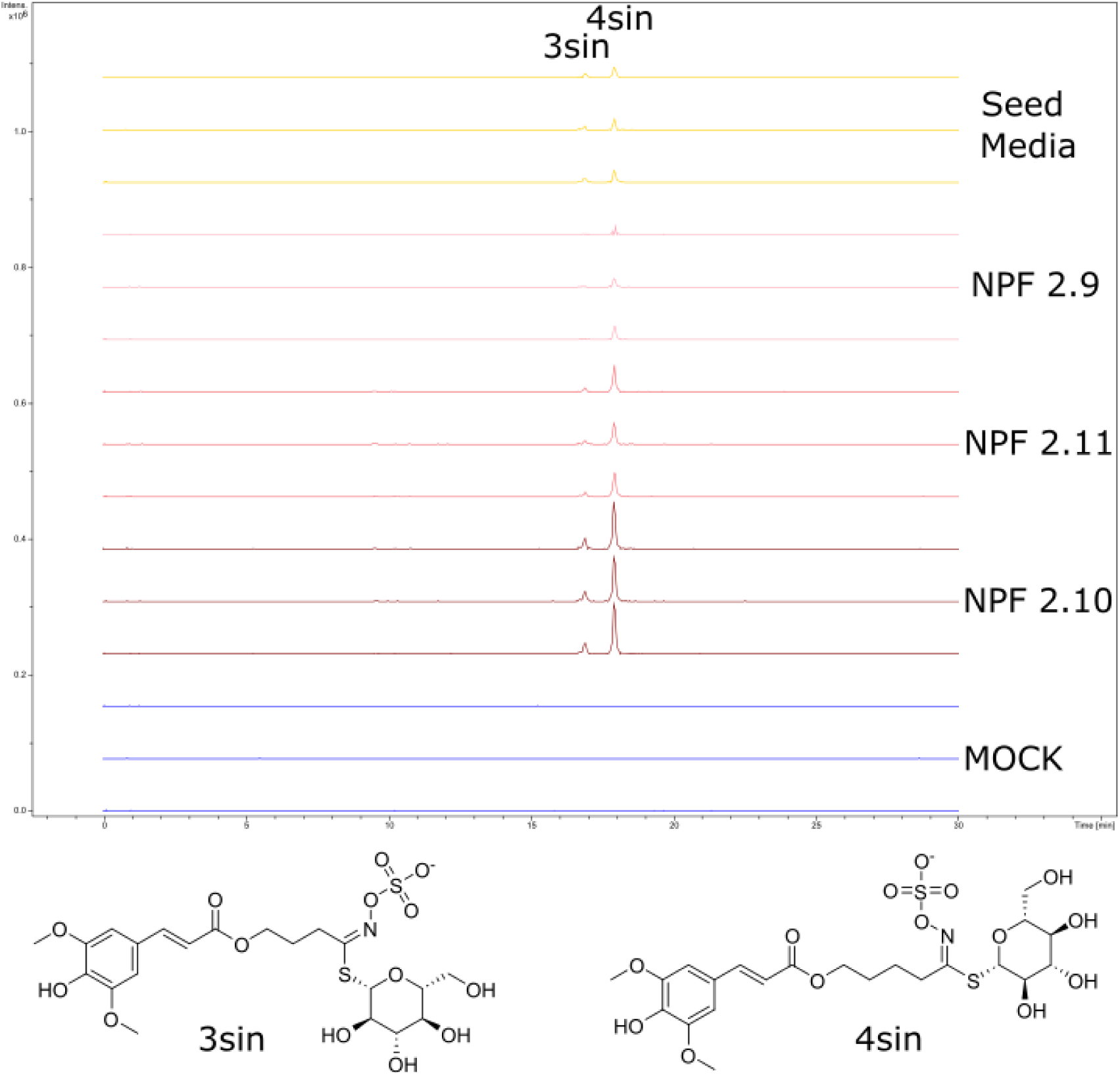
NPF2.10 accumulates the annotated sinapoylated glucosinolates over NPF2.11-expressing oocytes. Presence of the annotated 3-sinapoyloxypropyl-(3sin) and 4-sinapoyloxybutyl-(4sin) glucosinolates in oocytes exposed to 1:100 seed media. NPF2.10 in brown, NPF2.11 in red, and NPF2.9 in pink. Oocytes were assayed at pH 5 for 1 hour; replicates consisted of five oocytes each, and media samples consisted of 5µl each. Control oocytes in blue and seed media in yellow. Combined extracted ion chromatogram for 3sin (582.0957 +/-0.005 m/z) and 4sin (596.1113 +/-0.005 m/z). The structures of the metabolites differ in the length of the aliphatic chain.

On the other hand, we annotated two glucosinolates preferentially imported by NPF2.11 as 6’-O-benzoyloxy-glucoerucin (6’bz4mtb), and glucoraphasatin/dehydro4mtb (dH4mtb). 6’bz4mtb is the esterification product of benzoic acid and the sugar moiety of 4mtb and is present in *A. thaliana* seeds (Reichelt et al., 2002). dH4mtb is present in *Raphanus sativus* and not described in *A. thaliana* (J.-N. Kang et al., 2020). 6’bz4mtb was present in the seed media, although close to the detection level (Sup. 6). NPF2.11 accumulated 6’bz4mtb over 2.10, which in turn accumulated over NPF2.9, all three over media levels. NPF2.11 accumulated dH4mtb slightly over 2.10, whereas for NPF2.9, there was barely detection. dH4mtb was hardly detectable in the media Additionally, we annotated the flavonoid quercitrin (Q3R) as a non-glucosinolate substrate of the GTR’s (Fig. 4, Sup. 7), which accumulates in *A. thaliana* seeds (Routaboul et al., 2006). Q3R appeared in the biclustering as a feature that accumulated preferentially in oocytes assayed with seed over seedling extract when comparing within the same transporter, and accumulated preferentially in NPF2.9, then in NPF2.10, and to a lower extent in NPF2.11 when comparing oocytes assayed with the same extract. This feature did not follow a GTR-accumulation pattern attributable to any of the glucosinolates previously described in the heatmap, likely due to the different chemical class of this substrate.

The data analysis of this untargeted approach reflected the major known substrate specificities of the GTRs regarding indole-glucosinolates and suggested novel differential specificities. The biclustering represented the global NPF2.9 preference towards the indole-glucosinolate for two different extracts. Accordingly, transporter accumulation of metabolites clustered the features according to the functional groups of their substrates, such as the clustering of the indole glucosinolates or sinapoylated glucosinolates. This was very useful for feature annotation and hypothesizing on structural determinants for substrate recognition. When comparing multiple transporters, features described accumulation patterns that gave hints into chemical similarity, like Q3R versus glucosinolates. These accumulation patterns were conserved when using two mixtures as different as seed and seedling extracts. Some differential features of NPF2.10 versus 2.11 are known *A. thaliana* metabolites and thus constitute novel differences in substrate specificity for endogenous plant compounds.

### Untargeted screening of 14 NPFs

Next, we tested the method with additional transporters. The media signal for some of the previous metabolites was close to detection levels (Fig. 5, Sup. 6), which hindered assessing the media presence of these metabolites, a prerequisite for claiming metabolite transporter-mediated translocation. We also wanted to increase the output of the method by increasing the concentration of metabolites and, thus, potentially, the detection of import. We assayed a 1:20 dilution of the seedling extract with an additional eleven other NPF transporters induced or repressed upon germination (Narsai et al., 2011). The transporters are listed in Table 1.

**Table 1.** NPF transporters included, and some of their described substrate spectra.

| Transporter | Substrate | References |
| --- | --- | --- |
| AtNPF1.1 | Nitrate | (Hsu & Tsay, 2013) |
| AtNPF2.5 | Chloride, Gibberellins | (B. Li et al., 2017; Wulff et al., 2019) |
| AtNPF2.9 | Glucosinolates and Nitrate | (Jørgensen et al., 2017; Y.-Y. Wang & Tsay, 2011) |
| AtNPF2.10 | Glucosinolates and jasmonoyl-isoleucine | (Nour-Eldin et al., 2012; Saito et al., 2015) |
| AtNPF2.11 | Glucosinolates | (Nour-Eldin et al., 2012) |
| AtNPF2.13 | Nitrate, Gibberellins, Absciscic acid, and tunicamycin | (Binenbaum et al., 2023; Fan et al., 2009; Liu et al., 2023) |
| AtNPF3.1 | Gibberellins | (Tal et al., 2016) |
| AtNPF4.1 | Gibberellins, Absciscic acid, and jasmonoyl-isoleucine | (Chiba et al., 2015) |
| AtNPF4.6 | Gibberellins, Absciscic acid, and jasmonoyl-isoleucine | (Chiba et al., 2015) |
| AtNPF5.2 | Peptides | (Karim et al., 2007) |
| AtNPF6.2 | Nitrate | (Chiu et al., 2004) |
| AtNPF6.3 | Nitrate | (Tsay et al., 1993) |
| AtNPF7.2 | Nitrate | (J.-Y. Li et al., 2010) |
| AtNPF8.1 | Peptides | (Dietrich et al., 2004) |

First, we performed pairwise comparisons of the chemical features in the different transporter-expressing oocytes with the control group. The significant features were visually qualified according to their presence in media samples as a condition for transporter-mediated translocation, and not the effects of transporter expression or function on the oocytes’ metabolism. Some transporters accumulated low amounts of media metabolites, but noticeably lower than media levels. Coincidentally, these metabolites displayed high signals in the media and variable accumulation within the oocyte sample group. This heterogeneous accumulation contrasted with the previously validated homogeneous accumulation of glucosinolates. Consequently, they were disregarded as stemming from permeation/negligible amounts of transport (Sup. 8). This process validated hits for 8 of the 14 transporters (Sup. Tables 1-8).

Validated features accumulated in transporters from NPF clades 1, 2, and 3. We annotated glucosinolates and phenylpropanoids like sinapate esters, flavonols, or monolignol glucosides. Most glucosinolates accumulated in the GTRs (Sup. Tables 3, 4, and 5) but also to lesser extents in NPF2.13 and 3.1 (Sup. Tables 6 and 7). When examined, we observed that a benzoylated glucosinolate, 6’Obz4mtb, also accumulated in NPF1.1 (Sup. 9). In this experiment, NPF2.11 accumulated 6’Obz4mtb higher than NPF2.10, as previously observed (Sup. 6). NPF1.1 accumulated 6’Obz4mtb in comparable levels to NPF2.13 and 2.11, while NPF3.1 and 2.10 accumulated lower comparable levels. With the increased concentration of the seedling extract, the oocytes accumulated additional detectable features. For example, a feature with a m/z of 598.1058 accumulated across oocytes in the same pattern as 6’Obz4mtb, and barely over the detection levels (Sup. 10). We annotated this as 6′-O-benzoyloxy-4-benzoyloxybutylglucosinolate (6’Obz4bzo), a metabolite present in seeds (Reichelt et al., 2002). These two annotated benzoylated glucosinolates shared a similar accumulation pattern across the wider transporter set, where they accumulate in NPF1.1 to comparable levels to NPF2.11 and 2.13, and to lower levels in NPF2.10 and 3.1. In comparison, the previous experiment with the hundredfold diluted version of the extract GTRs presented non-detectable levels of 6’Obz4bzo (Sup. 11). Additionally, NPF1.1 also accumulated sinapate esters (Sup. Table 1).

In addition to these two annotated benzoylated glucosinolates, we observed a feature with a mass of 700.1370m/z accumulating in NPF1.1, 2.13, and 3.1 (Sup.12). According to the modularity of *A. thalian*a’s metabolome, and the capacity of these transporters to transport benzoylated glucosinolates, we hypothesized this metabolite to be the benzoylated version of 4sin, with a molecular formula of C_29_H_35_N_1_O_15_S_2_, and a predicted [M-H]-ion of 700.1375 matching the measured ion. The MS2 ionization spectra of the precursor ion displayed an ion of 96.95 m/z, which corresponded to the sulfate moiety typical of glucosinolate fragmentation, and two additional fragments of 112.9 and 116.9 (Sup. 13). Accordingly, we tentatively annotated this feature as 6′-O-benzoyloxy-4-sinapoyloxybutylglucosinolate (6’bz4sin).

Similarly, we annotated another feature imported by NPF1.1, NPF2.13, and NPF3.1, displaying the same accumulation pattern, but an earlier distinct elution time (Sup. 14), consistent with a tentatively annotated 6′-O-benzoyloxy-3-sinapoyloxypropylglucosinolate (6’bz3sin). The measured m/z of 686.1215 matched the predicted molecular ion of 686.1219 for C_28_H_33_N_1_O_15_S_2._ This precursor ion also displayed similar MS2 spectra with fragments of 96.95, 112.9, and 116.9, which reinforced our hypothesis on the homology of these structures (Sup. 15).

The accumulation of (6’Obz4bzo) in NPF1.1-expressing oocytes enabled the annotation of the two hypothetical structures (6’bz3sin and 6’bz4sin). NPF2.11 only accumulated comparable levels of 6’Obz4bzo to NPF1.1, 2.13, and 3.1 (Fig. 6). These three transporters accumulated the two hypothetical structures to levels comparable media, where NPF2.13 accumulated the highest levels. NPF2.13 and NPF3.1 (Sup. Tables 6 & 7) displayed the greatest promiscuity since they accumulated various sinapate esters, flavonols, and other glucosinolate classes. NPF2.13 and NPF3.1 did not accumulate higher levels of representative aliphatic or indole-glucosinolates over the GTRs (Sup. 16 and 17). We annotated 3-benzoyloxypropyl and 4-benzoyloxybutyl glucosinolates (3bzo and 4bzo), which are characteristic of the seeds and siliques of *A. thaliana* (Graser et al., 2001). NPF2.13 and 3.1 did not accumulate the benzoylated 3bzo and 4bzo over the GTRs (Sup. 18). Thus, it appeared that the benzoylation of the aliphatic chain of the glucosinolate did not elicit the same substrate recognition as the benzoylation of the sugar for these two transporters.

**Figure 6.**
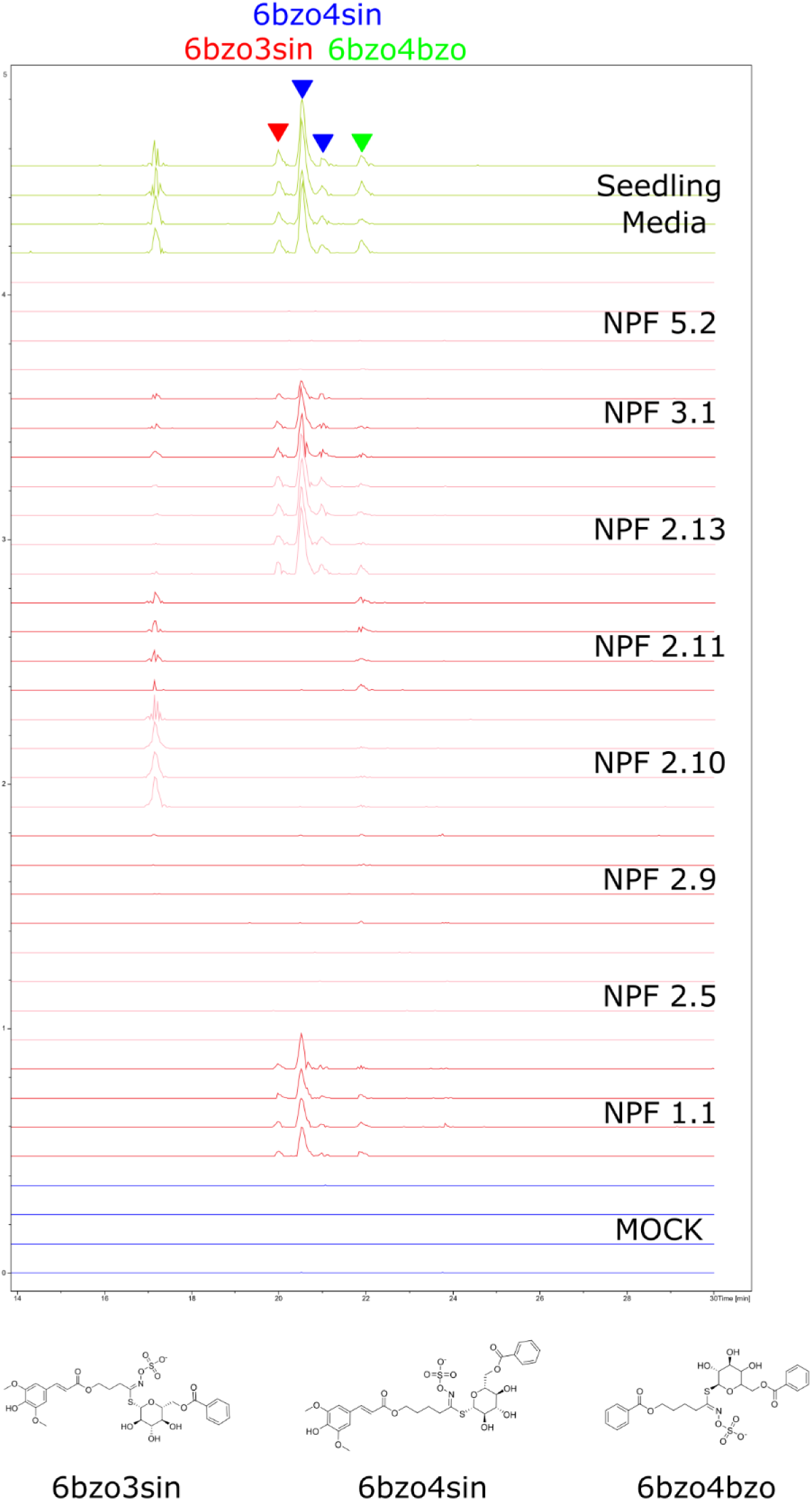
The up-concentration of the extract and leveraging the accumulation patterns for the annotation of substrates allow the expansion of the NPF substrate spectra towards hypothetical benzoylated glucosinolates. Presence of benzoylated glucosinolates 6′-O-benzoyloxy-4-benzoyloxybutylglucosinolate (6’bz4bzo), 6′-O-benzoyloxy-3-sinapoyloxypropylglucosinolate (6’bz3sin), and 6′-O-benzoyloxy-4-sinapoyloxybutylglucosinolate (6’bz4sin), in oocytes exposed to 1:20 seedling media. Oocytes were assayed at pH 5 for 1 hour; replicates consisted of five oocytes each, and media samples consisted of 5µl each. Control oocytes in blue and seedling media in green. Combined extracted ion chromatogram for 6’bz4bzo (598.1058 +/-0.005 m/z), 6’bz3sin (686.1219+/-0.005 m/z), and 6’bz4sin (700.1219 +/-0.005 m/z).

On the other hand, we did not annotate high glucosinolate import relative to the media from NPF2.5. Instead, NPF2.5 accumulated media levels of monolignol glucosides and some levels of flavonols (Sup. Table 2). The identity of the monolignol coniferyl alcohol glycoside (coniferin) was confirmed with a chemical standard (Sup. 19). This prompted us to examine the presence of the other major monolignol glucoside present in *A. thaliana,* syringin (Chu et al., 2014), whose level was close to the detection limit, and yet also accumulated in NPF2.5 (Sup. 20).

The flavonol quercitrin (Q3R) accumulated in negligible extents compared to media levels for most of the 14 NPF transporters we assayed (Sup. 21), except for NPF2.13, which accumulated comparably higher levels for three out of the four flavonols that we confirmed with chemical standards (Sup. 21 and 22). NPF2.13, 3.1, and 2.5 accumulated afzelin (K3R) and isoquercetin (Q3G) over the rest of the transporters (Sup. 22). The levels of accumulation of Q3R and astragalin (K3G) had to be discriminated from the in-source fragments corresponding to their aglycon since the molecular ions have the same mass. NPF2.13 and 3.1 accumulated higher levels of the Q3R than the rest of the transporters of the 1^st^ and 2^nd^ clades (Sup. 22), whereas the discriminating signal for K3G was too low to elaborate (Sup. 23).

Similarly, the high signal for an isomer of sinapoyl glucose in the media (SG) translated in most of the 14 NPF transporters accumulating negligible relative levels to the media (Sup. 24). Mock oocytes also accumulated levels of a metabolite that either coeluted with SG and had the same formula or represented endogenous oocyte SG import. The exception was NPF1.1, which accumulated two of the less abundant isomers at similar levels to the media. We could annotate disinapate hexoses (DSH) as substrates of NPF1.1, 2.5, 2.13, and NPF3.1 (Sup. 25). DSHs are intermediates in the phenylpropanoid pathway (Fraser et al., 2007; Lee et al., 2017). Notably, NPF1.1 and 2.5 displayed comparable accumulations to media levels towards different isomers of DSHs, which were likely isomers of disinapoyl glucose or disinapoyl fructose. NPF 2.13 and 3.1 accumulated most DSH isomers proportionally to media levels, suggesting less stereospecificity.

Most of the confirmed or annotated imported features contained sugar moieties. The exception was sinapoyl malate (SM), an important UV protectant (Dean et al., 2014), which accumulated in NPF5.2 and to a lower extent in NPF3.1 (Sup. 26).

Once we assessed phytochemical import in the NPF-expressing oocytes, we wanted to compare their chemical profiles as previously done. We performed the biclustering analysis of the 100 most significant features for the eight transporters with validated substrates to show their shared substrate specificities (Fig. 7). The unsupervised biclustering analysis separated all the oocytes according to the transporter they expressed, except for NPF5.2, which clustered with mock oocytes since there were no discriminating features accumulating in NPF5.2 in the top 100 most significant features. The GTRs clustered together, where NPF2.10 and NPF2.11 displayed the highest similarity, and NPF2.9 clustered closest to them. NPF2.13 and NPF3.1 clustered close to the GTRs since they also displayed a lower accumulation of glucosinolates. NPF1.1 and NPF2.5 clustered together and separately from the rest of the transporters since they did not accumulate the majority of glucosinolates, and they shared the accumulation of a DSH isomer. All significant features accumulated in transporter-expressing oocytes relative to mock, which indicated that the represented features likely stemmed from import.

**Fig 7.**
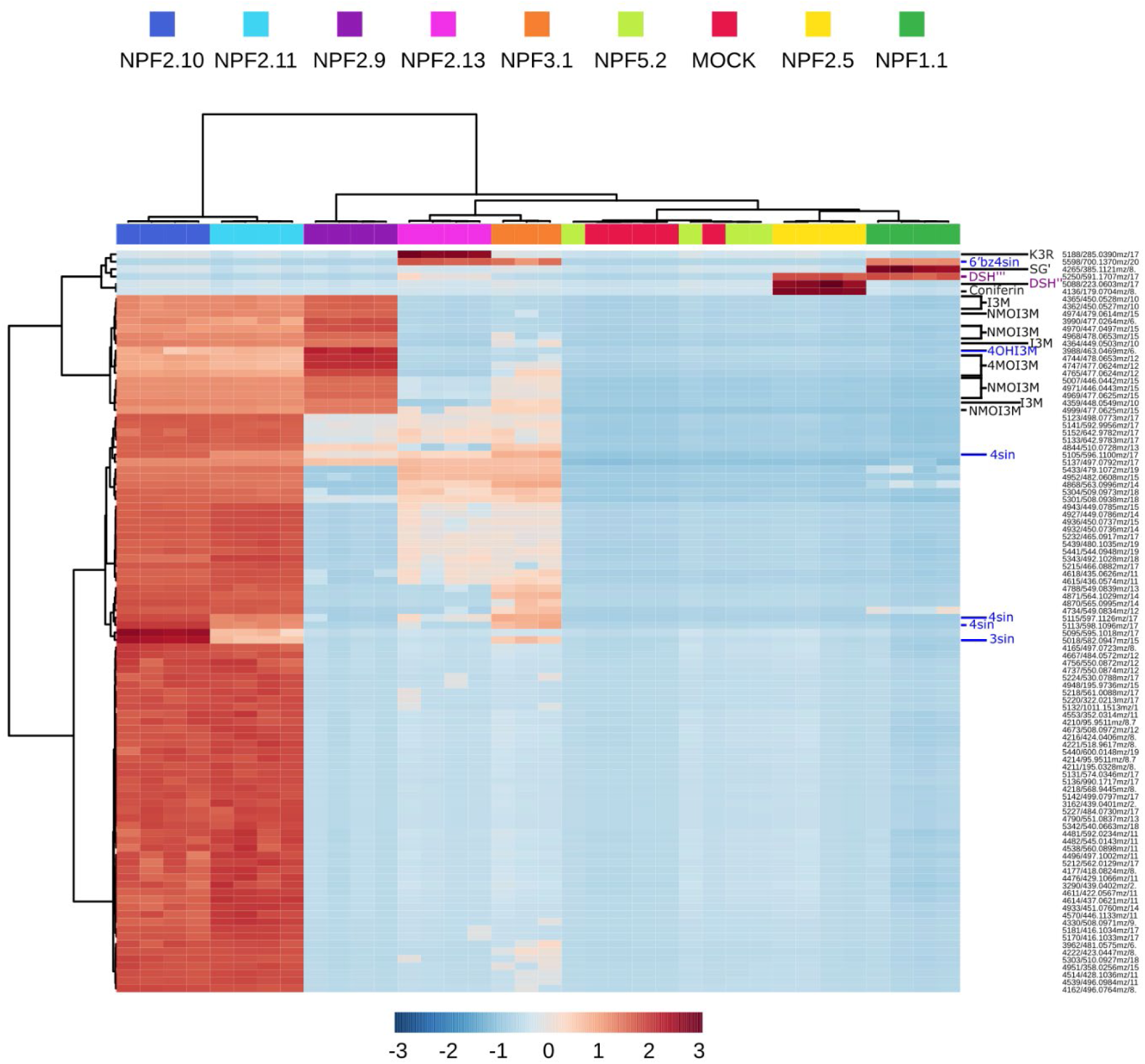
**Glucosinolate import generates most of the clusters separating selected NPF substrate spectra, while phenylpropanoid import contributes to the separation of some additional samples**. Unsupervised bicluster analysis of control (mock), and selected NPF-expressing oocyte samples assayed with 1:20 dilution of seedling-derived media. The relative intensities of metabolic features are represented on a color scale. Columns represent samples and rows metabolic features. Metabolic features identified with chemical standards are in black: glucobrassicin (I3M), neoglucobrassicin (NMOI3M), 4-methoxyglucobrassicin (4MOI3M), and Kaempferol-3-O-rhamnoside (K3R). Annotated glucosinolates in blue font: 3-sinapoyloxypropylglucosinolate (3sin), 4-sinapoyloxybutylglucosinolate (4sin), 4-Hydroxyglucobrassicin (4OHI3M), and 6′-O-benzoyloxy-4-sinapoyloxybutylglucosinolate (6’bz4sin). Features deriving from annotated phenylpropanoids in purple: Two isomers of disinapoyl hexose (DSH’’ and DSH’’’). Ward’s biclustering of the 100 most significant features (ANOVA). Sig. P-value <0.05. Oocytes were assayed at pH 5 for 1 hour with 1:20 dilutions of seedling extract; replicates consisted of five oocytes each. Features displaying a Relative Standard deviation of >30% in the QC were filtered. Features are logarithmically transformed (Log10) and Pareto scaled.

The most represented features in the heatmap could be annotated as glucosinolates, some of which we confirmed with chemical standards. Amongst these, indole glucosinolates still accumulated the highest in NPF2.9, forming their separate cluster, and the annotated sinapoylated glucosinolates 3sin and 4sin also separated NPF2.10 and NPF2.11. NPF 3.1 and NPF 2.13 accumulated lower relative levels of glucosinolates than NPF2.10 and 2.11, except for the annotated 6’bzo4sin. The different levels of 4sin and K3R contributed to the separation of NPF2.13 from NPF3.1. Some phenylpropanoids, like the first isomer of SG or the second isomer of DSH, accumulated differentially into NPF1.1 and NPF2.5, contributing to their separation.

Glucosinolates dominated the hundred most significant features in the multiple comparison of the oocytes, which was likely a combination of their diversity in *A. thaliana* (Blažević et al., 2020), their levels, the suitability of the analytical method, and the capacity of the GTRs to accumulate them above the media. Nevertheless, provided that the annotations were correct, the inclusion of more transporters in the biclustering enabled the observation of the previously described accumulation patterns, which corresponded to a greater variety of molecular classes of substrates.

NPF5.2 accumulation of sinapoyl malate (SM) was the first reported transporter activity towards SM. We wanted to confirm this observation due to the physiological relevance of SM as a photoprotector (Baker et al., 2020). We screened the entire AtNPF5 clade with pure SM to verify the feature’s identity and transport, and to evaluate the relative import activity towards the pure compound (Fig. 8). Most of the NPF5 clade did not accumulate significant levels of SM. SM import was restricted to the NPF5.2 subclade (NPF5.1, 5.2, and 5.3) (Léran et al., 2014), and NPF5.2-expressing oocytes accumulated SM to levels lower or comparable to the media, depending on the volume assumed for the oocyte (Stegen et al., 2000). The variability in SM accumulation in NPF5.2 transporter-expressing oocytes was high, likely due to differences in transporter expression or oocyte volume in the single-oocyte samples. NPF5.2 accumulation relative to the media was higher from the pure compound, which was performed at an external concentration of 20 µM. The levels of SM in the seedling extract are noticeably lower than 10 µM (Sup. 26), which could explain this relative difference. Alternatively, the import of other undetected compounds from the seedling media could have competed with SM import. Regardless, SM import is an expansion of the suggested peptide substrate spectrum of NPF5.2 (Karim et al., 2005).

**Figure 8.**
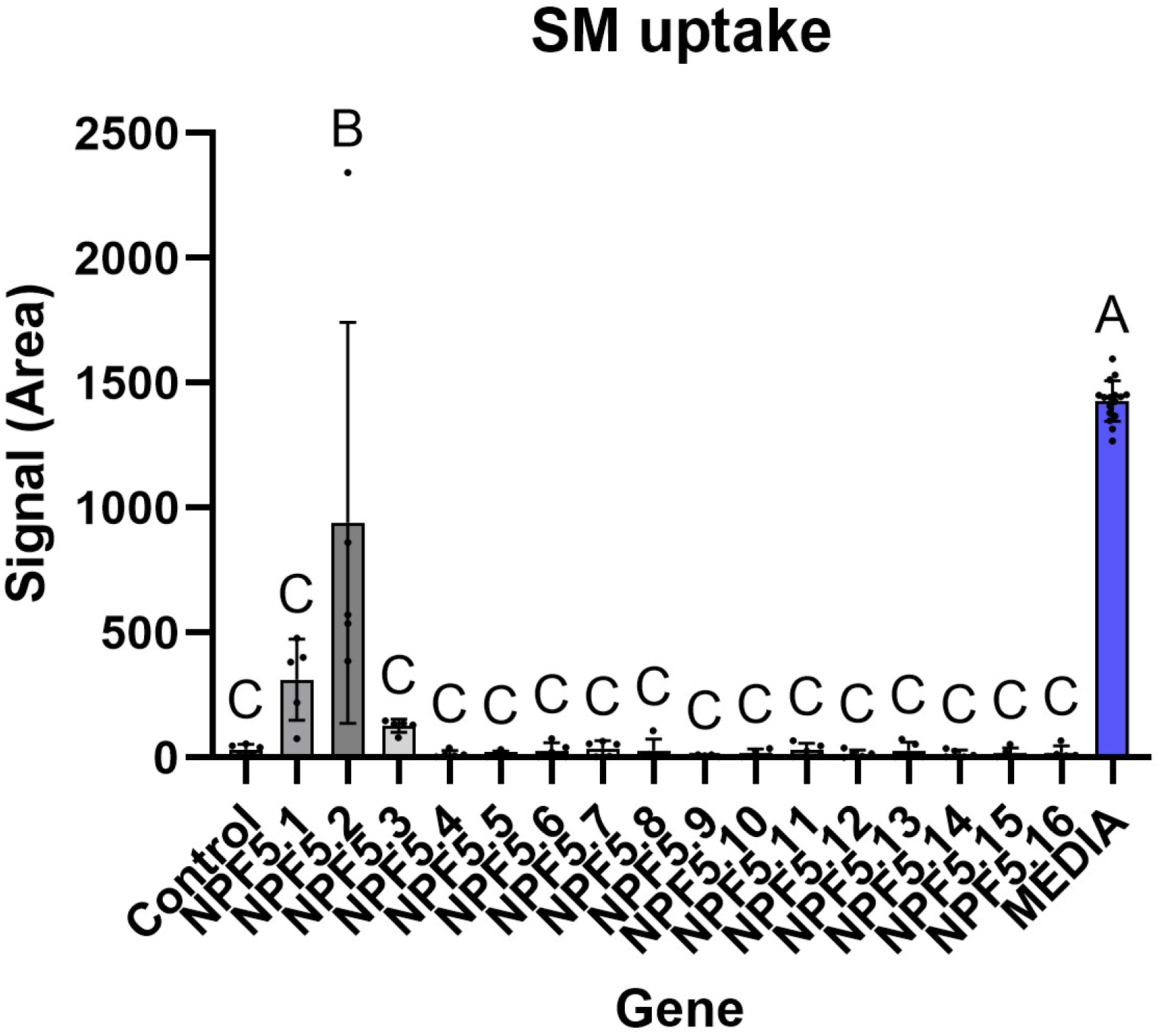
AtNPF5.2 imports an orphan metabolite. Targeted transport detection of pure sinapoyl malate (SM) in transporters of the NPF5 clade. Oocytes were assayed at pH 5 for 1 hour with 20µM of SM; replicates consisted of one oocyte each. Bars show the averages of 5 individual oocytes (n = 5) measured with multiple reaction monitoring (MRM), and error bars represent the standard deviation. The media signal corresponds to the signal equivalent to 1µl of external media. Letters represent the statistically different groups for a Tukey Multiple comparison test preceded by One-way ANOVA. Sig. P-value <0.05.

In this work, NPF 2.13 and 3.1 transported a broad and diverse array of compounds, including glucosinolates, flavonols, and sinapate esters in a non-stereospecific manner compared to NPF1.1 and 2.5. On top of that, previous reports claim nitrite and gibberellin transport for NPF3.1 (Sugiura et al., 2007; Tal et al., 2016) and nitrate, phytohormones, and the exogenous tunicamycin for NPF2.13 (Chiba et al., 2015; Fan et al., 2009; Liu et al., 2023; Nour-Eldin et al., 2012). Additionally, they could transport the molecules with a higher molecular weight that we annotated. We wanted to test this promiscuity with a pure sinapate ester, so we assayed the NPF 1, 2, and 3 clades with DiSinapoyl Sucrose (DSS), the only disinapate ester commercially available at the time of this work, and with a sizeable mass of 754 Daltons.

Most of the NPF 1, 2, and 3 transporters did not accumulate significant levels of DSS (Fig. 9). NPF2.13 displayed the highest accumulation of DSS, followed by NPF3.1 and 1.2, and lower levels in NPF2.12 transporter-expressing oocytes. NPF2.13 accumulated DSS over media levels, unlike the rest of the transporters. The variability in DSS accumulation in NPF2.13 transporter-expressing oocytes was high, likely due to differences in transporter expression of the single-oocyte samples or oocyte volume. The accumulation of DSS in NPF2.13 and NPF3.1 reinforced our hypothesis that these two transporters are highly promiscuous. NPF1.2 accumulation of DSS could indicate NPF1 clade activity towards glucosides decorated with phenolic rings, since most of the NPF1.1 substrates in this work contained this motif.

**Fig 9.**
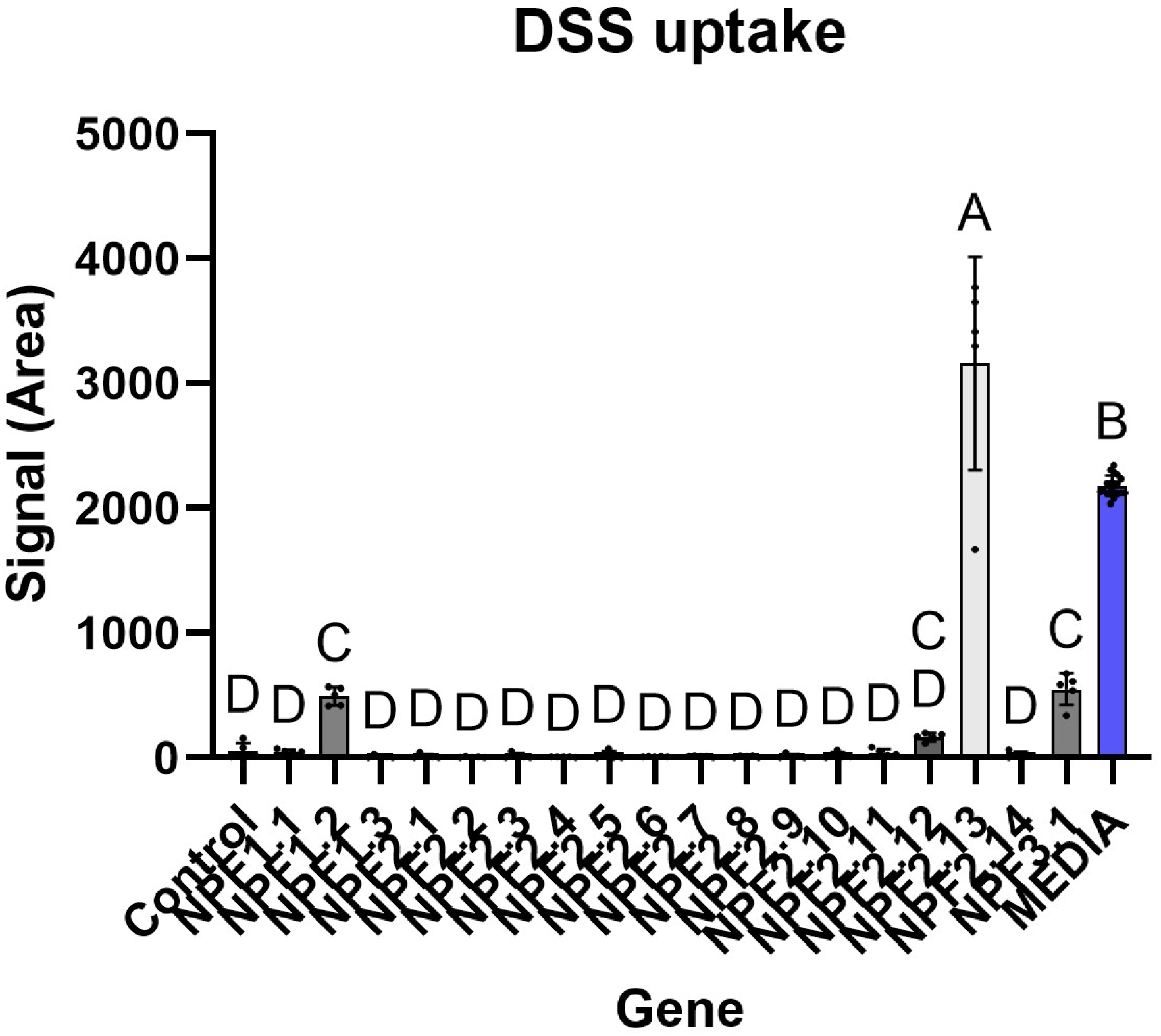
AtNPF2.13 and NPF3.1 can transport additional sinapate esters. Targeted transport detection of pure disinapoyl sucrose (DSS) in NPF1, 2, and 3 transporters. Oocytes were assayed at pH 5 for 1 hour with 20µM of DSS; replicates consisted of one oocyte each. Bars show the averages of 5 individual oocytes (n = 5) measured with multiple reaction monitoring (MRM), and error bars represent the standard deviation. The media signal corresponds to the signal equivalent to 1µl of external media. Letters represent the statistically different groups for a Tukey multiple comparison test preceded by One-way ANOVA. Sig. P-value <0.05.

## DISCUSSION

### Extract formulation

In this work, we have shown how the untargeted analysis of transporters exposed to metabolic extracts is a viable data-driven strategy to expand transporter substrate spectra. As a first step, we adjusted the metabolic extract dilution to alleviate the non-transporter-mediated permeation of metabolites into the oocyte. Various phytochemicals can alter the membrane fluidity of different organisms (Arora et al., 2000; Bard et al., 1988; Blankemeyer et al., 1997), which could explain the permeating capacity of the extracts above a certain dilution threshold. On top of that, the increased osmolarity of the undiluted extracts could explain the permeating capacity of the extracts we used. Thus, it is unsurprising that diluting undefined media is a successful strategy for performing transporter assays from complex mixtures (Krumpochova et al., 2012). Nevertheless, there does not seem to be a categorical threshold for the permeating capacity of the extracts according to the dilutions that we tested. The determination of the dilution factors indicated for oocyte membrane tightness will likely depend on the sensitivity of the analytical method and the choice of metabolites to make this extrapolation. Assays using plant extracts can render successful metabolite differences between control and transporter-containing membranes, even when the control counterparts might display metabolite leakage background levels (Demurtas et al., 2019).

Permeation likely changes from metabolite to metabolite, and it probably describes a membrane permeation continuum with media dilution. On the other hand, metabolite concentration has to be high enough for the transporters to display activity, depending on their affinities. Thus, the optimal dilution of the media probably peaks in a range while there is still some permeation of metabolites into the cell (or metabolite adsorption to the membrane). Additionally, one cannot rule out residual import by the endogenous transportome of the oocyte. We observed trace signals for phytochemicals in our control oocytes, as observed for sinapoyl glucose.

A significant technical limitation of approaches such as ours is the ability to expose the relevant metabolites to the transporters and detect and annotate them (Gründemann et al., 2005). To address this limitation, we hypothesized that changes in transporter expression correlate with substrate presence in their native tissues. Alternatively, screening with increasingly complex mixtures could increase the chances of extracts containing the relevant substrates. We have shown that the screening outcome depends on the material used for extract generation. Thus, mixing materials from different developmental stages, biosynthetic mutants, stressed plants, or even mixing organisms constitutes strategies to increase the likelihood of including transporter substrates, or at least structurally similar substrates to the physiological ones. While potentially beneficial, it could also increase the complexity of the data and the likelihood of including chemicals that inhibit transporters or permeate the oocyte membrane.

Metabolic extracts pose other challenges related to the benefits of screening with undefined mixtures. When screening with extracts, one runs a competition assay with many unknown substrates (Krumpochova et al., 2012), many of which are not detectable due to limitations on the analytical setup. In addition, the concentration of the metabolites can translate into different signal levels according to metabolite ionization efficiencies(Oss et al., 2010). Thus, it would be advisable to purify the particular compounds to assess the transporter relative import activities compared to the media and subsequently confirm the chemical identities of the compounds.

### Accumulation patterns to reflect the molecular determinants of substrate specificity

The method successfully reflected the transport activities of NPF2.9 compared to NPF2.10/2.11 from pure compound assays (Jørgensen et al., 2017). The ability of NPF2.9 to accumulate indole glucosinolates to a greater extent than NPF2.10 and 2.11 is likely due to the competition of NPF2.10 and 2.11 from other glucosinolates. In addition, the non-glucosinolate uptake, such as that of Q3R, reflects the GTRs’ ability to transport other molecules, and how these describe other accumulation patterns (Saito et al., 2015; Y.-Y. Wang & Tsay, 2011). The bicluster analysis takes the information level a step further, providing information at a glance on the relevance of individual features and how these accumulate differentially in the different oocytes and depending on the transporter. These multiple comparisons revealed big differences in oocyte chemical profile stemming from different extracts and also the subtle sub-specificities between functionally similar transporters (NPF2.10 vs NPF2.11) exposed to the same extract. The relative activities of NPF2.10 and NPF2.11 towards sinapoylated glucosinolates and benzoylated glucosinolates would be the first report of a difference in substrate specificity towards native glucosinolates for these two transporters. Two different extracts revealed NPF2.10’s sub-specificity towards sinapoylated glucosinolates, one used with two different dilutions (seedling). The only difference published for these two genes towards glucosinolates applies to artificial glucosinolates (Kanstrup, Jimidar, et al., 2023). Differences in affinity towards different glucosinolates have been reported for NPF2.10 according to their hydrophobicity, but not while comparing NPF 2.10 and NPF 2.11 (Chung et al., 2022).

The differential accumulation of sinapoylated glucosinolates and benzoylated glucosinolates across the GTRs (and NPF’s) exemplifies three advantages of this method. First, we can detect uptake for commercially unavailable developmental-stage-specific metabolites. Sinapoyl and benzoyl-decorated glucosinolates are described in seeds (Kliebenstein et al., 2007; Lee et al., 2017; Reichelt et al., 2002), which reflects the advantages of using different plant materials as metabolite sources to find unreported transporter activities.

While comparing substrate preferences of two or more transporters is usually carried out with single compounds, there are exceptions to this (Fairweather et al., 2021). In our case, we found substrate activity differences for compounds that are not commercially available, and the usage of complex media allows us to formulate hypotheses in substrate preferences that are very valuable for their posterior confirmation from purified compounds, essentially a data-driven approach.

We suggest that similar transporters are likely competed by similar substrates. Thus, when comparing closely related transporters such as NPF2.10 and 2.11, the transporter activity towards a particular molecule is likely competed by a similar range of other molecules. Therefore, the specificity of NPF2.10 towards a sinapoylated glucosinolate versus NPF2.11 is contextualized by their shared simultaneous competition of all the other glucosinolates that we can detect, plus the metabolites that we cannot detect. Therefore, generating hypotheses in the specificity of a transporter towards structural motifs benefits from comparing transporter activities with their closest homologs.

The second advantage refers to the daunting task of feature annotation in LC-MS. A big challenge in metabolomics is the correct annotation of metabolites (da Silva et al., 2015), which is an issue with more severe implications in understudied organisms, like plants. We used the knowledge of the model plant *A. thaliana* to alleviate this issue. Fortunately, there has been a serious development in recent years for the structural annotation of mass spectrometry features independent of the existence of experimental standardized spectral databases (Dührkop et al., 2019; Heuckeroth et al., 2024; Schmid et al., 2021; M. Wang et al., 2016; Watrous et al., 2012). We have used the extracts to reveal the selectivity of the transporters, but conversely, one could use transporters to screen complex metabolic mixtures for structurally similar molecules. The selectivity that a transporter confers the oocyte membrane transforms the oocyte into a collection of structurally related imported compounds. The domain knowledge of *A. thaliana* metabolism and the accumulation pattern of these features across several transporters and relative to the media levels allowed us to annotate some challenging features. Thus, it was possible to annotate heavily decorated glucosinolates like sinapoylated glucosinolates or 6’bz4mtb using accumulation patterns in NPF2.10 vs NPF2.11 and literature (Clarke, 2010; Reichelt et al., 2002). When assaying more transporters simultaneously, we observed unexpected activities such as the activity of NPF1.1 towards 6’bz4mtb. We could identify the accumulation pattern of 6’O-benzoylated glucosinolates in NPF1.1 2.13 and 2.13, which allowed us to hypothesize novel structures to annotate some imported features. Depending on the correct annotation of these two features, we could hypothesize a novel specificity of NPF1.1 towards a specific subset of glucosinolates. Alternatively, taken together with the activity towards sinapate esters, one could suggest that NPF1.1 displays activity towards molecules containing sugars decorated with phenyl rings. The comparison of transporter activity towards a vast collection of metabolites offers insight into which molecular patterns are important for transporter function and showcases how comparing the activity of many transporters towards the same mixture boosts the insight of the information acquired.

We annotated that the accumulation patterns of features within our data correspond to the preferences of the transporters towards given molecular moieties. These observations enable the strenuous task of metabolite annotation, one of the biggest challenges in mass spectrometry, especially for non-human metabolites. This strategy complements the analytical chemistry tools and facilitates the annotation of features with poor-quality MS2 data, or the challenging MS2 data derived from precursor ions with high mass.

Mass spectrometry allows for investigating metabolomes at motif level-with motifs being features representing MS2 fragments corresponding to recurring molecular moieties (van der Hooft et al., 2016). Thus, it is theoretically possible to examine the specificities of transporters towards the fragments of their substrates instead of the substrates themselves. In this work, we display and compare the activities of transporters towards MS1 features. Alternatively, transporters could be compared according to the MS2 motifs of their substrates. We think that implementing new data analysis techniques, especially the ones that already use MS2 data for metabolite pattern mining, will make the most of our approach (Schmid et al., 2021). The correlation of molecular patterns to the substrate specificity of transporters will offer invaluable insight into the structural determinants of substrate specificity. Linking transporters to molecular motifs will allow us to offer a mechanistic data-driven classification of their activity, opposite to the one offered by targeted approaches with a limited number of compounds.

The third advantage of the approach is assessing functional gene expression of transporters without the observation of transport activity. It is challenging to ensure the functional expression of transporters without a known substrate. These proteins can require post-translational modifications, protein stabilization, or the formation of multimers to display their full activity potential (Abplanalp et al., 2013; Orsel et al., 2006). Thus, exposing the transporters to many substrates increases the chance of observing transport activity and, therefore, being able to claim (at least partial) functional expression.

The functional expression of transporters adds uncertainty when interpreting their activity in heterologous hosts. All of the transporters we selected have been functionally expressed in oocytes (Binenbaum et al., 2023; Dietrich et al., 2004; Hsu & Tsay, 2013; Jørgensen et al., 2017; J.-Y. Li et al., 2010; Morales de los Ríos et al., 2021; Tal et al., 2016; Tsay et al., 1993; Wulff et al., 2019). Nevertheless, only eight of them displayed evident transport activity toward the detectable metabolites. We chose to only elaborate on the activities of transporters with verifiable transporter activity. Some of the previously described substrates for the discarded transporters, such as nitrates, are not typically applicable to LC-ESI-MS. Others, like the phytohormones, are presumably in too low concentrations, and the oocyte could have metabolized primary metabolites (Dworkin & Dworkin-Rastl, 1989). To alleviate these limitations, we could optimize sample preparation (our current protocols dilute the oocyte 30-fold for sample preparation), add different analytical techniques in parallel (like GC-MS, which is more suitable for the detection of primary metabolites and lipids), or optimize the extraction of metabolites, to name a few.

### Expansion of the NPF substrate spectra

Some of the new transport activities described here hold intrinsic biological and biotechnological value, such as the transport of sinapate glycosides and monolignol glycosides by NPF1.1 and NPF2.5 (Lee et al., 2017). To our knowledge, no transporters have been linked to the translocation of DSH, and only one ABC transporter has been linked to the transport of the monolignol p-coumaryl alcohol (Alejandro et al., 2012). We could not detect the presence of p-coumaryl alcohol glucoside in our media, but we detected the transport of the glycosylated forms of the other two main blocks of lignin biosynthesis, sinapyl and coniferyl alcohol, by NPF2.5. The transport of these compounds has been hypothesized to be exploited *in planta to* modify lignin biosynthesis (Lee et al., 2017).

We were also able to annotate flavonol transport. Exporters from the ABC and MATE families have previously been linked to the transport of the related anthocyanins (Behrens et al., 2019; Marinova et al., 2007). AtNPF2.8 was the first gene from the NPF family to be linked with flavonol import in planta (Grunewald et al., 2020), a gene closely related to the ones described here. These intriguing molecules regulate phytohormone activity (Tan et al., 2019) and modulate phytohormone transport (Brown et al., 2001). Additionally, the flavonoid Naringin is a transporter inhibitor (Bailey et al., 2007). The widespread activity of many NPFs towards flavonols suggests that these molecules are modulators of transporter activity.

The common denominator for most of the substrates, from glucosinolates to phenylpropanoids, confirmed or annotated, is glycosylation. This observation is interesting from a mechanistic and a physiological perspective. The activity from transporters to molecules with common structural features can help understand the structural determinants of substrate specificity, as observed for the SUC transporters (Bavnhøj et al., 2023; Rottmann et al., 2018). Hence, one could claim that NPFs from clades 1,2, and 3 display activity to molecules that contain sugar moieties, where additional motifs help to tune their activity. Altogether, we can hypothesize that the apparent promiscuity of transporters is a consequence of activity on substrates with shared structural motifs, which allowed us to find transporters for a commercially available compound without a transporter, like DSS.

From a physiological perspective, glycosylation changes specialized metabolite availability and localization (Dima et al., 2015; Francisco et al., 2013; Miao & Liu, 2010). The differential selectivity of transporters towards glycosylated molecules explains the ability of the plant to redirect the metabolite precursors to cellular compartments. Additionally, some metabolites display enhanced toxicity to the plant in their aglycon precursor (Alejandro et al., 2012; Väisänen et al., 2015), pointing to the role of transporters in detoxification, sequestration, and metabolic biosynthesis. Interestingly, glycosylation is one of the modulation mechanisms of phytohormones (K. Chen et al., 2020; Jackson et al., 2002; Jin et al., 2013; Noutoshi et al., 2012; Poppenberger et al., 2005), and transporters likely play a role in the redirection of these phytohormone conjugates.

Our wider screening suggests assessing substrate promiscuity before interpreting transporter function. A certain degree of promiscuity is expected for transporters (Jørgensen et al., 2017). We have found transporters with various degrees of promiscuity and an overall capacity of the NPF1, 2 and 3 clades to transport very different molecules. Based on literary evidence and present data, certain transporters such as NPF2.13 and 3.1 seem relatively more promiscuous within our detectable chemical space. Coincidentally, they are linked with hormone transport (Binenbaum et al., 2023; Tal et al., 2016). This poses major questions in interpreting their role in complex processes such as development, specifically the phenotypes that their mutants display. Based on their multispecificity and the structural diversity of phytohormones, their *in vitro* activity towards phytohormones is unsurprising. Insights into NPF2.13 and 3.1 promiscuity have built up from different targeted studies, such as the discovery of glucosinolate, phytohormone, nitrate, and tunicamycin transport from NPF2.13 (Chiba et al., 2015; Fan et al., 2009; Liu et al., 2023; Nour-Eldin et al., 2012). These approaches were not aimed at reporting comprehensive substrate spectra of a transporter, so they were not incentivized to describe their promiscuity. Thus, exposure to multiple compounds at a time would have revealed their promiscuous nature.

The promiscuity of these transporters questions the causality of hormone transport in the mutant phenotypes. For example, the *in vivo* import of fluorescently tagged GA by NPF3 could be due to NPF3.1 promiscuity (Tal et al., 2016), or the activity of the hypothetical *in vivo* glycosylation of these exogenous molecules. Analogously, the promiscuity of NPF2.13 is patent in the *in vitro* data we show, supported by its *in vivo* ability to transport tunicamycin, an exogenous glycosylated antibiotic (Liu et al., 2023). Consequently, we think approaches such as ours are instrumental in reassessing the current transporter roles and their multispecificity/promiscuity in particular. In evolutionary terms, multifunctional transporters with the capacity to specialize in a specific chemical space through duplication and neofunctionalization enable plants to evolve new metabolites and functions for the preexisting metabolites and thus are beneficial for the plant population (Jørgensen et al., 2017). Therefore, promiscuity is likely to be a relevant physiological trait other than an artifact. Hypothesis-driven targeted approaches are particularly susceptible to linking a promiscuous transporter substrate with the pleiotropic effects of a promiscuous transporter (Pochini et al., 2022). The targeted approaches for discovering transporters for phytohormones molecules may have found promiscuous proteins instead.

## CONCLUSIONS

We show that studying metabolite transport in plants can enormously benefit from data-driven approaches such as the one described here and that *X.laevis* oocytes provide an excellent platform for these endeavors. We have discovered new substrates for the NPF family and developed a tool to compare transporter substrate spectra in a less biased manner. The individual specificities need to be validated *in vivo*, but the observation of multispecificity is transformative in interpreting the transporter’s role. Additionally, this method uncovers shared substrate specificities that can be instrumental in generating higher-order mutants to confirm these interactions *in vivo.* We foresee that optimizing this approach and implementing the new bioinformatics tools, such as AI and molecular docking models with the new substrates, will hold insights into the determinants for the specificity mechanism and new transporter functions.

## Materials and methods

### Chemical standards

Neoglucobrassicin (NMOI3M), glucobrassicin (I3M), 4-methoxy-3-indolylmethyl glucosinolate (4MOI3M) sinalbin (pOHB), glucoraphanin (4msb) afzelin (K3R), isoquercetin (Q3G), quercitrin (Q3R), and astragalin (K3G) where purchased from Extrasynthese. 3,6’-Disinapoyl sucrose (DSS) and syringin were purchased from Biosynth. Coniferin/abietin was purchased from Sigma-Aldrich (SMB00103) Glucoerucin (4mtb) was purchased from C2 Bioengineering, Karlslunde, Denmark. Sinapoyl malate (SM) and sinapoyl glucose (SG) were a generous gift from Christoph Hald and Corinna Dawid from the Technical University of Munich.

### Oocyte media

Salts and the buffering agents for the Kulori (oocyte) media were purchased from Sigma-Aldrich. Oocytes were incubated in Kulori pH 7.4 before the assay (90 mM NaCl, 1 mM KCl, 1 mM MgCl_2_, 1 mM CaCl_2_, 5 mM HEPES adjusted to pH 7.4 with NaOH). Kulori pH 7.4 was supplemented with 100 µg/ml of amikacin. Assays were performed in Kulori pH 5 (90 mM NaCl, 1 mM KCl, 1 mM MgCl_2_, 1 mM CaCl_2_, 5 mM MES adjusted to pH 5 with NaOH).

The generation of plant extract derived media is detailed below.

### Seed propagation

To generate phytochemical extracts, mature *Arabidopsis thaliana* ecotype Columbia 0 (Col-0) seeds were bulk harvested from dried plants.

Plants were grown in 11 cm diameter pots of 8cm depth in a mixture of 4 volumes of Pindstrup peat moss (Denmark) with 1 volume of Sand nr. 2 DANSAND® (Denmark). The soil mix was heat-treated at 80°C in tap water saturation (Frederiksberg C., Denmark) before sowing. Three seeds were distributed per pot and stratified in the dark for 4 days at 4°C. Plants were grown in chambers in long-day conditions, 16h light at 21°C and 65% relative humidity/ 8 h dark at 19°C and 70% relative humidity, for approximately three months. Light intensity was measured at rosette level to be 150 µmol m–2 s–1. Plants were gradually tap-watered once the first 0.5cm of soil was visibly dry, and all the stems of the inflorescences were attached to a bamboo stick. Once one third of the plant’s siliques were mature, the bolts were wrapped in a paper bag, watered, and let to dry in the same chamber. After the complete plants were dry (2 weeks approximately) seeds were harvested and sieved.

Seeds were used at least 1 month after harvesting, but not longer than 1 year after harvesting. These seeds were considered to consist of 100% dry material for extract generation purposes.

### Seed extraction

400 mg of seeds were divided into aliquots of 40mg, in 1.5 ml conical tubes with two 3mm steel balls. 1ml of 70% Methanol (v/v) (≥99.9%, HiPerSolv CHROMANORM®) was added per aliquot and immediately homogenized in a homogenizer mill. The aliquots were homogenized at 30 Hz in 4 cycles of 2 minutes, randomizing tube distribution in the adaptor between cycles to ensure homogenous tissue disruption. The samples were centrifuged at 4°C for 10 minutes at 20000 *g* to pellet plant debris. The supernatants were pooled and then divided into two 5ml conical tubes. The pooled supernatants were dried completely in a SpeedVac™ connected to a CoolSafe™ at - 110°C and stored at-20 °C until resuspension.

Each tube contained the extract equivalent to 200mg of seed extract; accordingly, the pellet of the tube was resuspended in 2 ml of Kulori buffer at pH 5 with the aid of a sonicator bath (Branson 5510, 42KHz, 135 W) at room temperature and a clean metal spatula. This extract was considered 1:1 seed dilution, which was further diluted with Kulori buffer at pH 5.

### Seedling extraction

The propagated seeds were plate-grown axenically for 2 days under continuous light in ½ Murashige-Skoog (MS) agar media supplemented with 3% sucrose, matching the growing conditions described by (Narsai et al., 2011) but with half the light intensity.

150 mg of seeds were divided into 15 mg aliquots in conical tubes. Seed aliquots were surface-sterilized with 1 ml of 70% ethanol and 0.01% Triton X-100 for 10 minutes. Then the seeds were washed two times with 96% ethanol and left to dry in Whatman™ paper. Then seeds were distributed by carefully sprinkling them on a 12×12 cm square petri dish (688102, Greiner) with a media layer of 0.5-0.7. The media consisted of ½ Murashige-Skoog (MS) salts with vitamins (M0222, Duchefa) with 3% Sucrose (S0809, Duchefa) and 1% agar (M1002, Duchefa). Plated seeds were stratified in the dark for 2 days at 4°C. Then they were germinated for two days under continuous light (50 µmol m–2 s–1) at plate level for 2 days at 80% humidity. The seedlings were checked for fungal growth and carefully harvested from the agar. Seedlings were dried with Whatman™ paper and divided into 200 mg aliquots in conical tubes with two 3mm steel balls. The aliquots were flash-frozen in liquid N_2_. The total amount of seedling material was measured. The seedling aliquots were homogenized to a fine powder in a mill at 30 Hz in short intervals of 30 seconds. The adaptors were kept in cold (-20°C) and between each cycle, the seedling aliquots were flash-frozen to avoid thawing.

Seedling water content was measured to be around 85% by lyophilizing. Accordingly, each 200mg seedling powder aliquot contained 170µl of water which was mixed with 400µl of >99.9% methanol to a final dilution of 70% methanol and immediately homogenized with the mill for 4 cycles of 2 minutes. The samples were centrifuged at 4°C for 10 minutes at 20000*g* to pellet plant debris.

The supernatants were pooled and then divided into two 5ml conical tubes. The pooled supernatants were dried completely in a SpeedVac™ and stored at-20 °C until resuspension. 1 ml of Kulori buffer pH 5 was used per 100mg of dried seedling material, which was calculated according to 15% of the total initial seedling weight. Kulori buffer at pH 5 was used for the resuspension, with the aid of a sonicator and a metal spatula. This extract was considered 1:1 seedling dilution, which was further diluted with Kulori buffer at pH 5.

### cRNA generation

Plasmids containing transporter cDNA for cRNA generation were described by (Wulff et al., 2019). Briefly, *Xenopus laevis’* β-Globin 5’ UTR linked to transporter cDNA sequence was amplified using Phusion™ polymerase (New England Biolabs, U.S.A) according to the manufacturer’s indications with a forward primer (5′-AATTAACCCTCACTAAAGGGTTGTAATACGACTCACTATAGGG-3′) and a reverse primer (5′-TTTTTTTTTTTTTTTTTTTTTTTTTTTTTATACTCAAGCTAGCCTCGAG-3′) that incorporates the poly-A tail. The PCR amplicons were purified with E.Z.N.A Gel extraction kit (Omega Bio-tek, U.S.A) according to the manufacturer’s indications. The PCR amplicons were *in vitro* transcribed with the mMessagemMachine™T7 kit (Invitrogen™, U.S.A) according to the manufacturer’s indications, and normalized to an RNA concentration of 500 ng/µl.

### X. laevis’ import assays

30-40 Stage V-VI defolliculated *X. laevis* oocytes (Ecocyte Biosciences, Germany) were injected with 50.6 nl of cRNA solution for transporter expression or the same volume of nucleotide-free water (Invitrogen™, U.S.A) for the controls (mock). Glass capillaries were used to inject of 50.6 nl of cRNA (25 µg) or water using a Drummond Nanoject II and incubated for 4 days at 16°C in Kulori pH 7.4. The transporter-expressing oocytes were preincubated in Kulori pH 5 for 5 minutes before the assay. The oocytes were then transferred to the corresponding dilution of any of the extract formulations or the pure compounds and assayed for 1 hour. Then, oocytes were washed three times in 50 ml of Kulori pH5 and one time with 50 ml MiliQ water. Then, oocytes were visually inspected and the non-discarded oocytes were aliquoted, the supernatant removed, and the aliquots were kept on ice.

The oocyte aliquots were homogenized with 75 µl of 50% methanol with internal standards caffeine and sinalbin (para-OH-benzyl glucosinolate, pOHB), both at 2500 nM. The oocyte homogenates were centrifuged at 4°C for 15 minutes at 20000 *g* to pellet cellular debris. 67.5 µl of the supernatants were diluted with 102 µl of MiliQ water to a final dilution of 20% methanol and 500 nM internal standard, filtered through a 0.22 µM plate (MSGVN2250, Merck Millipore), and submitted to LCMS in 96-well Nunc ™ microplates (732-2623, Thermo-Fischer).

Media samples were prepared by aliquoting 3 or 5 µl of media after the oocyte assay. The spent media samples were mixed with 75 µl of 50% methanol with internal standards and processed parallelly to the oocyte samples.

QC sample for untargeted analysis was prepared by mixing a 5 µl aliquot of each oocyte sample, and in parallel, a 10µl sample of all the media samples. Equal volumes of media mix and oocyte mix were mixed and submitted in a glass vial as QC sample.

### Glucosinolate quantification by LC-MS-QqQ/MRM

Oocyte samples were subjected to analysis by liquid chromatography coupled to mass spectrometry. Chromatography was performed on an Advance UHPLC system (Bruker, Bremen, Germany). Separation was achieved on a Kinetex XB-C18 column (100 x 2.1 mm, 1.7 µm, 100 Å, Phenomenex, Torrance, CA, USA). Formic acid (0.05%) in water and acetonitrile (supplied with 0.05% formic acid) were employed as mobile phases A and B respectively. The elution profile was: 0-0.5 min, 2% B; 0.5-1.3 min, 2-30% B; 1.3-2.2 min 30-100% B, 2.2-2.8 min 100% B, 2.8-2.9 min, 100-2% B and 2.9-4.0 min 2% B. The mobile phase flow rate was 400 µl min^-1^. The column temperature was maintained at 40°C. The liquid chromatography was coupled to an EVOQ Elite TripleQuad mass spectrometer (Bruker, Bremen, Germany) equipped with an electrospray ion source (ESI). The instrument parameters were optimized by infusion experiments with pure standards. The ion spray voltage was maintained at 4000 V negative ion mode. Cone temperature was set to 350°C and cone gas to 20 psi. Heated probe temperature was set to 200°C and probe gas flow to 50 psi. Nebulizing gas was set to 60 psi and collision gas to 1.6 mTorr. Nitrogen was used as probe and nebulizing gas and argon as collision gas. Active exhaust was constantly on. Multiple reaction monitoring (MRM) was used to monitor analyte precursor ion → fragment ion transitions. The majority of transitions for the different glucosinolates were previously reported (Crocoll et al., 2016). Both Q1 and Q3 quadrupoles were maintained at unit resolution. Bruker MS Workstation software (Version 8.2.1, Bruker, Bremen, Germany) was used for data acquisition and processing. Linearity in ionization efficiencies were verified by analyzing dilution series. Quantification was performed either by external dilution series or by using the benzenic glucosinolate (sinalbin) as internal standard.

### Untargeted oocyte analysis by LC-MS-QToF

LC-MS/MS was performed on a Dionex UltiMate 3000 Quaternary Rapid Separation UHPLC^+^ focused system (Thermo Fisher Scientific, Germering, Germany). Separation was achieved on a Kinetex 1.7 μm XB-C18 column (100 × 2.1 mm, 1.7 μm, 100 Å, Phenomenex). For eluting 0.05% (v/v) formic acid in H2O and Acetonitrile [supplied with 0.05% (v/v) formic acid] were employed as mobile phases A and B, respectively. Gradient conditions were as follows: 0.0−1.0 min 2% B; 1.0−14.0 min 2−20% B; 14.0−20.0 min 20−45% B, 20.0−24.5 min 45−100% B, 24.5−26.5 min 100% B, 26.5−26.55 min 100−2% B, and 26.55−30.0 min 2% B. The flow rate of the mobile phase was 300 μL/min. The column temperature was maintained at 30°C. UV spectra for each sample were acquired at 229, 260, 310, and 345 nm. The UHPLC was coupled to a Compact micrOTOF-Q mass spectrometer (Bruker, Bremen, Germany) equipped with an electrospray ion source (ESI) operated in positive or negative ion mode. The ion spray voltage was maintained at +3750 V or - 4000 V in positive and negative ion mode, respectively. Dry temperature was set to 275 °C, and the dry gas flow was set to 8 L/min. Nitrogen was used as the dry gas, nebulizing gas, and collision gas. The nebulizing gas was set to 2.5 bar and collision energy to 10 eV in positive ion mode and 15 eV in negative ion mode, respectively. MS spectra were acquired in an *m/z* range from 50 to 1200 amu. and MS/MS spectra were acquired in a range from 200-800 amu. Sampling rate was 2 Hz. Na-formate clusters were used for mass calibration. All files were automatically calibrated by postprocessing.

### Analysis of Disinapoyl Sucrose and Sinapoyl Malate by LC-MS-QqQ/MRM

Samples were analyzed by liquid chromatography coupled to tandem mass spectrometry. Briefly, chromatography was performed on a 1290 Infinity II UHPLC system (Agilent Technologies). Separation was achieved on a Kinetex XB-C18 column (100 x 2.1 mm, 1.7 µm, 100 Å, Phenomenex, Torrance, CA, USA). Formic acid (0.05%, v/v) in water and acetonitrile (supplied with 0.05% formic acid, v/v) were employed as mobile phases A and B respectively. The elution profile for was: 0.0-0.2 min, 15 % B; 0.2-3.0 min, 15-65 % B; 3.0-3.5 min 65-100 % B, 3.5-4.4 min 100 % B, 4.4-4.5 min, 100-15 % B and 4.5-5.5 min 15 % B. The mobile phase flow rate was 400 µL/min. The column temperature was maintained at 40 °C. The liquid chromatography was coupled to an Ultivo Triplequadrupole mass spectrometer (Agilent Technologies) equipped with a Jetstream electrospray ion source (ESI). The ion spray voltage was set to-4000 V in negative ion mode. Dry gas temperature was set to 325 °C and dry gas flow to 11 L/min. Sheath gas temperature was set to 350 °C and sheath gas flow to 12 L/min. Nebulizing gas was set to 40 psi. Nitrogen was used as dry gas, nebulizing gas and collision gas. The instrument parameters were optimized for best detection of all metabolites with mixes of reference standards. Multiple reaction monitoring (MRM) was used to monitor precursor ion → fragment ion transitions (Sup. Table 9). Both Q1 and Q3 quadrupoles were maintained at unit resolution. Mass Hunter Quantitation Analysis for QQQ software (Version 10.1, Agilent Technologies) was used for data processing.

### Untargeted oocyte data pre-processing

The raw data files were assessed with Bruker Data Analysis software to ensure injection quality. The data were centroided and converted to.mzML format with Bruker Data Analysis software. The.mzML files were pre-processed with Mzmine 4.3.0. with the following parameters: Mass detection: Noise threshold MS_1_ =1000. Mass detection: Noise threshold MS_2_ =0. Chromatogram builder: Minimum consecutive scans =5; Minimum intensity for consecutive scans =1000; Minimum absolute height = 1000; Scan to scan accuracy(*m/z*) = 0.01 or 0.0ppm. Local minimum resolver: Dimension =Retention time; Chromatographic threshold 70%; Minimum search range RT =0.05; Minimum relative height = 0%; Minimum absolute height = 1000; Minimum ratio of peak top/edge = 1.7; Peak duration range 0.0 – 2 min, Minimum scans =5. 13 isotope filter: *m/z* tolerance =0.01 or 5ppm; Retention time tolerance =0.02 min.; Maximum charge= 2, Representative isotope =Most intense. Join aligner: *m/z* tolerance = 0.03 or 5ppm; Weight *m/ z* = 0.75; Retention time tolerance =0.3 min.; Weight for RT = 0.25. Feature list row filter: Minimum features in a row =2; Retention time =1-23 min. Gap filling: Intensity tolerance = 50%; m*/z* tolerance =0.3 or 5ppm; RT tolerance = 0.3 minutes; Minimum data points = 1. Statistics export (Metaboanalyst): Feature intensity = Area

### Data analysis with Metaboanalyst

Pairwise comparisons and Bicluster analysis were performed with www.metaboanalyst.ca online tools. Preprocessed files were uploaded for Peak intensities. No features with missing values were removed, and the missing values were imputed with the minimal value for every feature in the dataset. Features displaying more than 30% Relative standard deviation in the QC samples were removed, and no further feature filtering was performed according to variance and abundance. The normalization was performed according to the feature representing the molecular ion for the internal standard sinalbin (424.0324 *m/z)* and the data was Log10 transformed and Pareto scaled.

The bicluster analysis was performed on the selected transporters, on the normalized data, autoscaling for the features, the distance measured was Euclidean, and the clustering method was Ward’s. The represented features correspond to the top 100 most significant features according to the ANOVA test. (Sig. p-Value<0.05, adjusted according to the False Discovery Rate according to the Benjamini-Hochberg procedure)

Pairwise comparisons between transporter and mock were performed with the volcano plot tool, on the normalized data. The minimal fold change was set to 2, and the t-test was set with Sig. p-Value<0.05, adjusted according to the False Discovery Rate.

### Metabolite annotation

Features coeluting with the chemical standards listed above (less than 0.1 minutes difference), that presented exact mass values (*m/z)* with less than 0.005 units of difference to the standards’, and with matching MS_2_ spectra, were assigned level 1 annotations (identification).

Most of the structures were first hypothesized on their nature according to their accumulation pattern across the transporters and the observation of characteristic MS_2_ fragments, like the sulfated ions for glucosinolates (Clarke, 2010), or the aglycon moieties of the flavonoids. Then their formula was extracted from the relevant literature referenced or assembled by structural homology (6’bzo4sin and 6’bzo3sin). The values for their corresponding ions were calculated with Bruker Data Analysis software. If the predicted mass was less than 0.005 units (*m/z)* different from the measured ion, the features were assigned to a level 3 annotation.

## Author Contributions

Víctor de Prado Parralejo and Hussam H. Nour-Eldin conceived and designed the study. Hussam H. Nour-Eldin supervised the study. Víctor de Prado Parralejo performed the experiments. Christoph Crocoll analyzed the samples. Víctor de Prado Parralejo and Christoph Crocoll analyzed the data. Christos Theodorou drew the structures. Victor de Prado Parralejo drafted the manuscript, which all authors edited.

## Acknowledgements

The Villum Foundation grant awarded to Villum Investigator Professor Barbara Ann Halkier (#37798) supported this work. We would like to thank Christoph Hald and Corinna Dawid from the Technical University of Munich for the sinapoyl malate and sinapoyl glucose standards.

## Data availability

All data are available upon request.

