## Supplementary figures and tables for "A versatile method to expand and compare transporter substrate spectra"

### Supplementary material

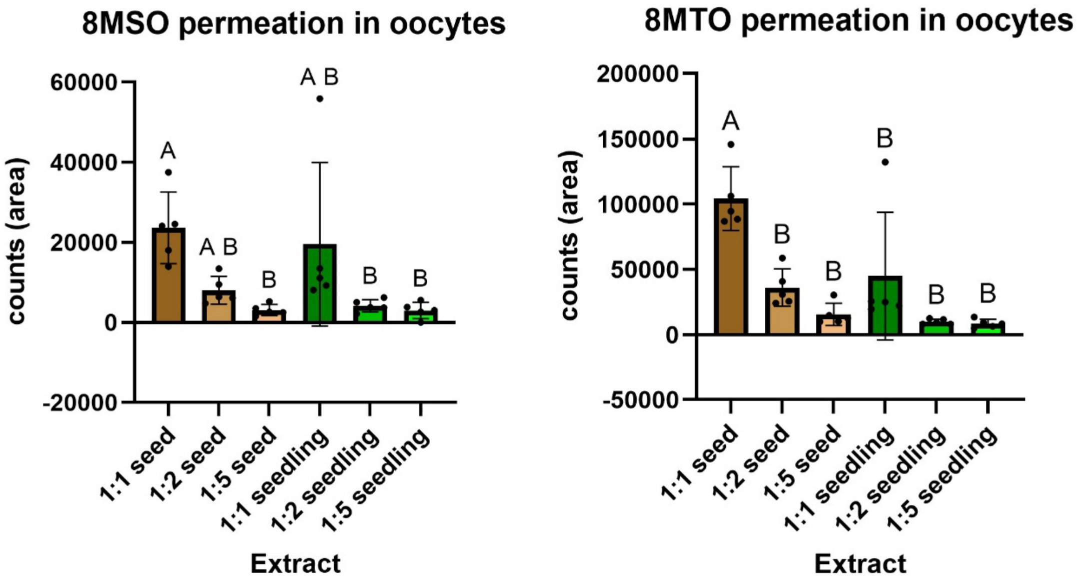

**Supplementary 1. Extract dilution alleviates membrane permeation.** Detection of the plant-derived glucohirsutin (8mso) and 8-Methylthiooctyl glucosinolate (8mto) in control oocytes exposed to the different dilutions of seed and seedling extracts. Oocytes were incubated in the different dilutions in Kulori media at pH 5 for 1 hour. Bars show the signal averages of 5 pools of 3 oocytes (n = 5) measured with multiple reaction monitoring MRM, and error bars represent the standard deviation. Letters represent the statistically different groups for a Tukey Multiple comparison test preceded by One-way ANOVA. Sig. p-value<0.05.

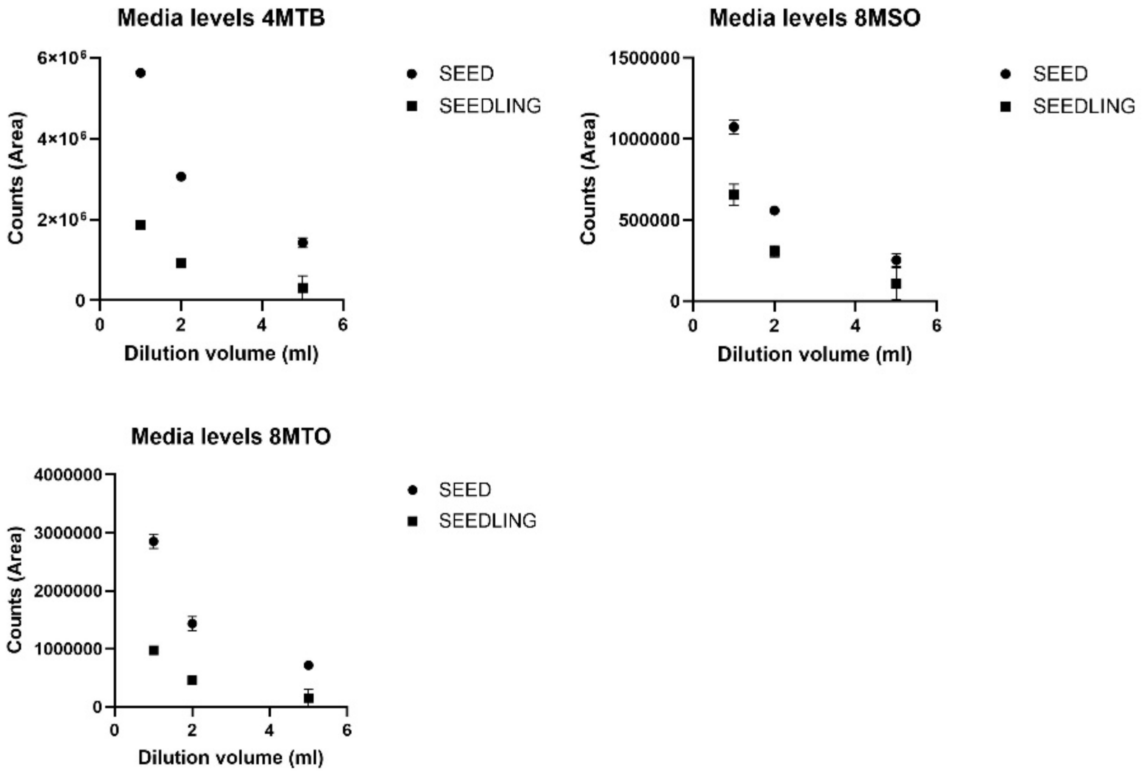

**Supplementary 2. Seed extract presents more of the major glucosinolates 4mtb, 8mto, and 8mso.** The plant-derived 4mtb, 8mto, and 8mso glucosinolates were detected in the different dilutions of seed and seedling extracts. 3µl of each dilution were measured twice.

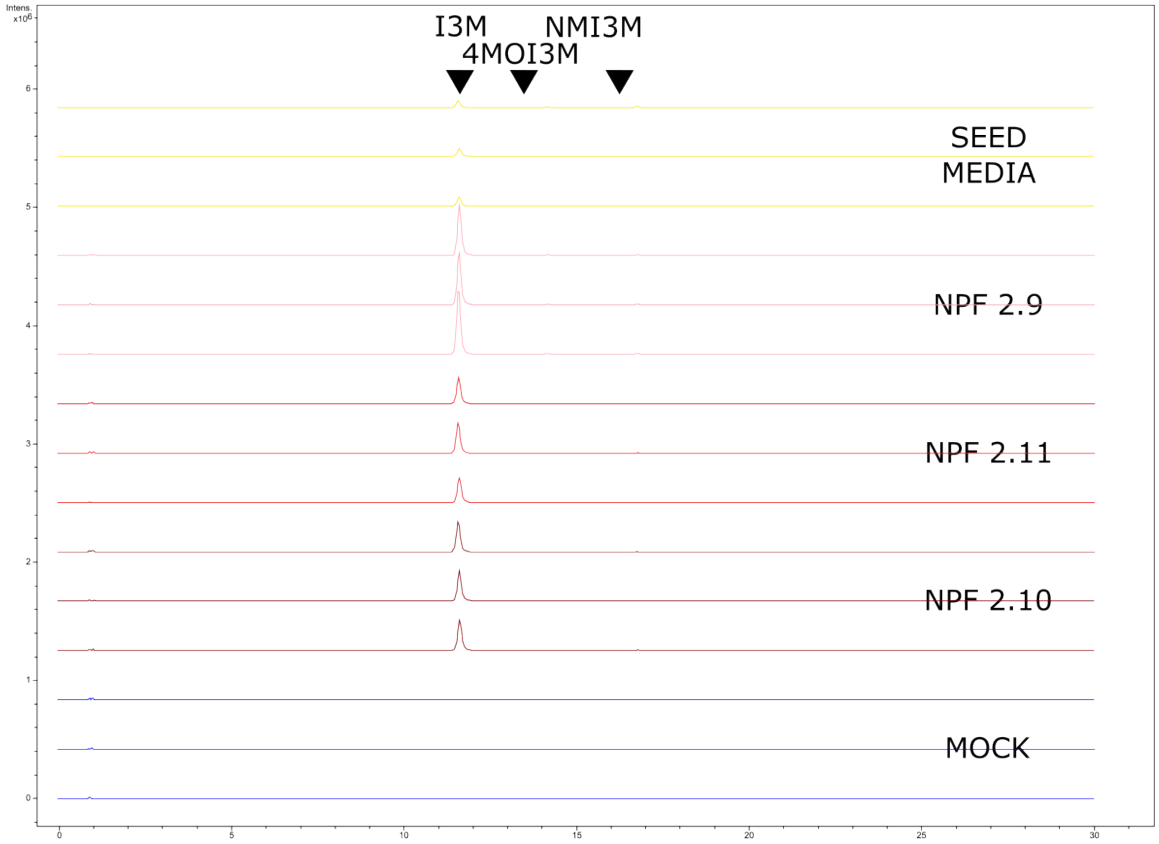

**Supplementary 3. The method reflects NPF2.9's preference for indole-glucosinolates.** Presence of indole-glucosinolates glucobrassicin (I3M), 4-methoxyglucobrassicin (4MOI3M), and neoglucobrassicin (NMOI3M) in oocytes exposed to 1:100 seed media. NPF2.10-expressing oocytes in brown, NPF2.11-expressing oocytes in red, and NPF2.9-expressing oocytes in pink. Control oocytes in blue and seed media in yellow. Oocytes were assayed at pH 5 for 1 hour; replicates consisted of five oocytes each, and media samples consisted of 5µl each. Combined extracted ion chromatogram for I3M (447.0537 +/- 0.005 m/z) and 4MOI3M and NMOI3M (477.0643 +/- 0.005 m/z).

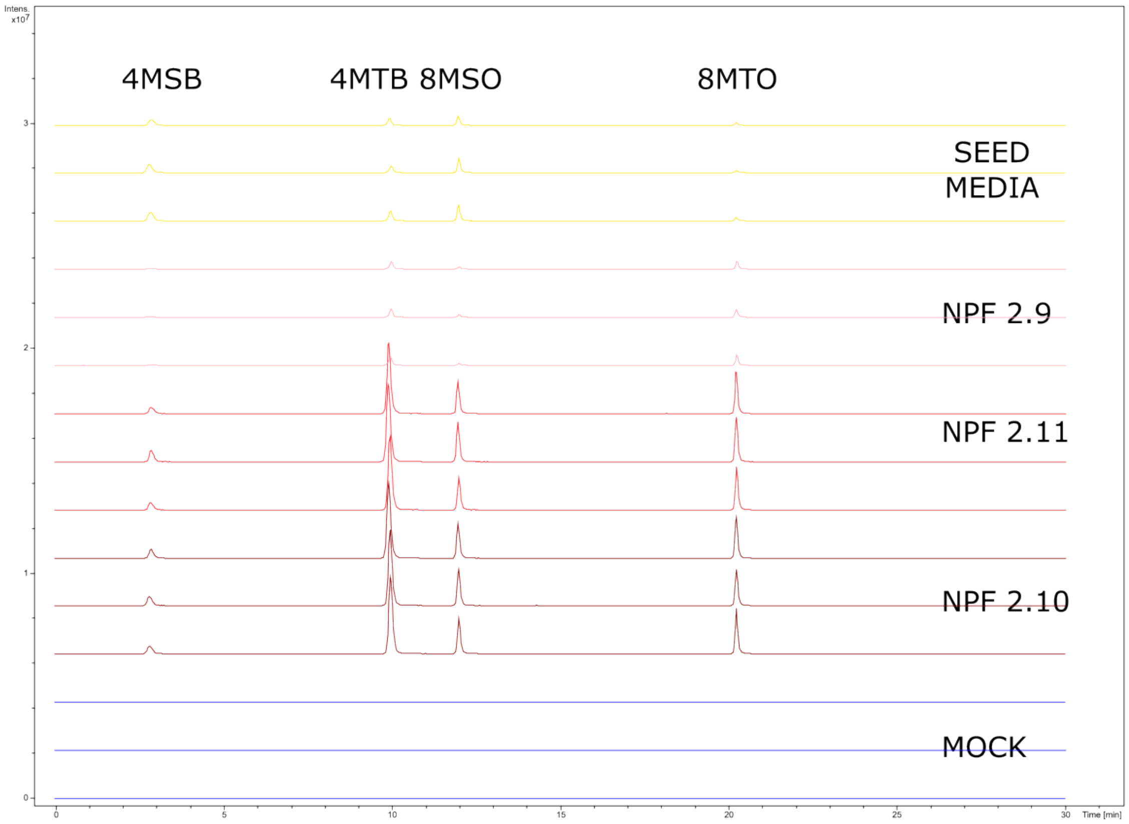

**Supplementary 4. NPF2.9 cannot accumulate some aliphatic glucosinolates over media levels.** Presence of aliphatic glucosinolates glucoerucin (4mtb), glucoraphanin (4msb), 8-methylthiooctyl glucosinolate (8mto) glucohirsutin (8mso) in oocytes exposed to 1:100 seed media NPF2.10-expressing oocytes in brown, NPF2.11-expressing oocytes in red, and NPF2.9-expressing oocytes in pink. Oocytes were assayed at pH 5 for 1 hour; replicates consisted of five oocytes each, and media samples consisted of 5µl each. Combined extracted ion chromatogram for 4msb (436.0411 +/- 0.005 m/z), 4mtb (420.0462 +/- 0.005 m/z), 8mso (492.1037 +/- 0.005 m/z) and 8mto (476.1088 +/- 0.005 m/z).

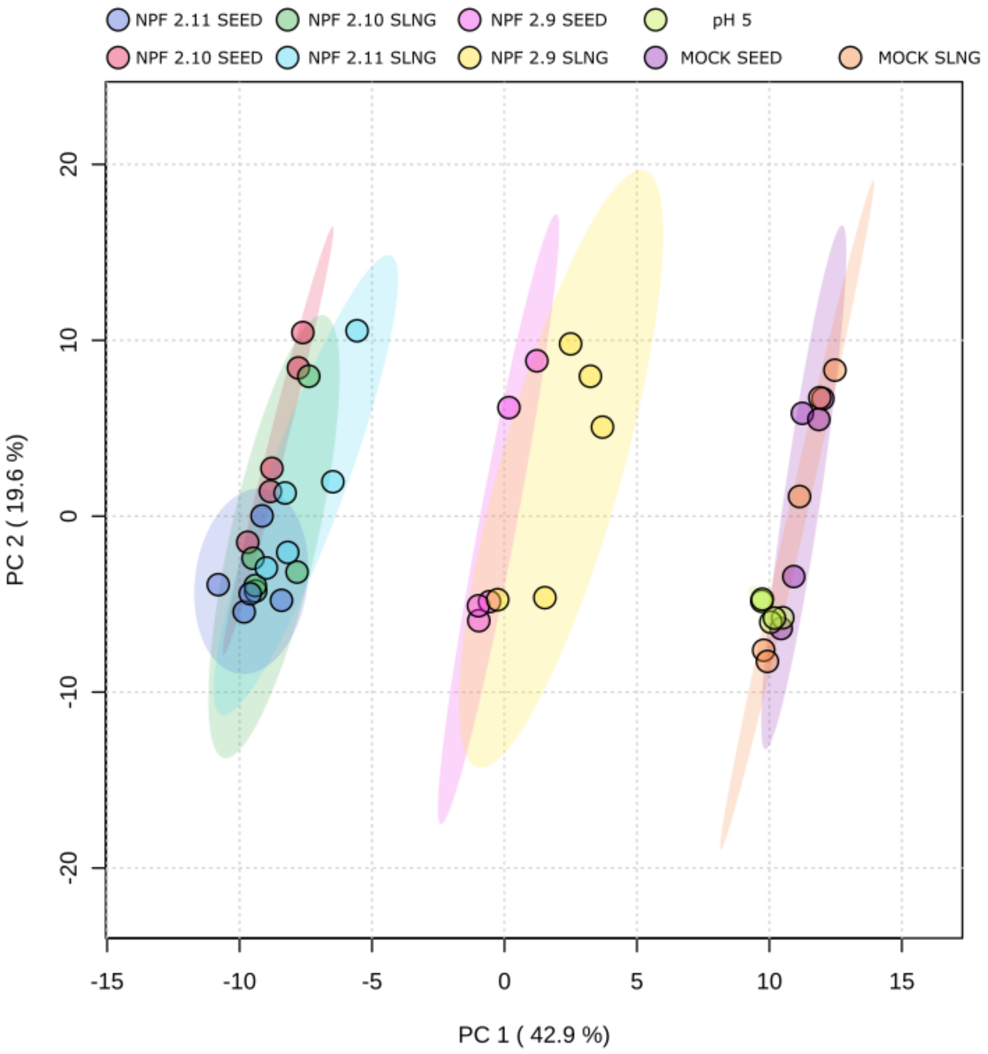

**Supplementary 5. The combination of the expressed transporter and the extract used defines the chemical composition of the assayed oocytes.** Principal Component Analysis (PCA) of the untargeted LCMS data obtained when assaying GTR-expressing oocytes with either a 1:100 dilution of seed (SEED) or a 1:100 dilution of seedling extract (SLNG). pH5 oocytes have been assayed with pH 5 empty media. Features displaying a relative standard deviation of >30% in the QC were filtered. Features were logarithmically transformed (Log.10) and Pareto scaled.

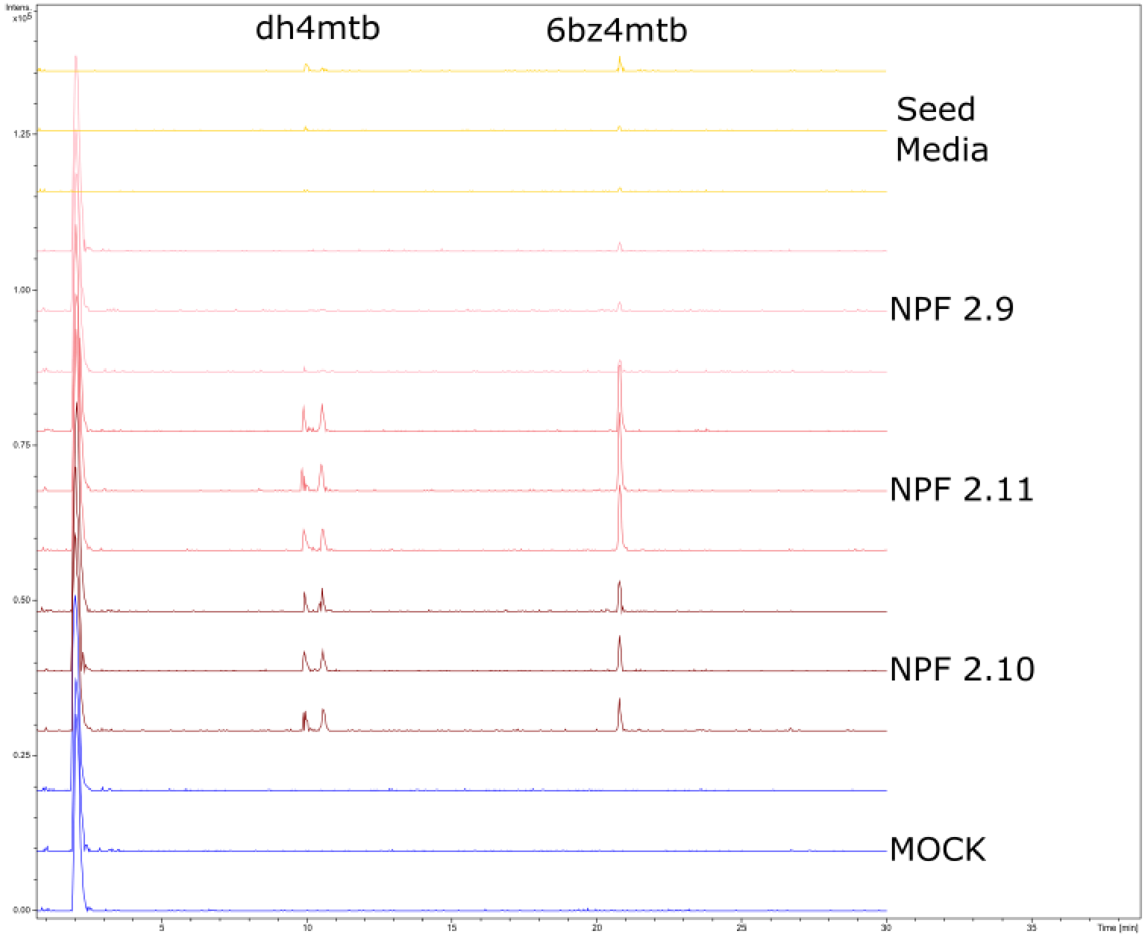

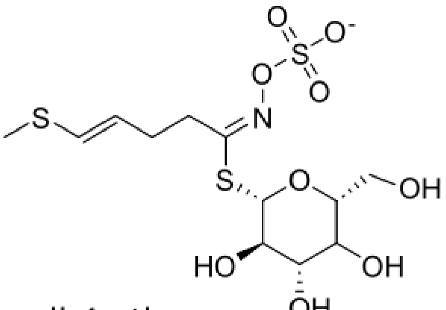

dh4mtb

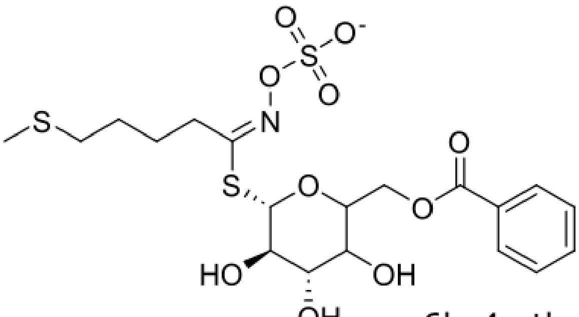

6bz4mtb

**Supplementary 6. NPF2.11 displays a higher accumulation of benzoylated glucosinolate 6bz4mtb, and slightly higher accumulation of glucoraphasatin.** Presence of 6-O-benzoyloxy-glucoerucin (6bz4mtb) and glucoraphasatin (dh4mtb) in oocytes exposed to 1:100 seed media. NPF2.10-expressing oocytes in brown, NPF2.11-expressing oocytes in red, and NPF2.9-expressing oocytes in pink. Oocytes were assayed at pH 5 for 1 hour; replicates consisted of five oocytes each, and media samples consisted of 5µl each. Combined extracted ion chromatogram for 6bz4mtb (524.0724 +/- 0.005 m/z) and glucoraphasatin (418.0305 +/- 0.005 m/z).

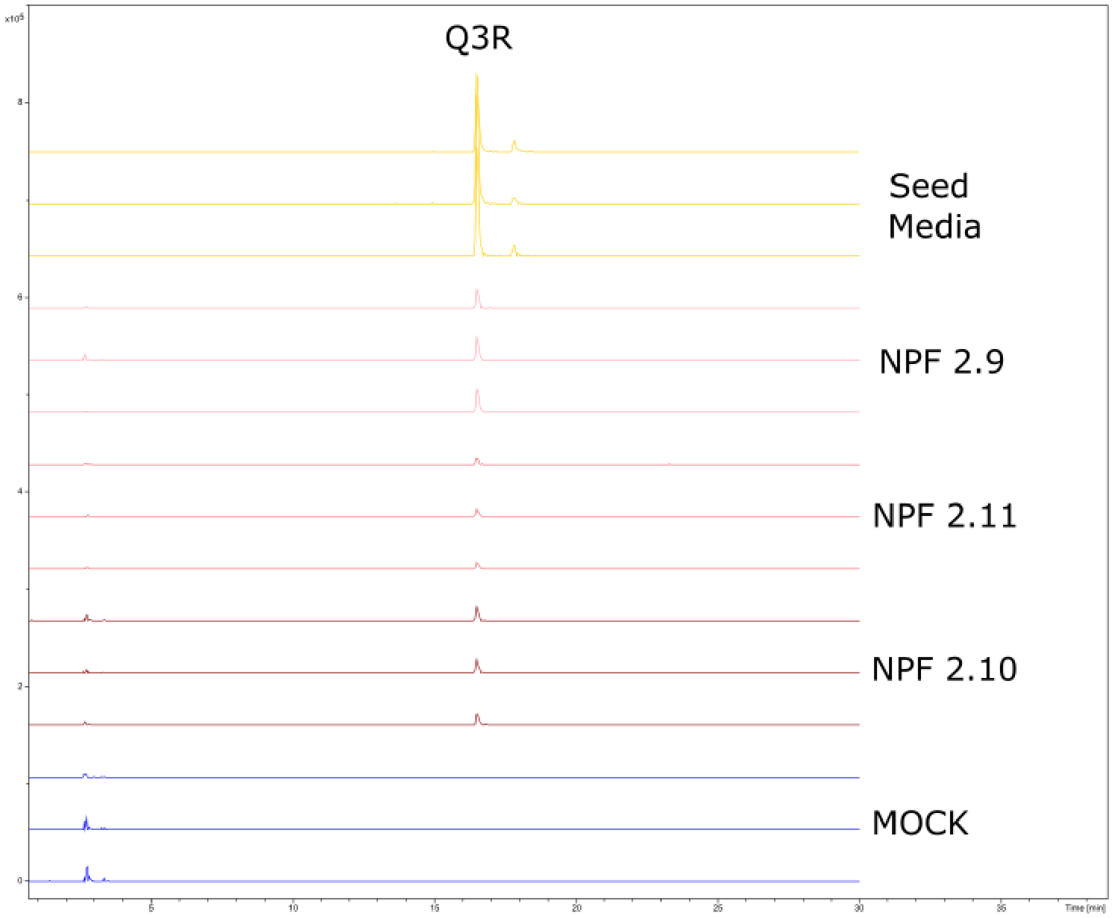

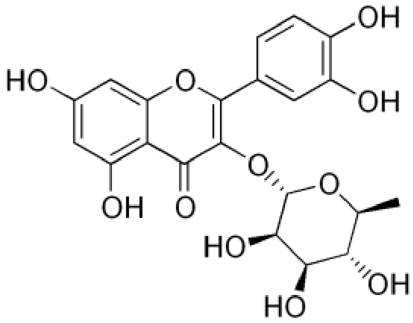

**Supplementary 7. The GTRs display an uptake of non-glucosinolate phytochemicals, which accumulate in a different pattern across genes.** Presence of Quercitrin (Q3R) in oocytes exposed to 1:100 seed media. NPF2.10-expressing oocytes in brown, NPF2.11-expressing oocytes in red, and NPF2.9-expressing oocytes in pink. Control oocytes in blue. Oocytes were assayed in pH5 for 1 hour; replicates consisted of five oocytes each, and media samples consisted of 5µl each. Extracted ion chromatogram for Q3R (447.0933 +/- 0.005 m/z).

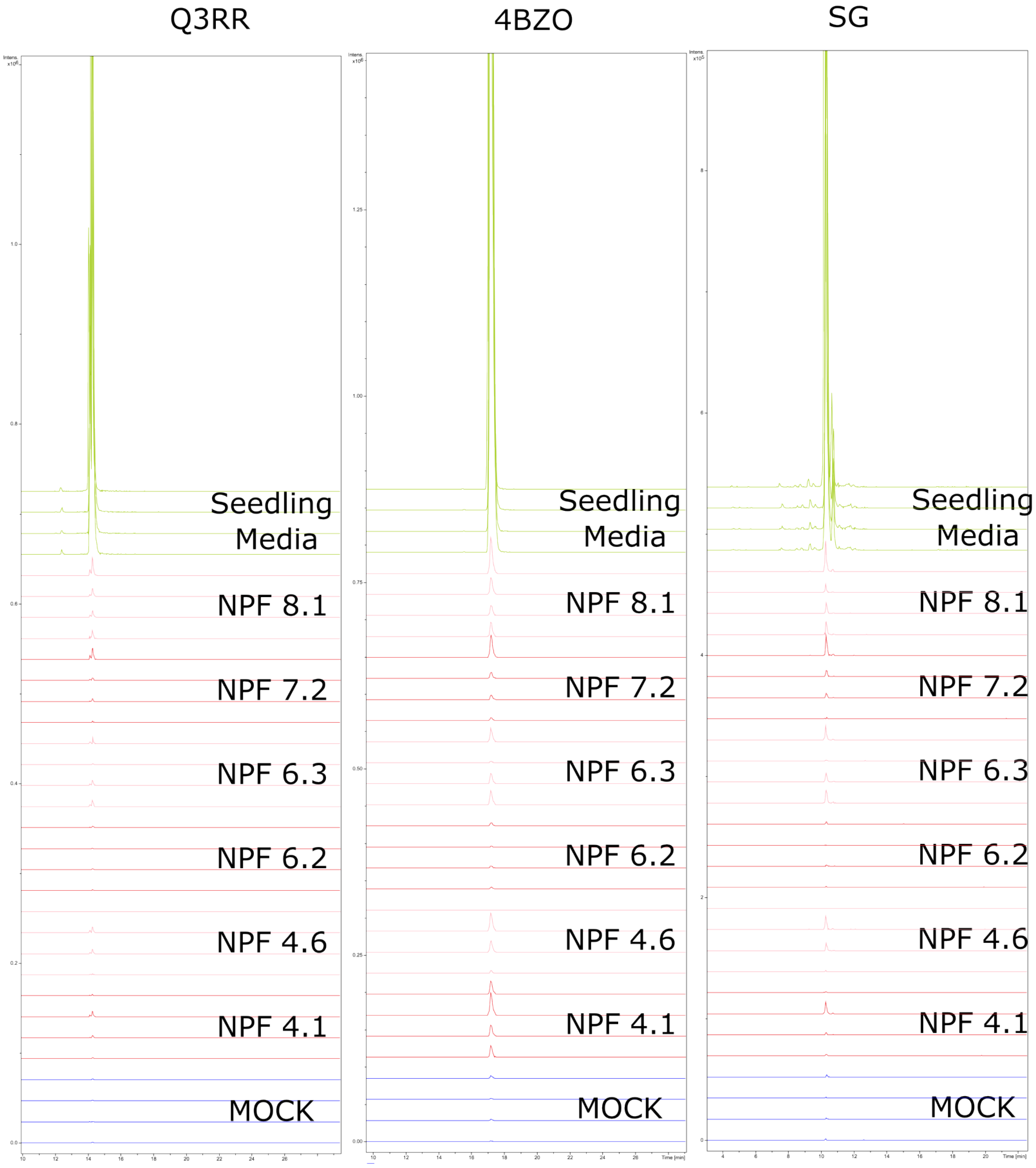

**Supplementary 8. The discarded transporters accumulate different classes of phytochemicals in orders of magnitude lower than media samples, in a heterogeneous manner.** Presence of Quercetin dirhamnoside (QRR), 4-benzoyloxybutyl glucosinolate (4bzo), and sinapoyl glucose (SG) in oocytes exposed to 1:20 seedling media. Transporter-expressing oocytes in red and pink, alternatively. Control oocytes in blue. Oocytes were assayed in pH5 for 1 hour; replicates consisted of five oocytes each, and media samples consisted of 5µl each. Extracted ion chromatogram for Q3R (593.1512 +/- 0.005 m/z). 4bzo (494.0796+/- 0.005 m/z), (494.0796+/- 0.005 m/z).

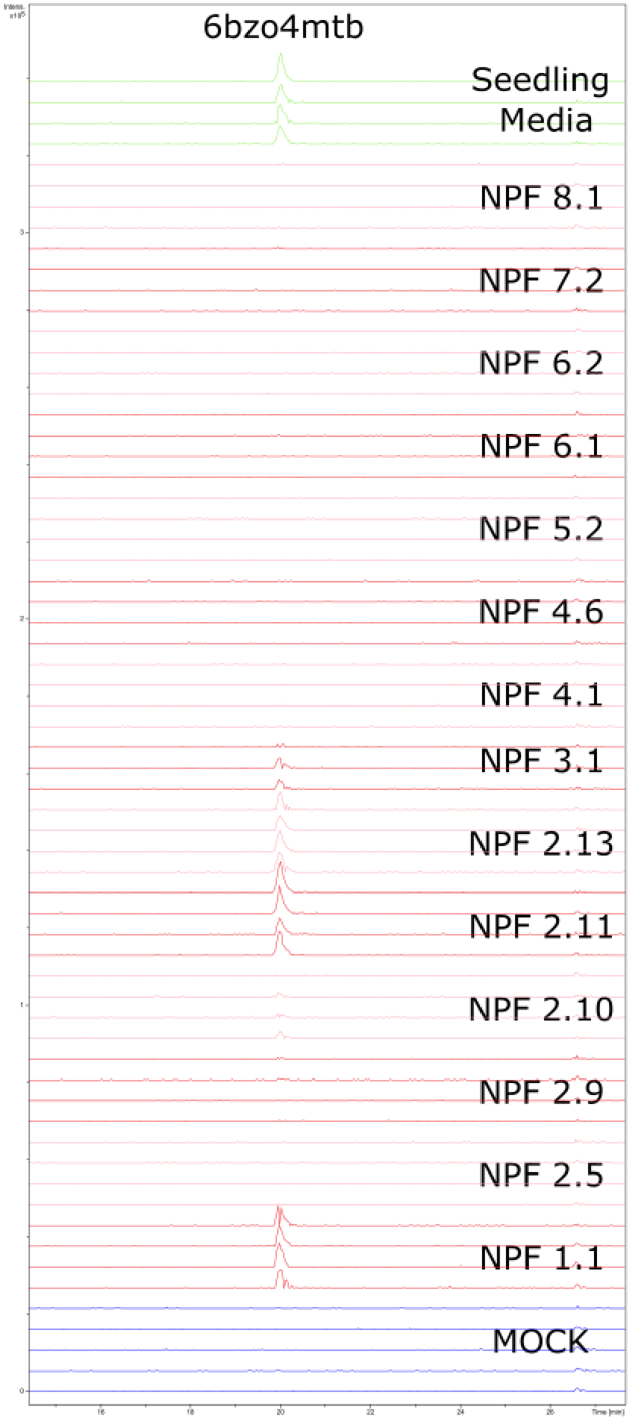

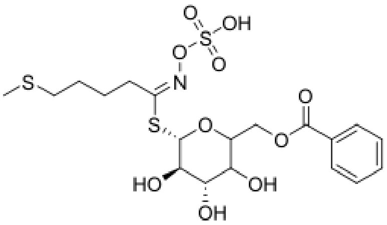

**Supplementary 9. Accumulation pattern of 6'bzo4mtb across 14 NPFs.** Presence of the annotated 6'-O-benzoyloxy-glucoerucin (6'bz4mtb) in oocytes exposed to 1:20 seedling media. Transporter-expressing oocytes in red and pink, alternatively. Control oocytes in blue. Oocytes were assayed in pH5 for 1 hour; replicates consisted of five oocytes each (n=4), and media samples consisted of 5µl each. Extracted ion chromatogram for 6'bzo4mtb (524.0724 +/- 0.005 m/z).

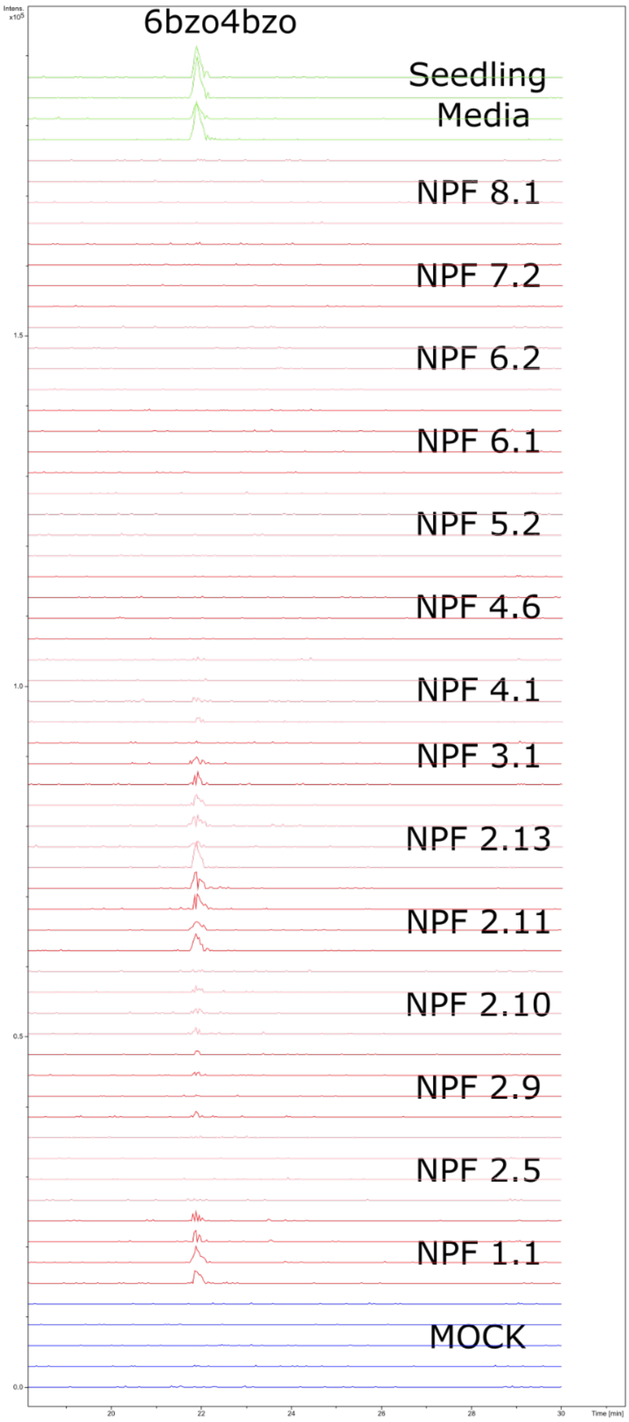

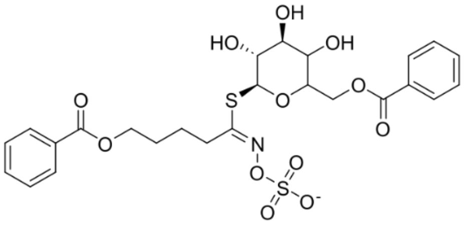

**Supplementary 10. Accumulation pattern of 6'bz4bzo across 14 NPFs.** Presence of 6'bz4bzo in oocytes exposed to 1:20 seedling media. Transporter-expressing oocytes in red and pink, alternatively. Control oocytes in blue. Oocytes were assayed at pH 5 for 1 hour; replicates consisted of five oocytes each, and media samples consisted of 5µl each. Extracted ion chromatogram for 6'bz4bzo(598.1058 +/- 0.005 m/z).

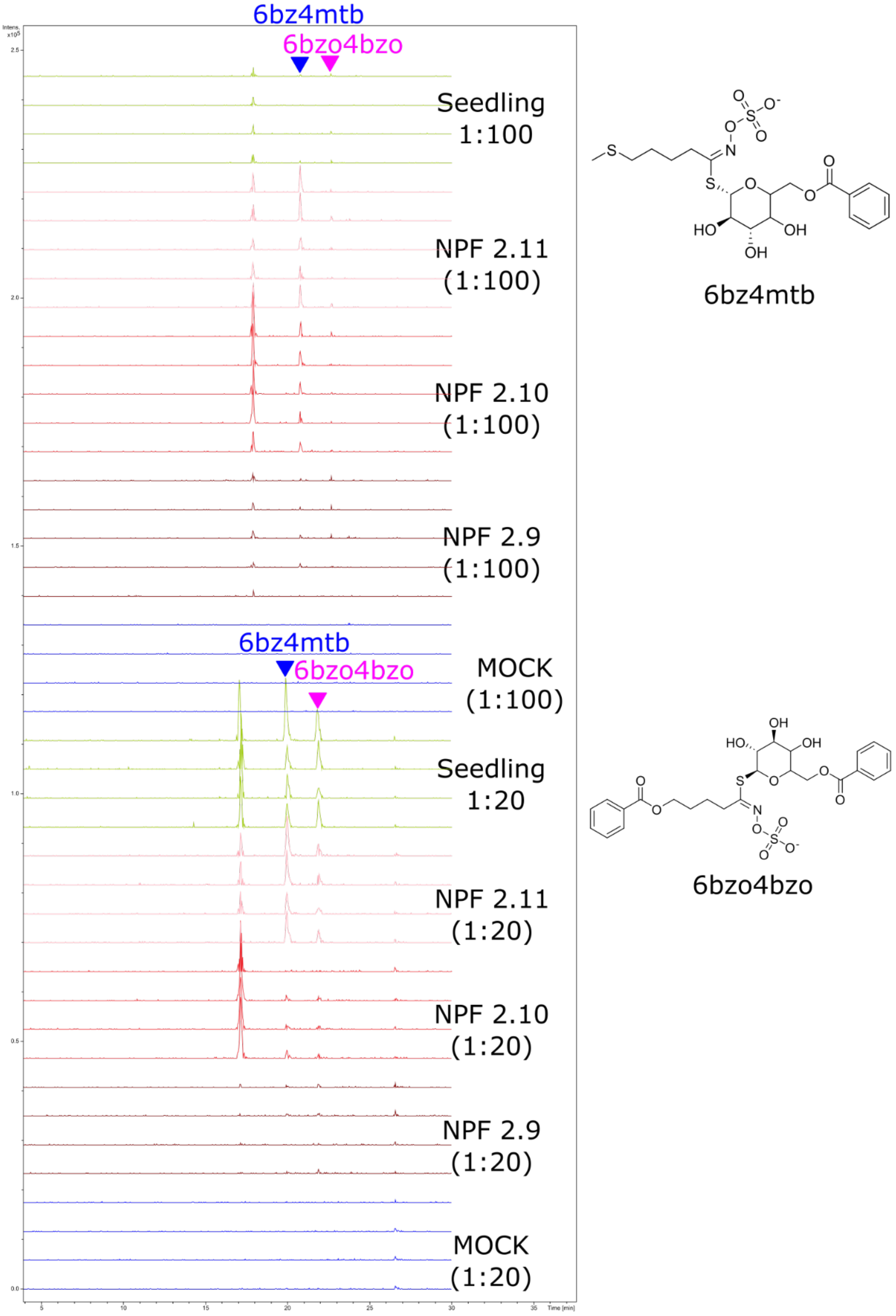

**Supplementary 11. The import of 6'bz4bzo is barely detectable from the hundredfold dilution of the extract.** Presence of 6'bz4bzo and 6'bz4mtb in oocytes exposed to 1:20 and 1:100 seedling media. NPF2.9-expressing oocytes in brown, NPF2.10-expressing oocytes in red, and NPF2.11-expressing oocytes in pink. Control oocytes in blue, media in green. Oocytes were assayed at pH 5 for 1 hour; replicates consisted of five oocytes each, and media samples consisted of 5µl each. Extracted ion chromatogram for 6'bz4bzo (598.1058 +/- 0.005 m/z) and 6'bzo4mtb (524.0724 +/- 0.005 m/z).

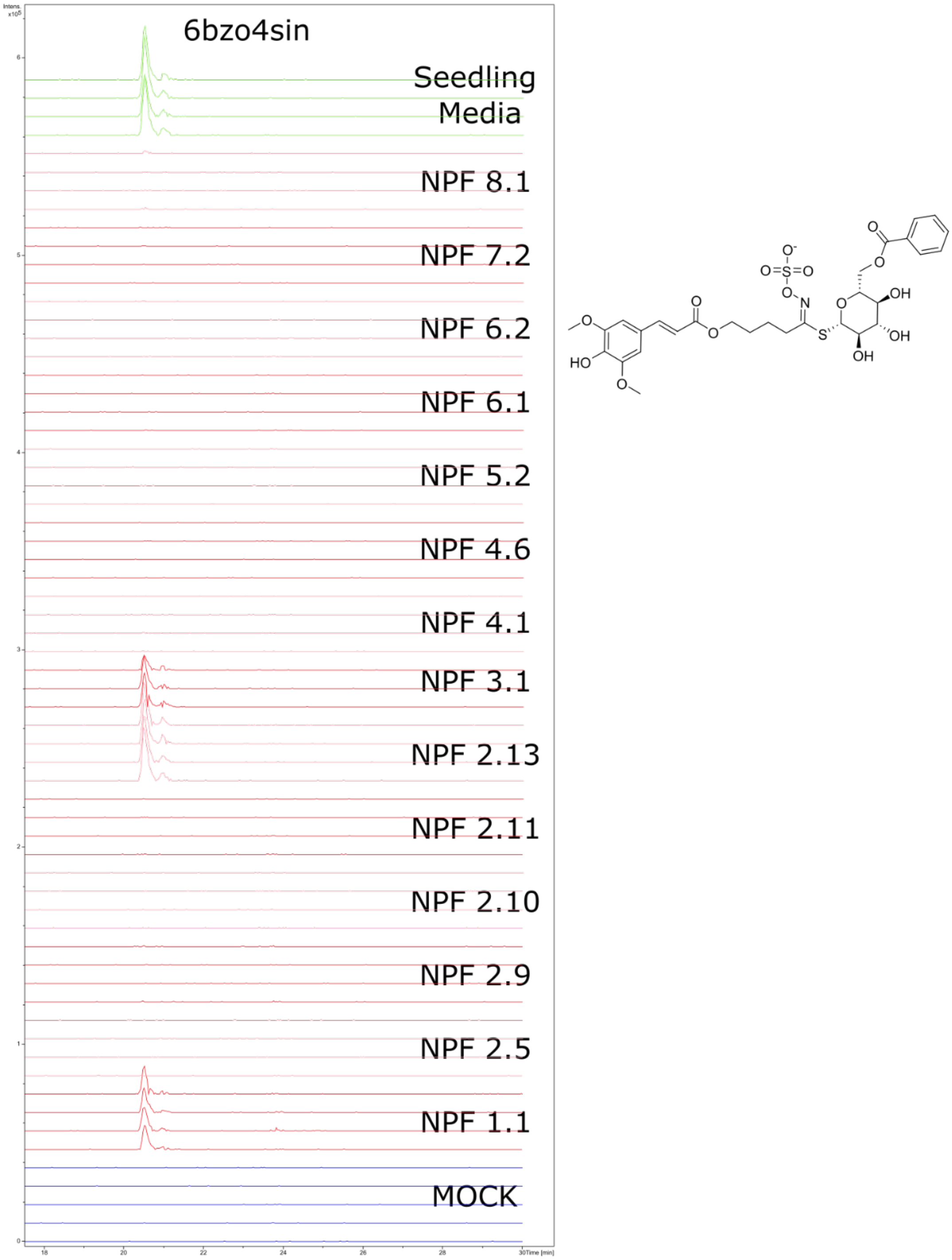

**Supplementary 12. Accumulation pattern of tentatively annotated 6'bz4sin across 14 NPFs.** Presence of annotated 6′-O-benzoyloxy-4-benzoyloxybutylglucosinolate (6'bz4bzo), in oocytes exposed to 1:20 seedling media. Transporter-expressing oocytes in red and pink, alternatively. Control oocytes in blue. Media in green. Oocytes were assayed in pH5 for 1 hour; replicates consisted of five oocytes each, and media samples consisted of 5µl each. Extracted ion chromatogram for 6'bz4sin (700.1375 +/- 0.005 m/z).

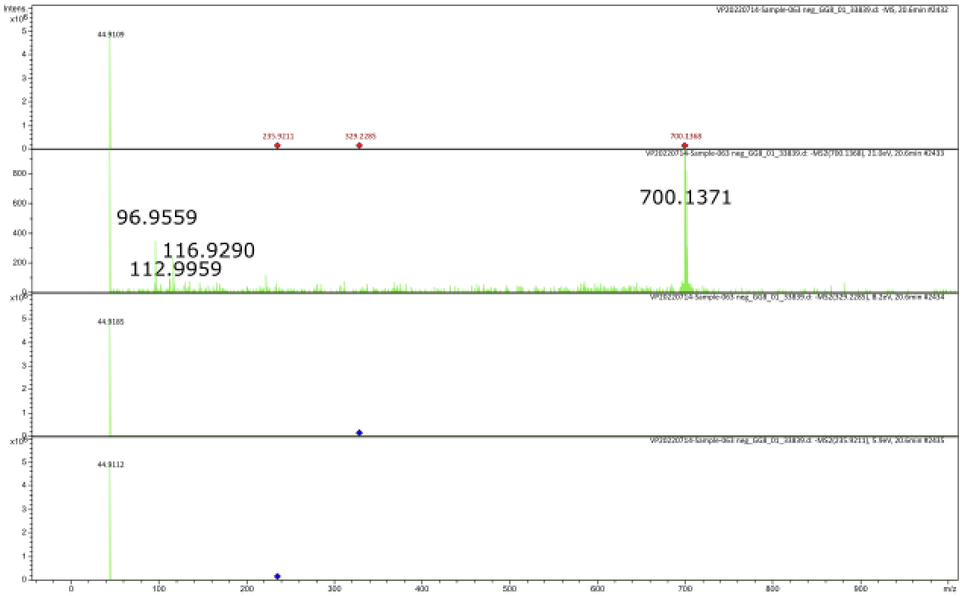

**Supplementary 13. MS2 pattern of tentatively annotated 6’bz4sin.** MS2 spectra of precursor ion (700.1371) attributed to 6′-O-benzoyloxy-4-benzoyloxybutylglucosinolate (6’bz4bzo). Second window from the top.

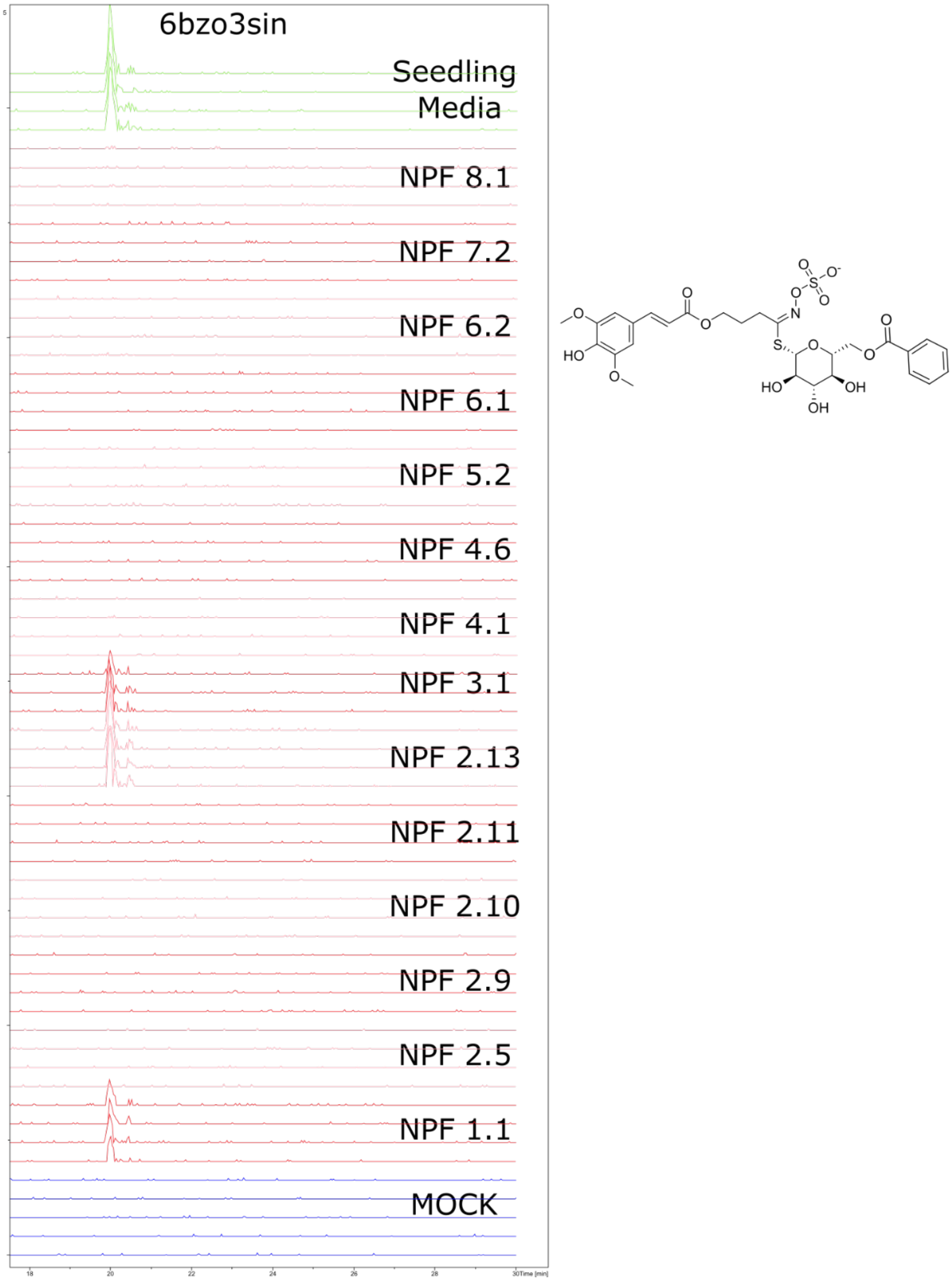

**Supplementary 14. Accumulation pattern of tentatively annotated 6’bz3sin across 14 NPFs.** Presence of annotated 6′-O-benzoyloxy-3-sinapoyloxypropylglucosinolate (6’bz3sin) in oocytes exposed to 1:20 seedling media. Transporter-expressing oocytes in red and pink, alternatively. Control oocytes in blue. Media in green. Oocytes were assayed at pH 5 for 1 hour; replicates consisted of five oocytes each, and media samples consisted of 5µl each. Extracted ion chromatogram for 6’bz3sin (686.1219+/- 0.005 m/z).

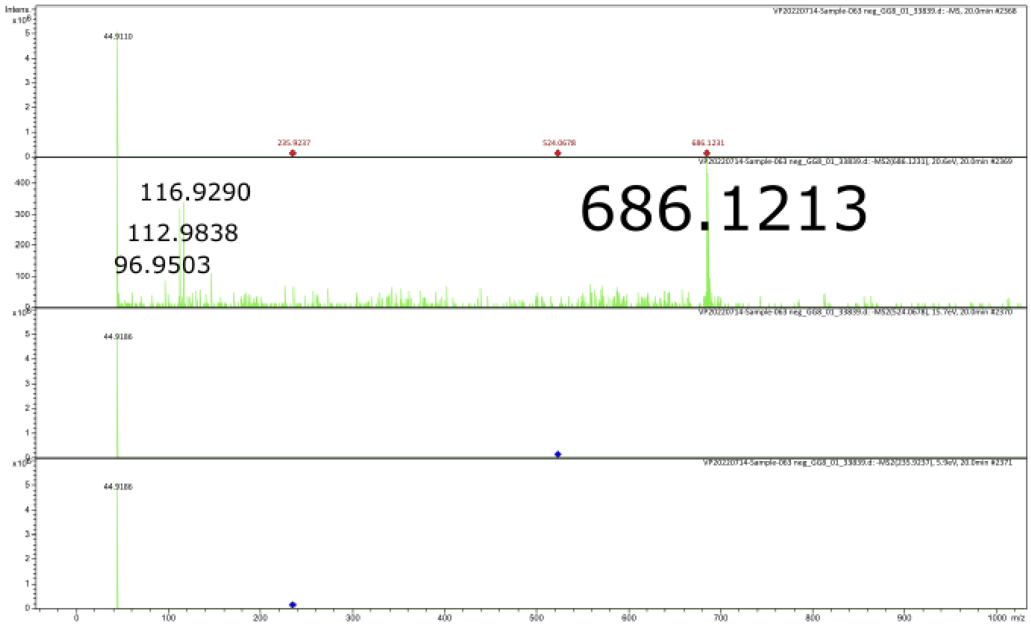

**Supplementary 15. MS2 pattern of tentatively annotated 6'bz3sin.** MS2 spectra of precursor ion (686.1213) attributed to 6′-O-benzoyloxy-3-sinapoyloxypropylglucosinolate (6'bz3sin). Second window from the top.

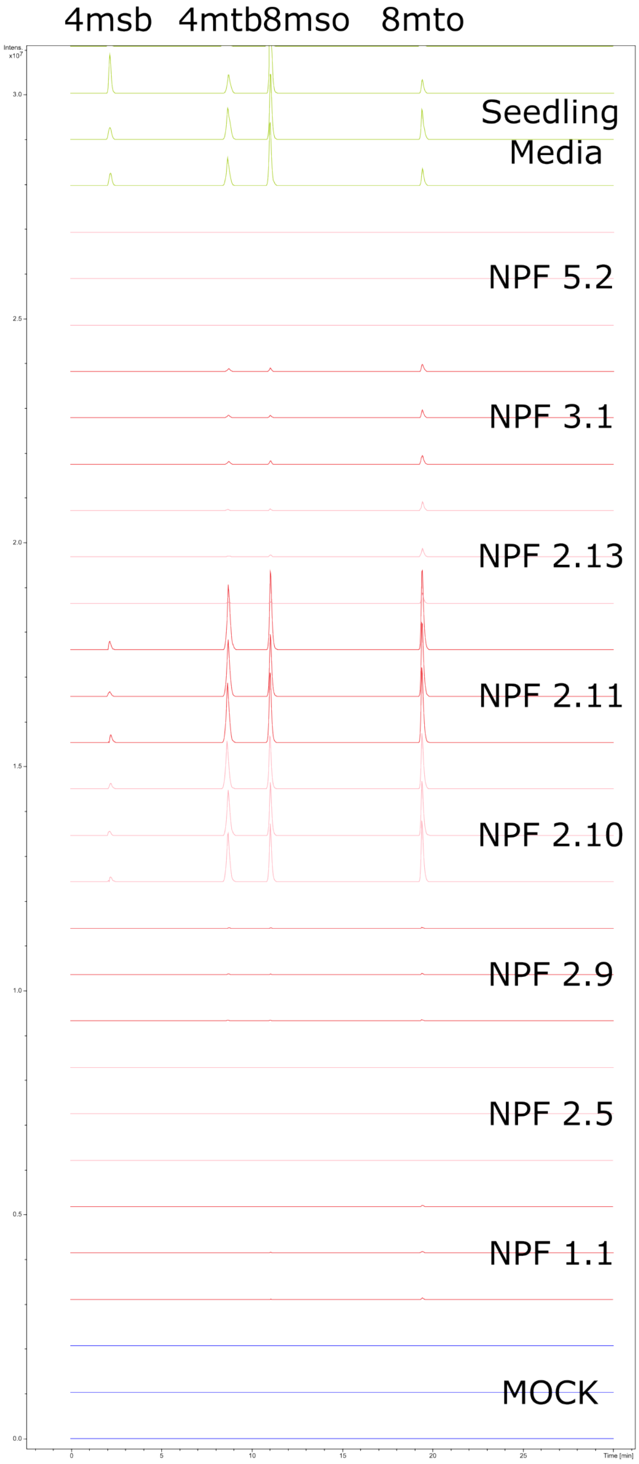

**Supplementary 16. Accumulation pattern of aliphatic glucosinolates across selected NPFs.** Presence of aliphatic glucosinolates glucoerucin (4mtb), glucoraphanin (4msb), 8-methylthiooctyl glucosinolate (8mto), glucohirsutin (8mso) in oocytes exposed to 1:20 seed media. Transporter-expressing oocytes in red and pink, alternatively. Control oocytes in blue. Media in green. Oocytes were assayed at pH 5 for 1 hour; replicates consisted of five oocytes each, and media samples consisted of 5μl each. Combined extracted ion chromatogram for 4msb (436.0411 +/- 0.005 m/z), 4mtb (420.0462 +/- 0.005 m/z), 8mso (492.1037 +/- 0.005 m/z) and 8mto (476.1088 +/- 0.005 m/z).

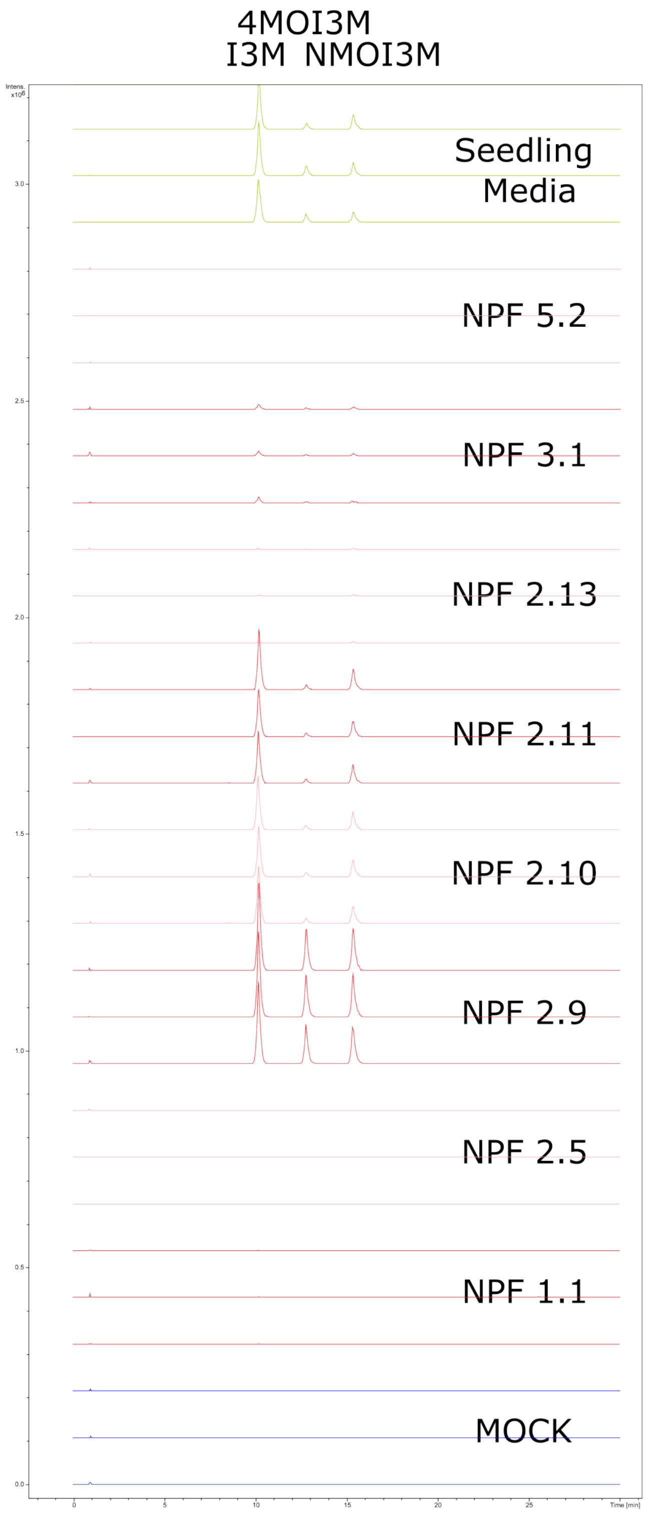

**Supplementary 17. Accumulation pattern of indole-glucosinolates across selected NPFs.** Presence of indole-glucosinolates glucobrassicin (I3M), 4-methoxyglucobrassicin (4MOI3M), and neoglucobrassicin (NMOI3M) in oocytes exposed to 1:20 seedling media. Oocytes were assayed at pH 5 for 1 hour; replicates consisted of five oocytes each, and media samples consisted of 5µl each. Transporter-expressing oocytes in red and pink, alternatively. Control oocytes in blue. Media in green. Combined extracted ion chromatogram for I3M (447.0537 +/- 0.005 m/z) and 4MOI3M and NMOI3M (477.0643 +/- 0.005 m/z).

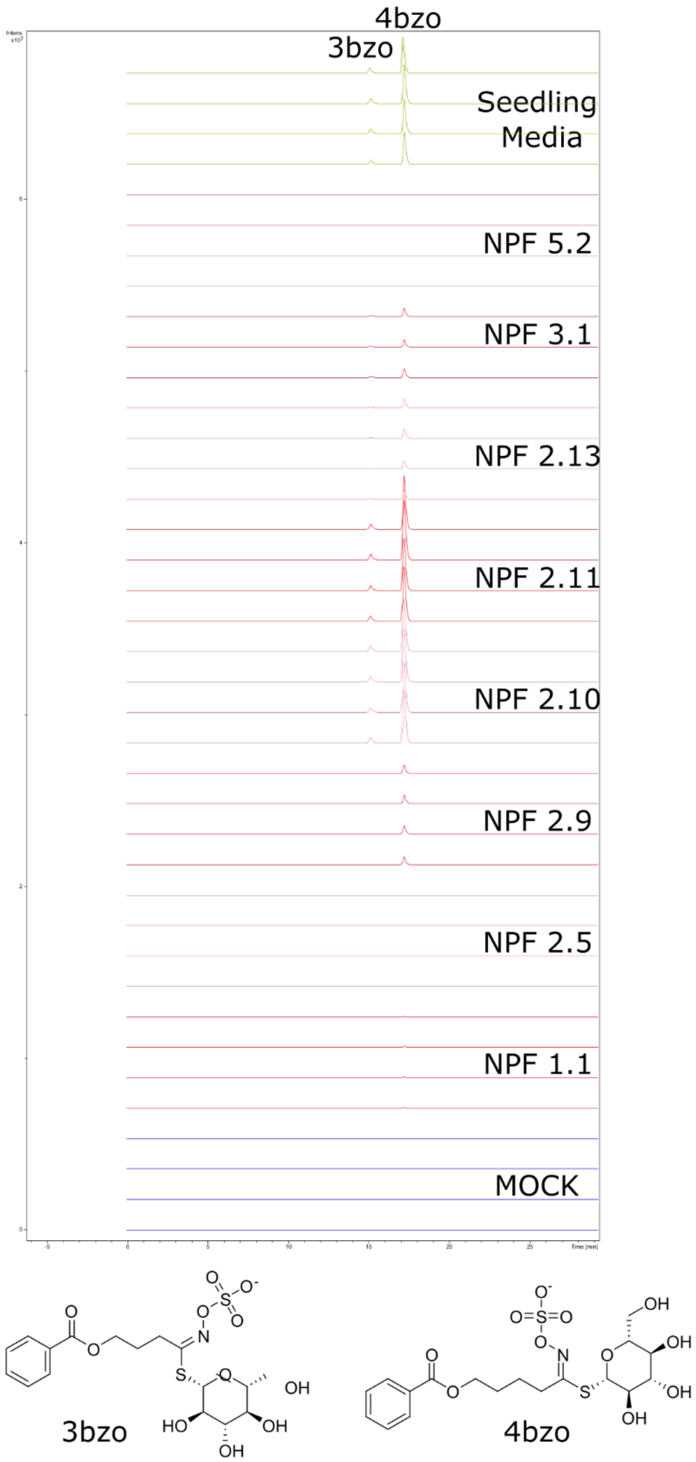

**Supplementary 18. Accumulation pattern of 3-benzoyloxypropyl and 4-benzoyloxybutyl glucosinolates (3bzo and 4bzo) across selected NPFs.** Presence of 3bzo and 4bzo in oocytes exposed to 1:20 seedling media. Oocytes were assayed at pH 5 for 1 hour; replicates consisted of five oocytes each, and media samples consisted of 5µl each. Transporter-expressing oocytes in red and pink, alternatively. Control oocytes in blue. Media in green. Combined extracted ion chromatogram for 3bzo (480.0640; +/- 0.005 m/z) and 4bzo (494.0796+/- 0.005 m/z).

Coniferin

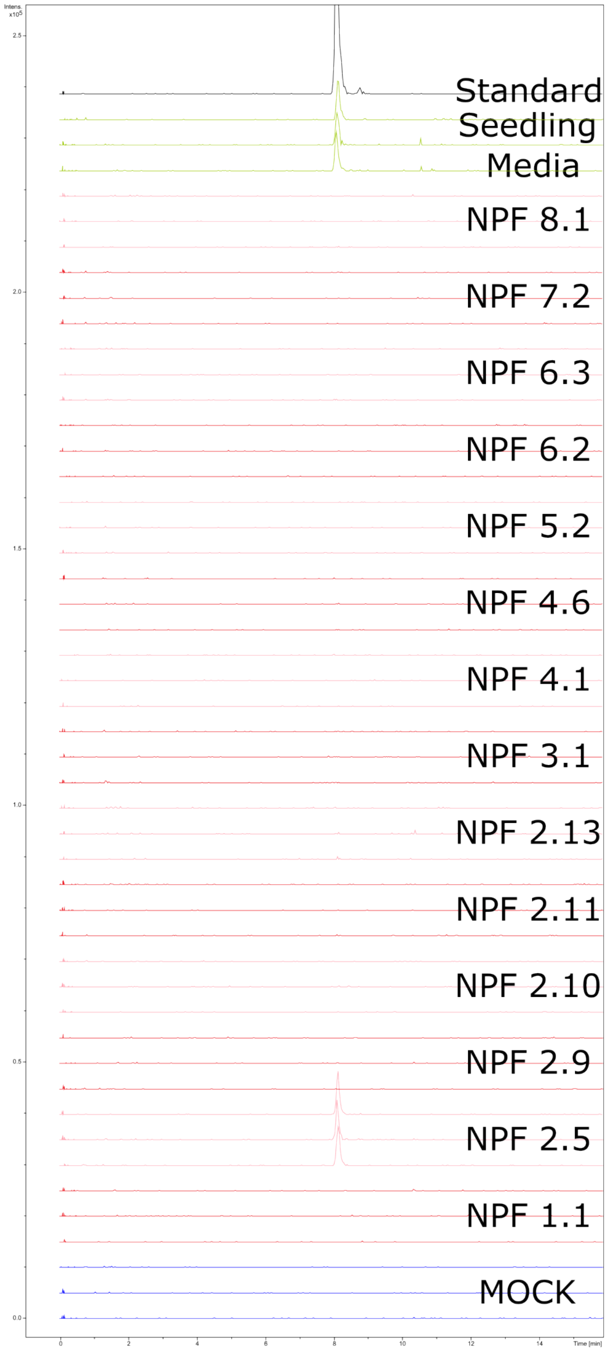

**Supplementary 19. NPF2.5 accumulates the monolignol glucoside coniferin.** Presence of coniferin in oocytes exposed to 1:20 seedling media. Transporter-expressing oocytes in red and pink, alternatively. Control oocytes in blue. Media in green. 10 µM chemical standard in black. Oocytes were assayed at pH 5 for 1 hour; replicates consisted of five oocytes each, and media samples consisted of 5µl each. Extracted ion chromatogram for the formic acid adduct of coniferin (387.1297 +/- 0.005 m/z).

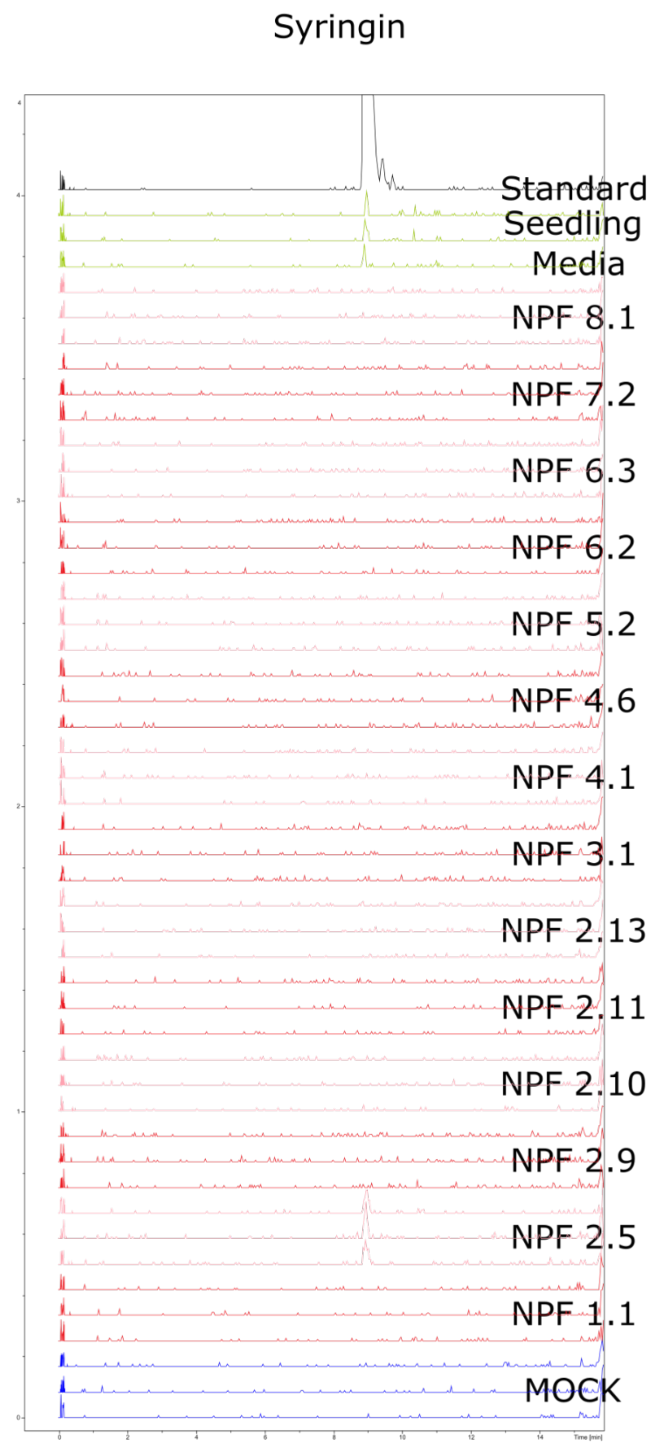

**Supplementary 20. NPF2.5 accumulates monolignol glucoside syringin.** Presence of syringin in oocytes exposed to 1:20 seedling media. Transporter-expressing oocytes in red and pink, alternatively. Control oocytes in blue. Media in green. 10 µM chemical standard in black. Oocytes were assayed at pH 5 for 1 hour; replicates consisted of five oocytes each, and media samples consisted of 5µl each. Extracted ion chromatogram for the in-source fragment of syringing corresponding to sinapyl alcohol (209.0819 +/- 0.005 m/z).

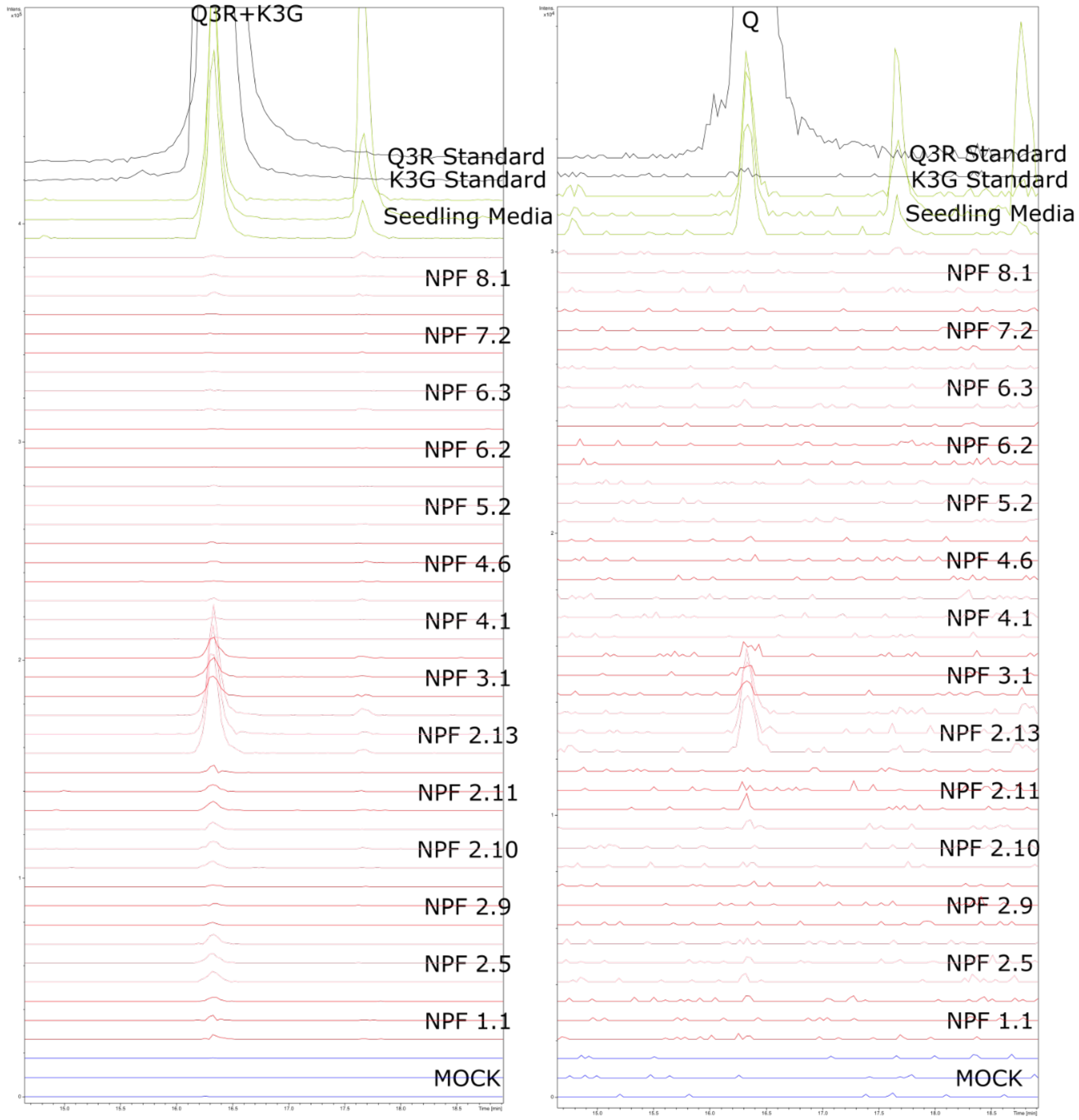

**Supplementary 21. Accumulation pattern of the flavonol quercitrin (Q3R) across 14 NPFs.** The first chromatogram corresponds to the ambiguous feature representing Q3R and K3G. The second chromatogram corresponds to the discriminating in-source fragment quercetin (Q). Transporter-expressing oocytes in red and pink, alternatively. Control oocytes in blue. Media in green. 10 µM chemical standards in black. Oocytes were assayed at pH 5 for 1 hour; replicates consisted of five oocytes each, and media samples consisted of 5µl each. Extracted ion chromatogram for Q3R and K3G (447.0933 +/- 0.005 m/z) and the in-source fragment Quercetin (Q) (301.0354 +/- 0.005 m/z).

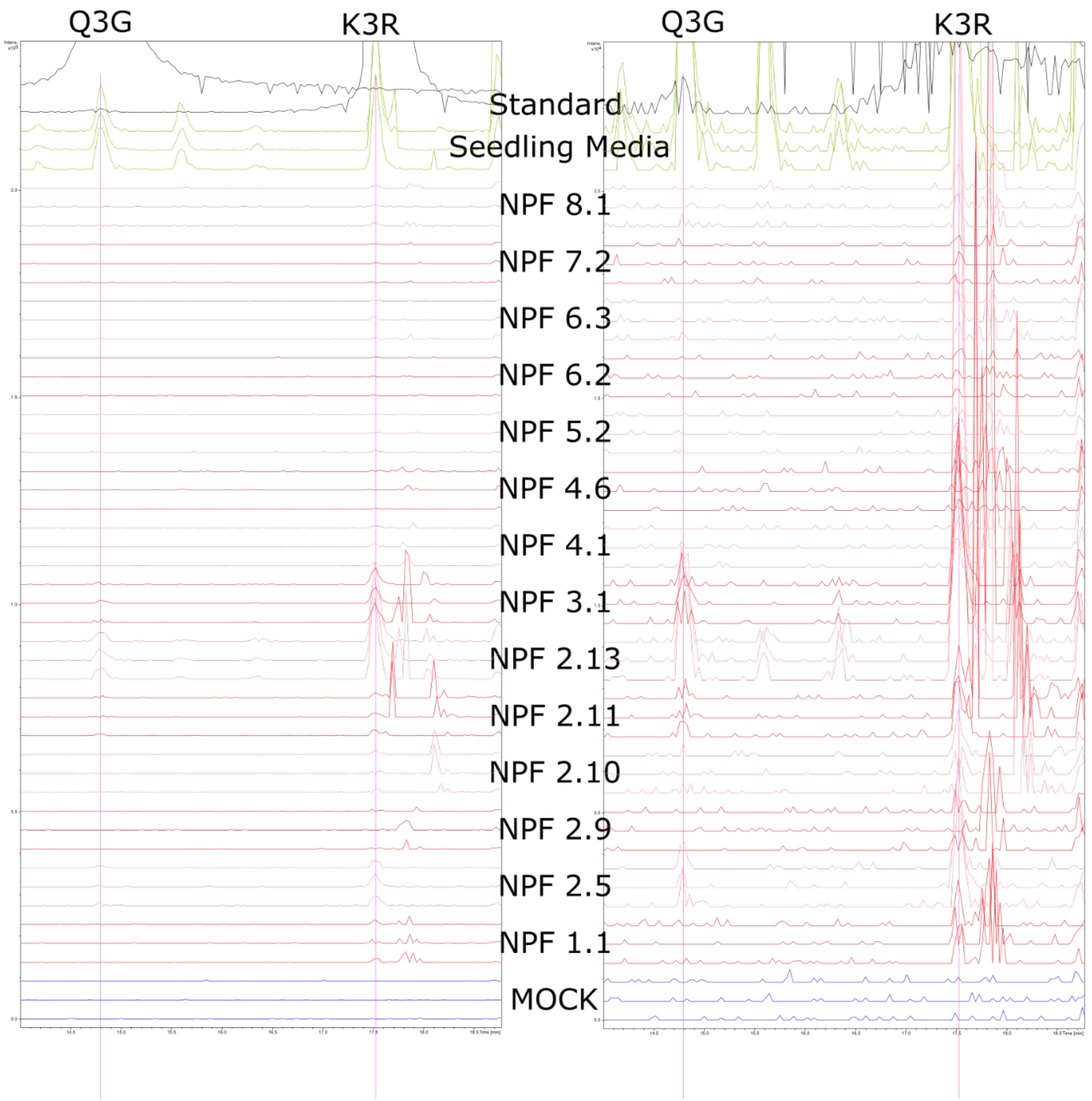

**Supplementary 22. Accumulation pattern of the flavonols isoquercetin (Q3G) and afzelin (k3R) across 14 NPFs.** Presence of K3R in oocytes exposed to 1:20 seedling media. Transporter-expressing oocytes in red and pink, alternatively. Control oocytes in blue. Media in green. 10 µM chemical standards in black. Oocytes were assayed at pH 5 for 1 hour; replicates consisted of five oocytes each, and media samples consisted of 5µl each. Combined Extracted ion chromatogram for K3R (431.0984 +/- 0.005 m/z) and Q3G (463.0882+/- 0.005 m/z).

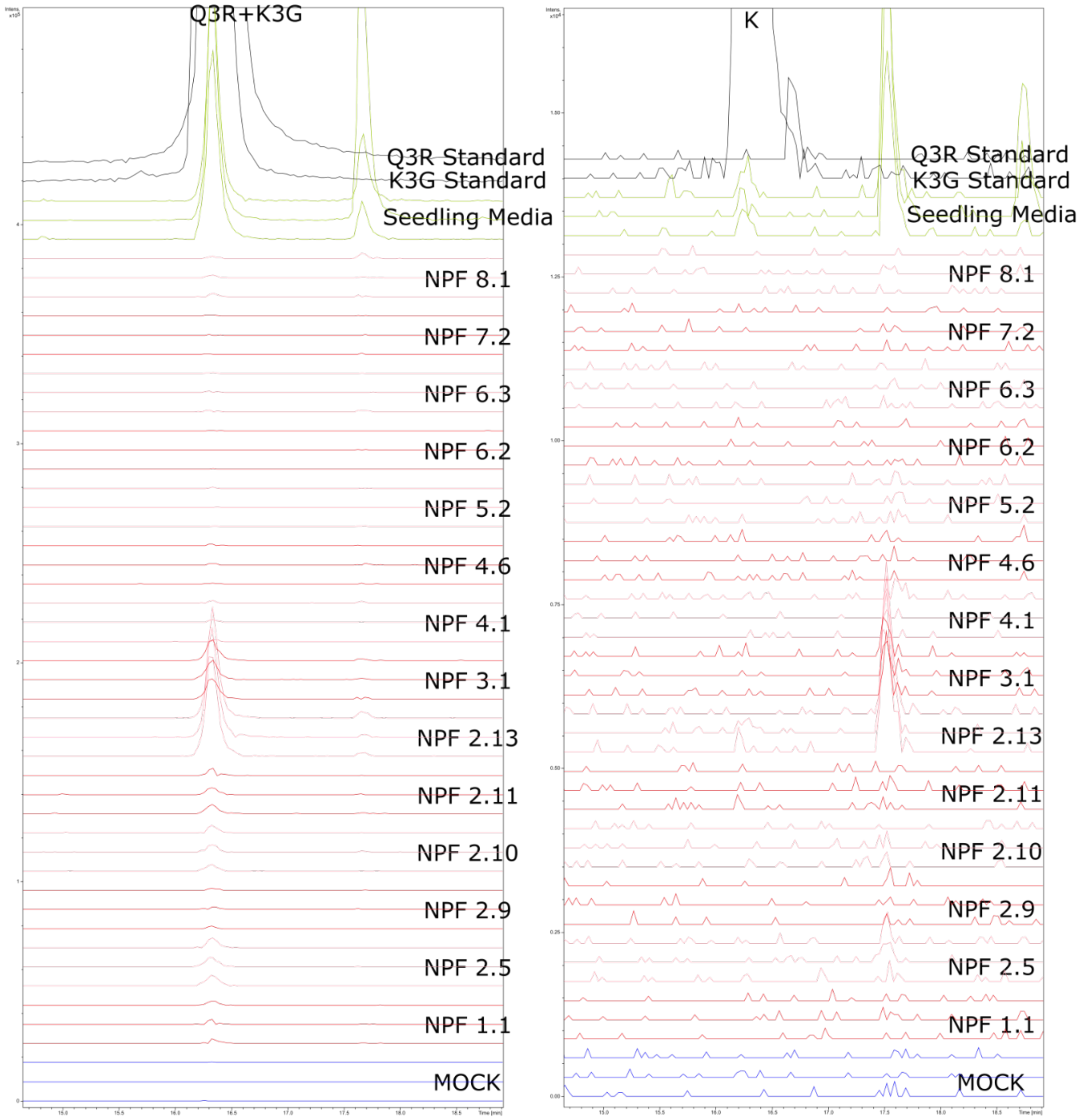

**Supplementary 23. Accumulation pattern of the flavonol astragalin (K3G) across 14 NPFs.** The first chromatogram corresponds to the ambiguous feature representing Q3R and K3G. The second chromatogram corresponds to the discriminating in-source fragment kaempferol (K). Presence of K3G in oocytes exposed to 1:20 seedling media. The presence of K3G cannot be detected from the discriminating in-source fragment due to the low signal. Transporter-expressing oocytes in red and pink, alternatively. Control oocytes in blue. Media in green. 10 µM chemical standards in black. Oocytes were assayed at pH 5 for 1 hour; replicates consisted of five oocytes each, and media samples consisted of 5µl each. Extracted ion chromatogram for Q3R and K3G (447.0933 +/- 0.005 m/z) and the in-source fragment Kaempferol (K) (285.0405 +/- 0.005 m/z) .

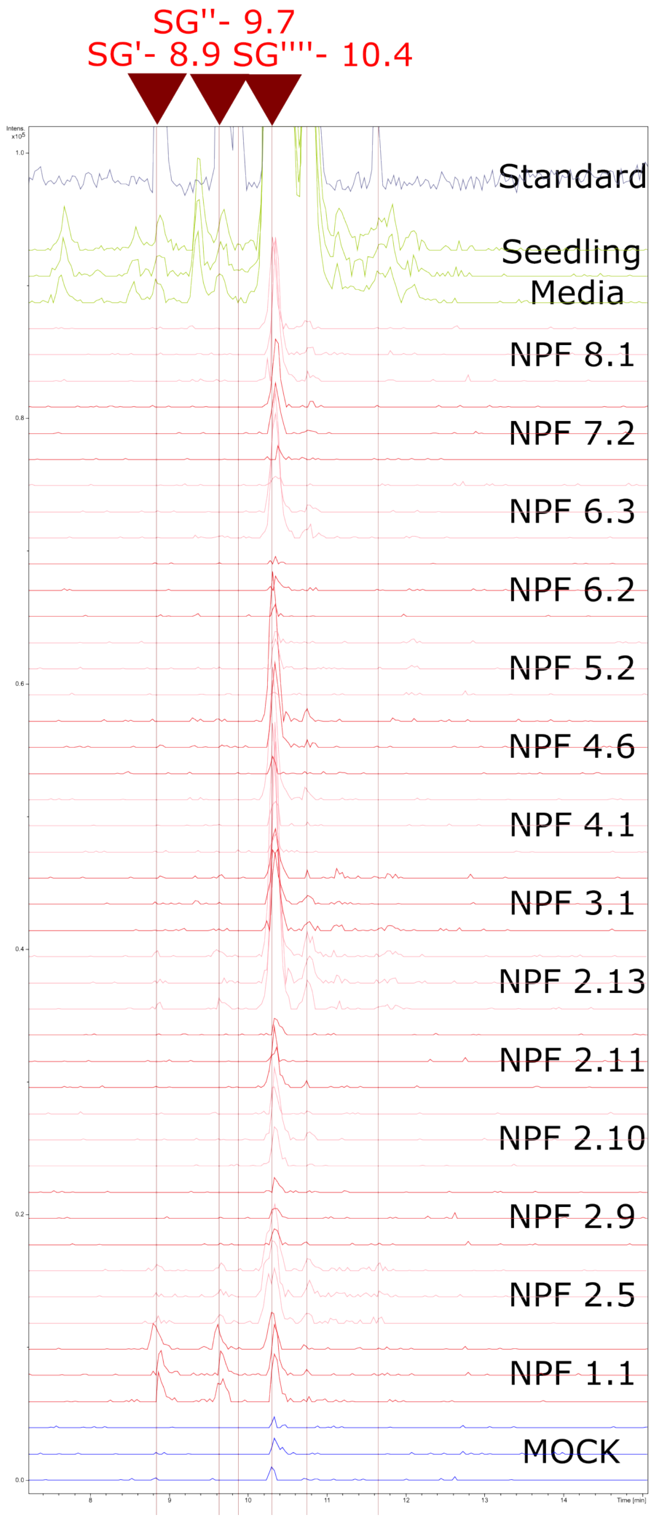

**Supplementary 24. Most transporters display accumulation levels of the most abundant isomer of sinapoyl glucose (SG), while NPF1.1 displays relatively high accumulation of two other isomers.** Presence of SG in oocytes exposed to 1:20 seedling Transporter-expressing oocytes in red and pink, alternatively. Control oocytes in blue. Media in green. 10 μM Chemical standard in black. Oocytes were assayed at pH 5 for 1 hour; replicates consisted of five oocytes each, and media samples consisted of 5μl each. Extracted ion chromatogram for SG (385.1140 +/- 0.005 m/z).

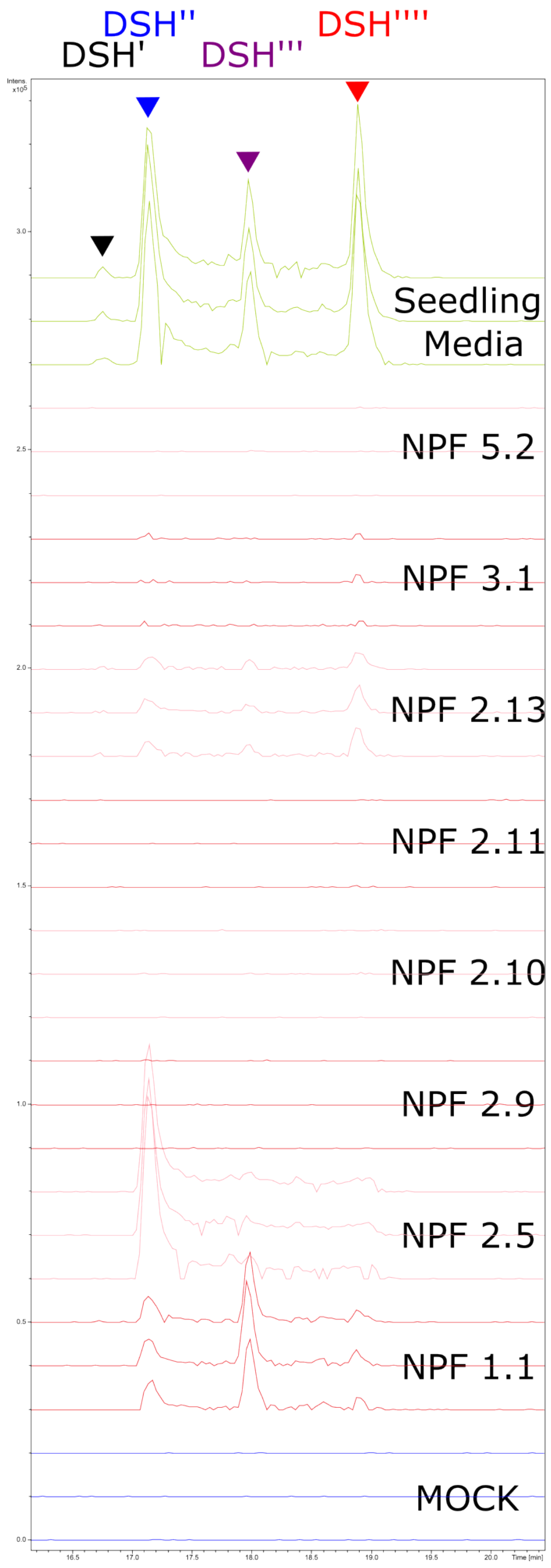

**Supplementary 25. NPF 1.1, 2.5, 2.13 and 3.1 display different stereospecificities towards isomers of the annotated disinapoyl Hexoses (DSH).** Presence of DSH isomers in oocytes exposed to 1:20 seedling media. Transporter-expressing oocytes in red and pink, alternatively. Control oocytes in blue. Media in green. Oocytes were assayed at pH 5 for 1 hour; replicates consisted of five oocytes each, and media samples consisted of 5µl each. Extracted ion chromatogram for DSH (591.1719 +/- 0.005 m/z).

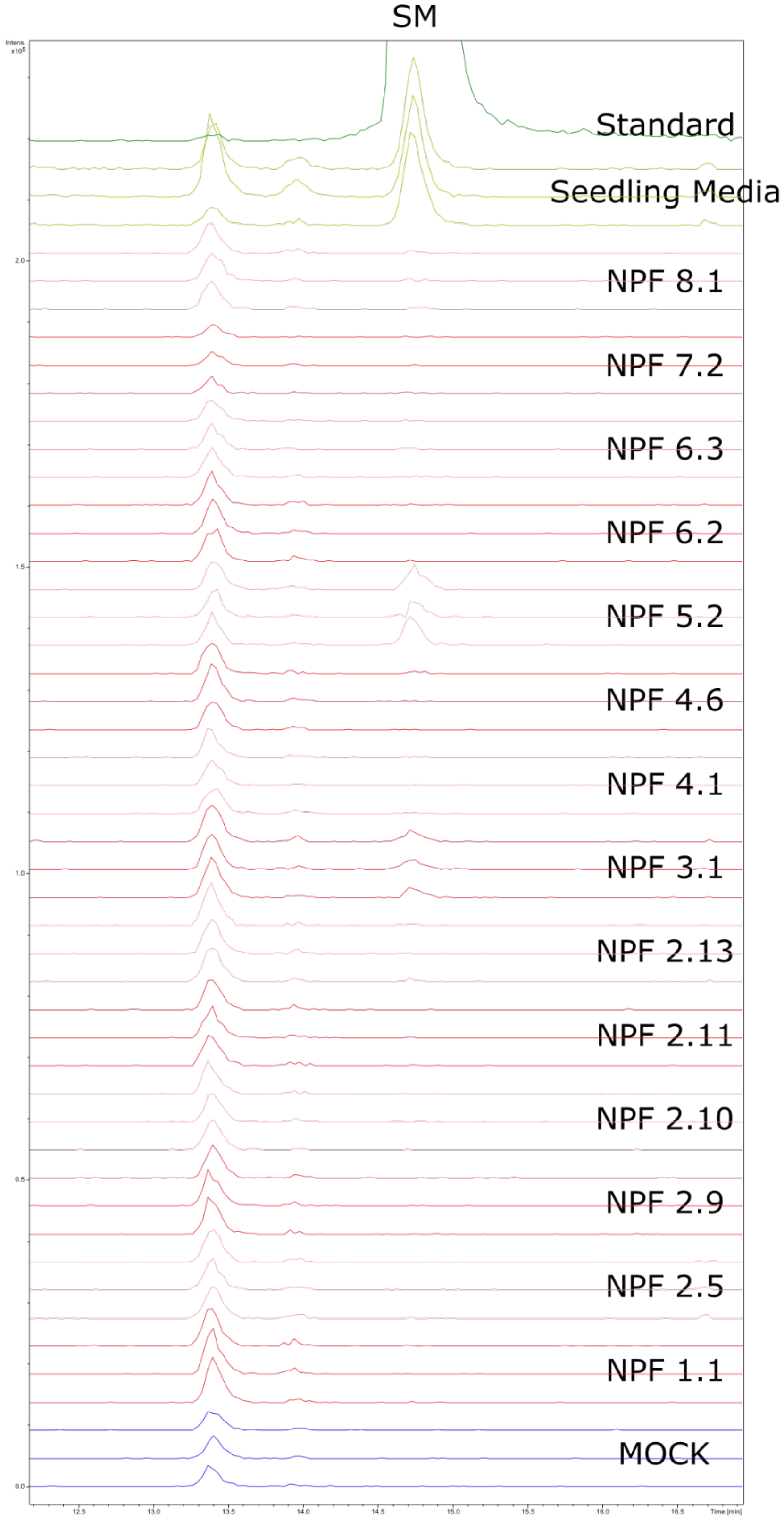

**Supplementary 26. NPF 5.2 imports the orphan substrate sinapoyl malate (SM).** Presence of SM in oocytes exposed to 1:20 seedling media. Transporter-expressing oocytes in red and pink, alternatively. Control oocytes in blue. Media in green. 10µM Chemical standard in black. Oocytes were assayed at pH 5 for 1 hour; replicates consisted of five oocytes each, and media samples consisted of 5µl each. Extracted ion chromatogram for the in-source fragment of SM corresponding to the sinapic acid in-source fragment (223.0612 +/- 0.005 m/z).

**Supplementary Table 1. Features accumulating in NPF1.1-expressing oocytes and present in the media.** FC: Fold change accumulation in transporter-expressing oocytes to mock oocytes. P.adjusted: p.value adjusted for False Discovery Rate according to the Benjamini-Hochberg method. Level: Annotation level of metabolic features according to the Metabolomics Standards Initiative (1: Molecular structure, confirmed by standards; 3: Possible structure; 4: Unknown feature). Pattern: Accumulation pattern of the feature across transporters or across transporters according to a chemical class.

| Feature | FC | p.ajusted | Annotation | Level | Formula | Pattern |
|---|---|---|---|---|---|---|
| 4219/421.0473mz/8.74min | 5.82 | 0.0017723 | Glucoerucin | 1 | C12H23NO9S3 | |
| 4264/247.0602mz/8.88min | 17.80 | 3.52E-07 | Sinapyl Glucose | 1 | C17H22O10 | |
| 4321/247.0602mz/9.66min | 24.35 | 6.13E-08 | Sinapyl Glucose | 1 | C17H22O10 | |
| 4387/385.1123mz/10.34min | 4.76 | 0.047592 | Sinapyl Glucose | 1 | C17H22O10 | |
| 4486/493.1048mz/11.06min | 4.56 | 0.042538 | Glucohirsutin | 3 | C16H31NO10S3 | |
| 4617/434.0600mz/11.69min | 4.00 | 0.020615 | Glucoberteroin | 3 | C13H25NO9S3 | |
| 4734/549.0834mz/12.76min | 3.72 | 0.042743 | | 4 | | Aliphatic glucosinolate |
| 4771/609.1455mz/13.02min | 4.18 | 0.0001633 | Quercetin-Rhamanoside-Hexoside | 3 | C27H30O16 | |
| 4868/563.0996mz/14.14min | 3.47 | 0.01711 | | 4 | | Aliphatic glucosinolate |
| 4872/593.1502mz/14.25min | 12.13 | 0.013862 | Keampferol-Rhamanoside-Hexoside | 3 | C27H30O15 | |
| 4885/593.1503mz/14.33min | 12.13 | 0.013862 | Quercetin-diRhamnoside | 3 | C27H30O15 | |
| 4928/448.0760mz/14.99min | 7.57 | 0.0001178 | Glucolesquerellin | 3 | C14H27NO9S3 | |
| 4953/480.0622mz/15.15min | 34.29 | 9.13E-06 | Glucomalcomiin | 3 | C17H23NO11S2 | |
| 5005/577.1553mz/15.59min | 13.83 | 1.67E-07 | Keaempferol-diRhamnoside | 3 | C27H30O14 | |
| 5052/447.0916mz/16.33min | 9.88 | 3.34E-06 | Quercitrin/Astragalin | 1 | C21H20O11 | |
| 5091/591.1706mz/17.15min | 48.54 | 3.36E-08 | Disinapoyl glucose | 3 | C28H28O14 | |
| 5144/494.0786mz/17.23min | 10.09 | 0.01321 | 4-benzoyloxybutyl glucosinolate | 3 | C18H25NO11S2 | |
| 5184/274.0709mz/17.46min | 5.58 | 7.17E-05 | | 4 | | Endogenous to oocyte, higher in NPF2.5 |
| 5226/462.0919mz/17.84min | 28.43 | 0.0048813 | 7-(methylthio)heptylglucosinolate | 3 | C15H29NO9S3 | |
| 5250/591.1707mz/17.97min | 20.48 | 8.05E-08 | Disinapoyl glucose | 3 | C28H32O14 | |
| 5264/698.1612mz/18.13min | 4.42 | 1.66E-05 | | 4 | | Benzoyl glycoside/NPF1.1, 2.13 and 3.1 |
| 5355/591.1707mz/18.90min | 14.62 | 8.31E-08 | Disinapoyl glucose | 3 | C28H32O14 | |
| 5367/626.1030mz/18.99min | 3.39 | 2.53E-05 | | 4 | | Benzoyl glycoside/NPF1.1, 2.13 and 3.1 |
| 5434/476.1075mz/19.45min | 49.15 | 0.0041652 | 8-(Methythio)octylglucosinolate | 3 | C16H31NO9S3 | |
| 5523/524.0706mz/20.01min | 40.72 | 6.91E-08 | 6′-O-benzoyloxy-glucoerucin | 3 | C19H27NO10S3 | |
| 5524/686.1215mz/20.02min | 2.69 | 2.83E-05 | 6'bz3sin | 3 | C28H33NO15S2 | |
| 5598/700.1370mz/20.55min | 45.40 | 3.36E-08 | 6'bz4sin | 3 | C29H35NO15S2 | |
| 5693/700.1361mz/21.01min | 3.28 | 0.0077392 | 6'bz4sin | 3 | C29H35NO15S2 | |
| 5702/668.1498mz/21.05min | 6.87 | 3.79E-06 | | 4 | | |
| 5855/598.1045mz/21.92min | 10.87 | 3.79E-06 | 6′-O-benzoyloxy-4-benzoyloxybutylglucosinolate | 3 | C25H29NO12S2 | |

**Supplementary Table 2. Features accumulating in NPF2.5-expressing oocytes and present in the media.** FC: Fold change accumulation in transporter-expressing oocytes to mock oocytes. P.adjusted: p.value adjusted for False Discovery Rate according to the Benjamini-Hochberg method. Level: Annotation level of metabolic features according to the Metabolomics Standards Initiative (1: Molecular structure, confirmed by standards; 3: Possible structure; 4: Unknown feature). Pattern: Accumulation pattern of the feature across transporters or across transporters according to a chemical class.

| Feature | FC | p.ajusted | Annotation | Level | Formula | Pattern |
|---|---|---|---|---|---|---|
| 4136/179.0704mz/8.10min | 175.6 | 1.26E-10 | Coniferin | 1 | C16H22O8 | |
| 4387/385.1123mz/10.34min | 14.2 | 0.0089941 | Sinapyl Glucose | 1 | C17H22O10 | |
| 5052/447.0916mz/16.33min | 31.6 | 7.25E-08 | Quercitrin | 1 | C21H20O11 | |
| 5063/477.1024mz/16.58min | 5.9 | 1.81E-07 | Isorhamnetin Glucoside | 3 | C22H22O12 | |
| 5091/591.1706mz/17.15min | 348.1 | 1.73E-09 | Disinapoyl glucose | 3 | C28H28O14 | |
| 5184/274.0709mz/17.46min | 47.5 | 1.03E-08 | | 4 | | Endogenous to oocyte, higher in NPF2.5 |
| 5189/431.0963mz/17.52min | 12.0 | 2.22E-07 | Afzelin | 1 | C21H20O10 | |
| 5203/461.1066mz/17.70min | 19.9 | 1.03E-08 | isorhamnetin rhamnoside | 3 | C22H22O11 | |
| 5250/591.1707mz/17.97min | 22.2 | 3.82E-08 | Disinapoyl glucose | 3 | C28H32O14 | |
| 5355/591.1707mz/18.90min | 54.2 | 5.64E-09 | Disinapoyl glucose | 3 | C28H32O14 | |

**Supplementary Table 3. Features accumulating in NPF2.9-expressing oocytes and present in the media.** FC: Fold change accumulation in transporter-expressing oocytes to mock oocytes. P.adjusted: p.value adjusted for False Discovery Rate according to the Benjamini-Hochberg method. Level: Annotation level of metabolic features according to the Metabolomics Standards Initiative (1: Molecular structure, confirmed by standards; 3: Possible structure; 4: Unknown feature). Pattern: Accumulation pattern of the feature across transporters or across transporters according to a chemical class.

| Feature | FC | p.ajusted | Annotation | Level | Formula | Pattern |
|---|---|---|---|---|---|---|
| 3988/463.0469mz/6.55min | 29.123 | 1.81E-09 | 4-Hydroxyglucobrassicin | 3 | C16H20N2O10S2 | |
| 4164/495.0738mz/8.45min | 13.068 | 1.86E-07 | | 4 | | Aliphatic Glucosinolate |
| 4188/478.0866mz/8.60min | 6.5555 | 0.023737 | Glucoibarin | 3 | C15H29NO10S3 | |
| 4231/420.0447mz/8.74min | 9.765 | 0.019321 | Glucoerucin | 1 | C12H23NO9S3 | |
| 4358/447.0522mz/10.20min | 2464.1 | 1.66E-13 | Glucosbrassicin | 1 | C16H20N2O9S2 | |
| 4513/492.1026mz/11.07min | 5.9717 | 0.021362 | Glucohirsutin | 3 | C16H31NO10S3 | |
| 4617/434.0600mz/11.69min | 58.875 | 7.95E-11 | Glucoberteroin | 3 | C13H25NO9S3 | |
| 4747/477.0624mz/12.81min | 108.01 | 7.01E-13 | 4-methoxyglucobrassicin | 1 | C17H22N2O10S2 | |
| 4844/510.0728mz/13.67min | 11.599 | 2.85E-09 | | | C15H29NO12S3 | Aliphatic Glucosinolate |
| 4928/448.0760mz/14.99min | 20.365 | 1.19E-07 | Glucolesquerellin | 3 | C14H27NO9S3 | |
| 4953/480.0622mz/15.15min | 48.467 | 3.87E-06 | Glucomalcomiin | 3 | C17H23NO11S2 | |
| 4969/477.0625mz/15.39min | 87.058 | 7.01E-13 | Neoglucobrassicin | 1 | C17H22N2O10S2 | |
| 5105/596.1100mz/17.20min | 11.517 | 3.68E-08 | 4-sinapoyloxybutylglucosinolate | 3 | C22H31NO14S2 | |
| 5144/494.0786mz/17.23min | 196.76 | 0.00015808 | 4-benzoyloxybutyl glucosinolate | 3 | C18H25NO11S2 | |
| 5184/274.0709mz/17.46min | 12.064 | 2.68E-07 | | 4 | | Endogenous to oocyte, higher in NPF2.5 |
| 5226/462.0919mz/17.84min | 32.122 | 0.0030519 | 7-(methylthio)heptylglucosinolate | 3 | C15H29NO9S3 | |
| 5301/508.0938mz/18.57min | 2.2411 | 5.64E-07 | 5-benzoyloxypentil glucosinolate | 3 | C19H27NO11S2 | |
| 5434/476.1075mz/19.45min | 43.336 | 0.0034924 | (Methythio)octylglucosinolate | 3 | C16H31NO9S3 | |

**Supplementary Table 4. Features accumulating in NPF2.10-expressing oocytes and present in the media.** FC: Fold change accumulation in transporter-expressing oocytes to mock oocytes. P.adjusted: p.value adjusted for False Discovery Rate according to the Benjamini-Hochberg method. Level: Annotation level of metabolic features according to the Metabolomics Standards Initiative (1: Molecular structure, confirmed by standards; 3: Possible structure; 4: Unknown feature). Pattern: Accumulation pattern of the feature across transporters or across transporters according to a chemical class.

| Feature | FC | p.ajusted | Annotation | Level | Formula | Pattern |
|---|---|---|---|---|---|---|
| 3021/436.0410mz/2.24min | 1278.4 | 2.10E-13 | Glucoraphanin | 1 | C12H23NO10S3 | |
| 3349/358.0256mz/2.41min | 4.3 | 2.91E-09 | Sinigrin | 3 | | |
| 3743/450.0545mz/4.12min | 244.5 | 1.42E-13 | Glucosalyssin | 3 | C13H25NO10S3 | |
| 3950/464.0709mz/6.32min | 484.0 | 1.31E-13 | Glucohesperin | 3 | C14H27NO10S3 | |
| 3962/481.0575mz/6.48min | 66.3 | 2.19E-09 | | 4 | | Aliphatic Glucosinolate |
| 3988/463.0469mz/6.55min | 5.6 | 2.28E-05 | 4-Hydroxyglucobrassicin | 3 | C16H20N2O10S2 | |
| 4164/495.0738mz/8.45min | 289.2 | 8.68E-05 | | 4 | | Aliphatic Glucosinolate |
| 4177/418.0824mz/8.58min | 6.3 | 2.61E-08 | n-Hydroxyhexyl glucosinolate | 3 | C13H25NO10S2 | |
| 4180/495.0738mz/8.45min | 404.8 | 9.56E-13 | | 4 | | Aliphatic Glucosinolate |
| 4188/478.0866mz/8.60min | 655.3 | 2.22E-05 | Glucoibarin | 3 | C15H29NO10S3 | |
| 4231/420.0447mz/8.74min | 811.8 | 4.09E-05 | Glucoerucin | 1 | C12H23NO9S3 | |
| 4330/508.0971mz/9.82min | 46.8 | 1.46E-11 | 8-(Methylsulfonyl)octy glucosinolate | 3 | C16H31NO11S3 | |
| 4358/447.0522mz/10.20min | 1763.1 | 7.01E-14 | Glucosbrassicin | 1 | C16H20N2O9S2 | |
| 4387/385.1123mz/10.34min | 6.1 | 0.019307 | Sinapyl Glucose | 1 | C17H22O10 | |
| 4513/492.1026mz/11.07min | 607.0 | 1.24E-05 | Glucohirsutin | 3 | C16H31NO10S3 | |
| 4611/422.0567mz/11.68min | 6.5 | 3.25E-10 | Gluconasturtiin | 3 | C15H21NO9S2 | |
| 4617/434.0600mz/11.69min | 2102.7 | 7.01E-14 | Glucoberteroin | 3 | C13H25NO9S3 | |
| 4667/484.0572mz/12.10min | 7.4 | 4.71E-09 | | 4 | | Aliphatic Glucosinolate |
| 4673/508.0972mz/12.17min | 5.5 | 1.60E-10 | 8-(Methylsulfonyl)octy glucosinolate | 3 | C16H31NO11S3 | |
| 4734/549.0834mz/12.76min | 192.0 | 4.21E-12 | | 4 | | Aliphatic Glucosinolate |
| 4747/477.0624mz/12.81min | 14.4 | 6.75E-11 | 4-methoxyglucobrassicin | 1 | C17H22N2O10S2 | |
| 4788/549.0839mz/13.11min | 228.6 | 7.34E-13 | | 4 | | Aliphatic Glucosinolate |
| 4845/402.0882mz/13.68min | 117.3 | 1.00E-12 | n-hexyl glucosinolate | 3 | C13H25NO9S2 | |
| 4868/563.0996mz/14.14min | 628.6 | 7.44E-14 | | 4 | | Aliphatic Glucosinolate |
| 4928/448.0760mz/14.99min | 1908.4 | 7.01E-14 | Glucolesquerellin | 3 | C14H27NO9S3 | |
| 4953/480.0622mz/15.15min | 3701.5 | 6.63E-09 | Glucomalcomiin | 3 | C17H23NO11S2 | |
| 4999/477.0625mz/15.39min | 311.6 | 1.01E-13 | Neoglucobrassicin | 1 | C17H22N2O10S2 | |
| 5018/582.0947mz/15.85min | 11.8 | 1.27E-10 | 3-sinapoyloxypropylglucosinolate | 3 | C21H29NO14S2 | |
| 5052/447.0916mz/16.33min | 22.9 | 1.11E-11 | Quercitrin | 1 | C21H20O11 | |
| 5105/596.1100mz/17.20min | 426.9 | 1.31E-13 | 4-sinapoyloxybutylglucosinolate | 3 | C22H31NO14S2 | |
| 5144/494.0786mz/17.23min | 1568.4 | 7.33E-06 | 4-benzoyloxybutyl glucosinolate | 3 | C18H25NO11S2 | |
| 5170/416.1033mz/17.31min | 46.6 | 1.11E-10 | n-Heptyl glucosinolate | 3 | C14H27NO9S2 | |
| 5184/274.0709mz/17.46min | 7.4 | 0.002825 | | 4 | | Endogenous to oocyte, higher in NPF2.5 |
| 5203/461.1066mz/17.70min | 2.9 | 2.81E-05 | isorhamnetin rhamnoside | 3 | C22H22O11 | |
| 5226/462.0919mz/17.84min | 2290.4 | 1.40E-05 | 7-(methylthio)heptylglucosinolate | 3 | C15H29NO9S3 | |
| 5301/508.0938mz/18.57min | 27.1 | 2.29E-12 | 5-benzoyloxypentil glucosinolate | 3 | C19H27NO11S2 | |
| 5343/492.1028mz/18.79min | 22.1 | 1.11E-11 | 8-Methyl-thio-Hydroxyoctilglucosinolate | 3 | C16H31NO10S3 | |
| 5434/476.1075mz/19.45min | 2350.5 | 2.73E-05 | (Methythio)octylglucosinolate | 3 | C16H31NO9S3 | |

**Supplementary Table 5. Features accumulating in NPF2.11-expressing oocytes and present in the media.** FC: Fold change accumulation in transporter-expressing oocytes to mock oocytes. P.adjusted: p.value adjusted for False Discovery Rate according to the Benjamini-Hochberg method. Level: Annotation level of metabolic features according to the Metabolomics Standards Initiative (1: Molecular structure, confirmed by standards; 3: Possible structure; 4: Unknown feature). Pattern: Accumulation pattern of the feature across transporters or across transporters according to a chemical class.

| Feature | FC | p.ajusted | Annotation | Level | Formula | Pattern |
|---|---|---|---|---|---|---|
| 3021/436.0410mz/2.24min | 1473.9 | 3.12E-12 | Glucoraphanin | 1 | C12H23NO10S3 | |
| 3349/358.0256mz/2.41min | 2.2 | 1.62E-06 | Sinigrin | 3 | C10H17NO9S2 | |
| 3743/450.0545mz/4.12min | 356.8 | 3.41E-10 | Glucosalyssin | 3 | C13H25NO10S3 | |
| 3950/464.0709mz/6.32min | 461.3 | 4.78E-11 | Glucohesperin | 3 | C14H27NO10S3 | |
| 3962/481.0575mz/6.48min | 67.4 | 1.45E-10 | | 4 | | Aliphatic pattern |
| 3988/463.0469mz/6.55min | 5.5 | 1.83E-09 | 4-Hydroxyglucobrassicin | 3 | C16H20N2O10S2 | |
| 4164/495.0738mz/8.45min | 246.8 | 9.26E-05 | | 4 | | Aliphatic Glucosinolate |
| 4177/418.0824mz/8.58min | 7.8 | 9.12E-09 | n-Hydroxyhexyl glucosinolate | 3 | C13H25NO10S2 | |
| 4180/495.0738mz/8.45min | 278.2 | 0.001094 | | 4 | | Aliphatic Glucosinolate |
| 4188/478.0866mz/8.60min | 643.9 | 2.42E-05 | Glucoibarin | 3 | C15H29NO10S3 | |
| 4231/420.0447mz/8.74min | 942.8 | 3.74E-05 | Glucoerucin | 1 | C12H23NO9S3 | |
| 4330/508.0971mz/9.82min | 94.5 | 1.48E-10 | 8-(Methylsulfonyl)octy glucosinolate | 3 | C16H31NO11S3 | Aliphatic Glucosinolate |
| 4358/447.0522mz/10.20min | 1579.9 | 2.35E-13 | Glucosbrassicin | 1 | C16H20N2O9S2 | |
| 4513/492.1026mz/11.07min | 719.7 | 1.09E-05 | Glucohirsutin | 3 | C16H31NO10S3 | |
| 4611/422.0567mz/11.68min | 8.4 | 1.24E-10 | Gluconasturtiin | 3 | C15H21NO9S2 | |
| 4617/434.0600mz/11.69min | 2987.3 | 2.35E-13 | Glucoberteroin | 3 | C13H25NO9S3 | |
| 4667/484.0572mz/12.10min | 6.7 | 4.41E-10 | | 4 | | Aliphatic Glucosinolate |
| 4673/508.0972mz/12.17min | 6.3 | 9.40E-10 | 8-(Methylsulfonyl)octy glucosinolate | 3 | C16H31NO11S3 | |
| 4734/549.0834mz/12.76min | 197.8 | 7.49E-12 | | 4 | | Aliphatic Glucosinolate |
| 4747/477.0624mz/12.81min | 10.8 | 1.73E-10 | 4-methoxyglucobrassicin | 1 | C17H22N2O10S2 | |
| 4788/549.0839mz/13.11min | 218.0 | 1.66E-10 | | 4 | | Aliphattic profil |
| 4845/402.0882mz/13.68min | 133.1 | 6.29E-12 | n-hexyl glucosinolate | 3 | C13H25NO9S2 | |
| 4868/563.0996mz/14.14min | 499.1 | 7.58E-13 | | 4 | | Aliphatic Glucosinolate |
| 4928/448.0760mz/14.99min | 2494.0 | 2.35E-13 | Glucolesquerellin | 3 | C14H27NO9S3 | |
| 4953/480.0622mz/15.15min | 3578.9 | 7.71E-09 | Glucomalcomiin | 3 | C17H23NO11S2 | |
| 4969/477.0625mz/15.39min | 41.8 | 1.02E-11 | Neoglucobrassicin | 1 | C17H22N2O10S2 | |
| 5018/582.0947mz/15.85min | 2.7 | 2.71E-07 | 3-sinapoyloxypropylglucosinolate | 0 | C21H29NO14S2 | |
| 5052/447.0916mz/16.33min | 21.2 | 8.56E-08 | Quercitrin | 1 | C21H20O11 | |
| 5105/596.1100mz/17.20min | 150.1 | 5.62E-13 | 4-sinapoyloxybutylglucosinolate | 3 | C22H31NO14S2 | |
| 5144/494.0786mz/17.23min | 1493.8 | 7.88E-06 | 4-benzoyloxybutyl glucosinolate | 3 | C18H25NO11S2 | |
| 5170/416.1033mz/17.31min | 63.0 | 1.41E-11 | n-Heptyl glucosinolate | 3 | C14H27NO9S2 | |
| 5184/274.0709mz/17.46min | 21.9 | 2.49E-10 | | 4 | | Endogenous to oocyte, higher in NPF2.5 |
| 5189/431.0963mz/17.52min | 2.8 | 0.043307 | Afzelin | 1 | C21H20O10 | |
| 5203/461.1066mz/17.70min | 4.8 | 0.00022 | isorhamnetin rhamnoside | 3 | C22H22O11 | |
| 5226/462.0919mz/17.84min | 2552.3 | 1.33E-05 | 7-(methylthio)heptylglucosinolate | 3 | C15H29NO9S3 | |
| 5301/508.0938mz/18.57min | 24.3 | 6.89E-12 | 5-benzoyloxypentil glucosinolate | 3 | C19H27NO11S2 | |
| 5343/492.1028mz/18.79min | 38.6 | 3.50E-11 | 8-Methyl-thio-Hydroxyoctilglucosinolate | 3 | C16H31NO10S3 | |
| 5434/476.1075mz/19.45min | 2912.3 | 2.43E-05 | (Methythio)octylglucosinolate | 3 | C16H31NO9S3 | |
| 5523/524.0706mz/20.01min | 52.9 | 5.17E-08 | 6bz4mtb | 3 | C19H27NO10S3 | |

| | | | | | |
|---|---|---|---|---|---|
| 5855/598.1045mz/21.92min | 16.9 | 2.52E-05 | 4-(benzoyloxy)-6′-O-benzoyloxybutylglucosinolate | 3 | C25H29NO12S2 |

**Supplementary Table 6. Features accumulating in NPF2.13-expressing oocytes and present in the media.** FC: Fold change accumulation in transporter-expressing oocytes to mock oocytes. P.adjusted: p.value adjusted for False Discovery Rate according to the Benjamini-Hochberg method. Level: Annotation level of metabolic features according to the Metabolomics Standards Initiative (1: Molecular structure, confirmed by standards; 3: Possible structure; 4: Unknown feature). Pattern: Accumulation pattern of the feature across transporters or across transporters according to a chemical class.

| RT | FC | p.ajusted | Annotation | Level | Formula | Pattern |
|---|---|---|---|---|---|---|
| 3021/436.0410mz/2.24min | 5.0 | 0.045024 | Glucoraphanin | 1 | C12H23NO10S3 | |
| 3950/464.0709mz/6.32min | 7.6 | 0.001186 | Glucohesperin | 3 | C14H27NO10S3 | |
| 4164/495.0738mz/8.45min | 12.9 | 4.33E-07 | | 4 | | Aliphatic glucosinolate |
| 4188/478.0866mz/8.60min | 22.6 | 0.002344 | Glucoibarin | 3 | C15H29NO10S3 | |
| 4231/420.0447mz/8.74min | 20.2 | 0.004478 | Glucoerucin | 1 | C12H23NO9S3 | |
| 4358/447.0522mz/10.20min | 26.5 | 1.74E-07 | Glucosbrassicin | 1 | C16H20N2O9S2 | |
| 4387/385.1123mz/10.34min | 41.7 | 0.000573 | Sinapyl Glucose | 1 | C17H22O10 | |
| 4451/385.1126mz/10.77min | 11.7 | 4.00E-06 | Sinapyl Glucose | 1 | C17H22O10 | |
| 4513/492.1026mz/11.07min | 17.9 | 0.002036 | Glucohirsutin | 3 | C16H31NO10S3 | |
| 4617/434.0600mz/11.69min | 102.5 | 5.64E-10 | Glucoberteroin | 3 | C13H25NO9S3 | |
| 4771/609.1455mz/13.02min | 75.1 | 2.91E-09 | Quercetin-Rhmanoside-Hexoside | 3 | C27H30O16 | |
| 4868/563.0996mz/14.14min | 34.6 | 4.03E-09 | | 4 | | Aliphatic glucosinolate |
| 4872/593.1502mz/14.25min | 14.7 | 0.005443 | Keampferol-Rhmanoside-Hexoside | 3 | C27H30O15 | |
| 4885/593.1503mz/14.33min | 71.8 | 0.000573 | Quercetin-diRhamnoside | 3 | C27H30O15 | |
| 4899/623.1606mz/14.51min | 7.7 | 2.56E-05 | Isorhamnetin-Rhamnoside-Hexoside | 3 | C28H32O16 | |
| 4919/463.0866mz/14.81min | 3.4 | 0.000573 | | 1 | C21H20O12 | |
| 4928/448.0760mz/14.99min | 164.0 | 1.75E-10 | Glucolesquerellin | 3 | C14H27NO9S3 | Aliphatic glucosinolate |
| 4940/448.0759mz/14.99min | 161.6 | 9.99E-11 | Glucolesquerellin | 3 | C14H27NO9S3 | Aliphatic glucosinolate |
| 4953/480.0622mz/15.15min | 540.3 | 1.70E-07 | Glucomalcomiin | 3 | C17H23NO11S2 | |
| 4969/477.0625mz/15.39min | 2.5 | 1.56E-05 | Neoglucobrassicin | 1 | C17H22N2O10S2 | |
| 5005/577.1553mz/15.59min | 109.2 | 5.12E-10 | Keaempferol-diRhamnoside | | | |
| 5019/642.0988mz/15.84min | 2.1 | 0.04276 | | 4 | | NPF3.1 and NPF 2.13 |
| 5023/607.1660mz/15.92min | 15.8 | 2.31E-07 | Isorhamnetin-diRhmanoside | 3 | C28H32O15 | |
| 5052/447.0916mz/16.33min | 369.3 | 9.99E-11 | Quercitrin | 1 | C21H20O11 | |
| 5063/477.1024mz/16.58min | 3.3 | 0.002016 | Isorhamnetin Glucoside | 3 | C22H22O12 | |
| 5071/657.1807mz/16.78min | 2.7 | 0.001409 | | 4 | | Only NF 2.13 |
| 5091/591.1706mz/17.15min | 24.9 | 3.80E-08 | Disinapoyl glucose | 3 | C28H28O14 | |
| 5105/596.1100mz/17.20min | 22.6 | 2.12E-08 | 4-sinapoyloxybutylglucosinolate | 2 | C22H31NO14S2 | |
| 5144/494.0786mz/17.23min | 233.5 | 9.16E-05 | 4-benzoyloxybutyl glucosinolate | 2 | C18H25NO11S2 | |
| 5178/591.1708mz/17.16min | 20.9 | 2.82E-07 | Disinapoyl glucose | 3 | C28H28O14 | |
| 5184/274.0709mz/17.46min | 11.1 | 3.28E-06 | | 4 | | Endogenous to oocyte, higher in NPF2.5 |
| 5189/431.0963mz/17.52min | 115.2 | 2.67E-10 | Afzelin | 1 | C21H20O10 | |
| 5203/461.1066mz/17.70min | 46.1 | 1.76E-09 | isorhamnetin rhamnoside | 3 | C22H22O11 | |
| 5226/462.0919mz/17.84min | 283.0 | 0.000176 | 7-(methylthio)heptylglucosinolate | 3 | C15H29NO9S3 | |
| 5250/591.1707mz/17.97min | 2.5 | 0.000297 | Disinapoyl glucose | 3 | C28H32O14 | |
| 5264/698.1612mz/18.13min | 4.6 | 5.78E-05 | | 4 | | Benzoyl glycoside/NPF1.1, 2.13 and 3.1 |
| 5301/508.0938mz/18.57min | 5.4 | 1.08E-09 | 5-benzoyloxypentil glucosinolate | 3 | C19H27NO11S2 | |
| 5343/492.1028mz/18.79min | 3.5 | 0.000299 | 8-Methyl-thio-Hydroxyoctilglucosinolate | 3 | C16H31NO10S3 | |
| 5355/591.1707mz/18.90min | 29.6 | 8.63E-07 | Disinapoyl glucose | 3 | C28H32O14 | |

| | | | | | | |
|---|---|---|---|---|---|---|
| 5367/626.1030mz/18.99min | 14.9 | 1.44E-06 | | 4 | | Benzoyl glycoside/NPF1.1, 2.13 and 3.1 |
| 5368/596.1287mz/19.00min | 7.1 | 0.010107 | | 4 | | 2.11, 2.13 and 3.1 |
| 5413/626.1036mz/19.38min | 9.7 | 1.70E-06 | | 4 | | Benzoyl glycoside/NPF1.1, 2.13 and 3.1 |
| 5434/476.1075mz/19.45min | 375.3 | 0.000243 | (Methythio)octylglucosinolate | 3 | C16H31NO9S3 | |
| 5523/524.0706mz/20.01min | 42.5 | 1.74E-07 | 6′-O-benzoyloxy-glucoerucin | 3 | C19H27NO10S3 | |
| 5524/686.1215mz/20.02min | 8.3 | 2.72E-07 | NOT 6bzo4mtb | 4 | | Benzoyl glycoside/NPF1.1, 2.13 and 3.1 |
| 5545/524.0705mz/20.02min | 35.1 | 0.004125 | 6′-O-benzoyloxy-glucoerucin | 3 | C19H27NO10S3 | |
| 5598/700.1370mz/20.55min | 105.9 | 1.80E-09 | 6'bz4sin | 3 | C29H35NO15S2 | |
| 5693/700.1361mz/21.01min | 17.1 | 6.29E-08 | 6'bz4sin | 3 | C29H35NO15S2 | |

**Supplementary Table 7. Features accumulating in NPF3.1-expressing oocytes and present in the media.** FC: Fold change accumulation in transporter-expressing oocytes to mock oocytes. P.adjusted: p.value adjusted for False Discovery Rate according to the Benjamini-Hochberg method. Level: Annotation level of metabolic features according to the Metabolomics Standards Initiative (1: Molecular structure, confirmed by standards; 3: Possible structure; 4: Unknown feature). Pattern: Accumulation pattern of the feature across transporters or across transporters according to a chemical class.

| Feature | FC | p.ajusted | Annotation | Level | Formula | Pattern |
|---|---|---|---|---|---|---|
| 3021/436.0410mz/2.24min | 39.3 | 0.00015155 | Glucoraphanin | 1 | C12H23NO10S3 | |
| 3743/450.0545mz/4.12min | 6.8 | 0.019635 | Glucosalyssin | 3 | C13H25NO10S3 | |
| 3950/464.0709mz/6.32min | 21.5 | 8.13E-05 | Glucohesperin | 3 | C14H27NO10S3 | |
| 4164/495.0738mz/8.45min | 32.1 | 2.15E-07 | | 4 | | Aliphatic Glucosinolate |
| 4188/478.0866mz/8.60min | 36.1 | 0.00016266 | Glucoibarin | 3 | C15H29NO10S3 | |
| 4231/420.0447mz/8.74min | 48.0 | 4.47E-05 | Glucoerucin | 1 | C12H23NO9S3 | |
| 4358/447.0522mz/10.20min | 168.6 | 3.29E-08 | Glucosbrassicin | 1 | C16H20N2O9S2 | |
| 4387/385.1123mz/10.34min | 10.1 | 0.004689 | Sinapyl Glucose | 1 | C17H22O10 | |
| 4513/492.1026mz/11.07min | 34.9 | 0.00013858 | Glucohirsutin | 3 | C16H31NO10S3 | |
| 4617/434.0600mz/11.69min | 134.1 | 9.67E-08 | Glucoberteroin | 3 | C13H25NO9S3 | |
| 4734/549.0834mz/12.76min | 16.7 | 0.00099383 | | 4 | | Aliphatic Glucosinolate |
| 4747/477.0624mz/12.81min | 2.9 | 0.0001466 | 4-methoxyglucobrassicin | 1 | C17H22N2O10S2 | |
| 4771/609.1455mz/13.02min | 19.2 | 0.00013465 | Quercetin-Rhamnoside-Hexoside | 3 | C27H30O16 | |
| 4788/549.0839mz/13.11min | 15.9 | 7.22E-06 | | 4 | | Aliphatic Glucosinolate |
| 4844/510.0728mz/13.67min | 14.7 | 1.32E-06 | | 4 | | Aliphatic Glucosinolate |
| 4868/563.0996mz/14.14min | 124.2 | 1.43E-07 | | 4 | | Aliphatic Glucosinolate |
| 4872/593.1502mz/14.25min | 6.5 | 0.024594 | Keampferol-Rhamnoside-Hexoside | 3 | C27H30O15 | |
| 4885/593.1503mz/14.33min | 10.1 | 0.0091581 | Quercetin-diRhamnoside | 3 | C27H30O15 | |
| 4889/565.0995mz/14.16min | 6.4 | 0.022361 | | 4 | | Aliphatic Glucosinolate |
| 4916/223.0601mz/14.73min | 2.4 | 1.87E-06 | Sinapoyl malate | 1 | C15H16O9 | |
| 4928/448.0760mz/14.99min | 172.3 | 6.68E-09 | Glucolesquerellin | 3 | C14H27NO9S3 | |
| 4953/480.0622mz/15.15min | 641.1 | 1.40E-08 | Glucomalcomiin | 3 | C17H23NO11S2 | |
| 4969/477.0625mz/15.39min | 5.2 | 7.51E-07 | Neoglucobrassicin | 1 | C17H22N2O10S2 | |
| 5005/577.1553mz/15.59min | 11.2 | 0.00039862 | Keaempferol-diRhamnoside | 3 | C27H30O14 | |
| 5018/582.0947mz/15.85min | 2.6 | 5.72E-05 | 3-sinapoyloxypropylglucosinolate | | C21H29NO14S2 | |
| 5019/642.0988mz/15.84min | 3.6 | 0.012765 | | 4 | | NPF 3.1 and NPF 2.13 |
| 5052/447.0916mz/16.33min | 76.3 | 1.29E-09 | Quercitrin | 1 | C21H20O11 | |
| 5105/596.1100mz/17.20min | 69.3 | 1.02E-06 | 4-sinapoyloxybutylglucosinolate | 3 | C22H31NO14S2 | |
| 5144/494.0786mz/17.23min | 221.1 | 4.28E-07 | 4-benzoyloxybutyl glucosinolate | 3 | C18H25NO11S2 | |
| 5184/274.0709mz/17.46min | 14.5 | 4.87E-05 | | 4 | | Endogenous to oocyte, higher in NPF2.5 |
| 5189/431.0963mz/17.52min | 22.9 | 2.52E-07 | Afzelin | 1 | C21H20O10 | |
| 5203/461.1066mz/17.70min | 7.0 | 1.73E-05 | isorhamnetin rhamnoside | 3 | C22H22O11 | |
| 5209/597.1274mz/17.75min | 6.0 | 0.0037688 | | 4 | | NPF 3.1 and NPF 2.13 |
| 5226/462.0919mz/17.84min | 253.9 | 4.40E-07 | 7-(methylthio)heptylglucosinolate | 3 | C15H29NO9S3 | |
| 5264/698.1612mz/18.13min | 3.3 | 0.0020606 | | 4 | | NPF 3.1, NPF 2.13 and NPF 1.1 (Not DSG) |
| 5301/508.0938mz/18.57min | 6.4 | 9.94E-06 | 5-benzoyloxypentil glucosinolate | 3 | C19H27NO11S2 | |
| 5343/492.1028mz/18.79min | 2.9 | 1.32E-06 | 8-Methyl-thio-Hydroxyoctilglucosinolate | 3 | C16H31NO10S3 | |
| 5355/591.1707mz/18.90min | 6.8 | 0.0025946 | Disinapoyl glucose | 3 | C28H32O14 | |
| 5364/492.1029mz/18.80min | 3.6 | 5.98E-07 | 8-Methylthio-hydroxyoctyl glucosinolate | 3 | C16H31NO10S3 | |

| | | | | | | |
|---|---|---|---|---|---|---|
| 5367/626.1030mz/18.99min | 7.6 | 0.031891 | | 4 | | NPF 3.1, NPF 2.13 and NPF 1.1 |
| 5413/626.1036mz/19.38min | 9.9 | 0.00012915 | | 4 | | NPF 3.1, NPF 2.13 and NPF 1.1 |
| 5434/476.1075mz/19.45min | 339.4 | 4.58E-07 | 8-(Methythio)octylglucosinolate | 3 | C16H31NO9S3 | |
| 5523/524.0706mz/20.01min | 14.5 | 0.047643 | 6′-O-benzoyloxy-glucoerucin | 3 | C19H27NO10S3 | |
| 5524/686.1215mz/20.02min | 4.4 | 0.011905 | | 4 | | NPF 3.1, NPF 2.13 and NPF 1.1 (Not 6′-O-benzoyloxy-glucoerucin ) |
| 5598/700.1370mz/20.55min | 59.3 | 0.0007104 | 6′-O-benzoyloxy-4-sinapoyloxybutylglucosinolate | 3 | C29H35NO15S2 | |
| 5693/700.1361mz/21.01min | 11.7 | 9.92E-06 | 6′-O-benzoyloxy-4-sinapoyloxybutylglucosinolate | 3 | C29H35NO15S2 | |

**Supplementary Table 8. Features accumulating in NPF5.2-expressing oocytes and present in the media.** FC: Fold change accumulation in transporter-expressing oocytes to mock oocytes. P.adjusted: p.value adjusted for False Discovery Rate according to the Benjamini-Hochberg method. Level: Annotation level of metabolic features according to the Metabolomics Standards Initiative (1: Molecular structure, confirmed by standards; 3: Possible structure; 4: Unknown feature). Pattern: Accumulation pattern of the feature across transporters or across transporters according to a chemical class.

| Feature | FC | p.ajusted | Annotation | Level | Formula | Pattern |
|---|---|---|---|---|---|---|
| 3763/303.0707mz/4.33min | 3.0913 | 0.003024 | | 4 | | Only NPF 5.2 |
| 4916/223.0601mz/14.73min | 7.6567 | 0.0001683 1 | Sinapyl malate | 1 | $C_{11}H_{12}O_5$ | |

**Supplementary Table 9: MRM transitions, fragmentor voltages, and collision energies (CE) used for LC-MS/MS detection of disinapoyl sucrose and sinapoyl malate**. Q1: Voltage setting for quadrupole 1. Q3: Voltage setting for quadrupole 3. QT: quantifier ion.

| Analyte | Retention time (min) | Q1 (m/z) | Q3 (m/z) | Fragmentor (V) | CE (V) |
|---|---|---|---|---|---|
| Disinapoyl sucrose (DSS) | 1.94 | 753.2 | 547.5 | 140 | 28 |
| | 1.94 | 753.2 | 205.2$^{QT}$ | 140 | 36 |
| | 1.94 | 753.2 | 190.4 | 140 | 68 |
| | 1.94 | 753.2 | 189.7 | 140 | 68 |
| | 1.94 | 753.2 | 174.8 | 140 | 92 |
| Sinapoyl malate (SM) | 0.95 | 339.1 | 223.0$^{QT}$ | 50 | 8 |
| | 0.95 | 339.1 | 93.0 | 50 | 68 |
| | 0.95 | 339.1 | 120.9 | 50 | 52 |
| | 0.95 | 339.1 | 148.9 | 50 | 36 |
| | 0.95 | 339.1 | 164.1 | 50 | 24 |
